# Mycorrhizal and endophytic fungi structure contrasting but interdependent assembly processes in forest below-ground symbiosis

**DOI:** 10.1101/2024.02.17.580831

**Authors:** Mikihito Noguchi, Hirokazu Toju

## Abstract

Interactions between plants and diverse root-associated fungi are essential drivers of forest ecosystem dynamics. The nature of the symbiosis in root systems is potentially dependent on multiple ecological factors/processes such as host/symbiont specificity, background soil microbiome structure, inter-root sharing/dispersal of symbionts, and fungus–fungus interactions within fine roots. Nonetheless, it has remained a major challenge to reveal the mechanisms by which those multiple factors/processes determine the assembly of mycorrhizal and endophytic fungal communities. Based on a framework of joint species distribution modeling, we here examined how root-associated fungal community structure was collectively formed through filtering by host plants, associations with background soil fungi, spatial autocorrelation, and symbiont–symbiont interactions. In our analysis targeting 1,615 root-tip samples collected in a cool-temperate forest dominated by ectomycorrhizal plants, statistical models including all the four ecological factors/processes best explained the fine-scale community structure of root-associated fungi. Meanwhile, among partial models including subsets of those ecological factors/processes, those including background soil microbiome structure and within-root fungus–fungus interactions showed the highest performance. When fine-root distributions of respective fugal species/taxa were examined, ectomycorrhizal fungi tended to show stronger associations with background soil community structure and stronger spatially-autocorrelated patterns than other fungal guilds. In contrast, the distributions of root-endophytic fungi were inferred to depend greatly on fungus–fungus interactions. A network statistical analysis further suggested that some endophytic fungi, such as those belonging to the ascomycete genera *Phialocephala* and *Leptodontidium*, were involved in webs of positive and negative interactions with other root-associated fungi. These results suggest that basic assembly rules can differ between mycorrhizal and endophytic fungi, both of which are major components of forest ecosystems. Consequently, knowledge of how multiple ecological factors/processes differentially drive the assembly of multiple fungal functional guilds is indispensable for comprehensively understanding the mechanisms by which terrestrial ecosystem dynamics are organized by plant–fungal symbiosis.

## INTRODUCTION

Interactions between plants and root-associated fungi are essential components of forest ecosystems (Bennett et al., 2017; Dickie et al., 2014; Kadowaki et al., 2018; Mcguire, 2007; Van der Putten et al., 2013). Among major groups of root-associated fungi, mycorrhizal fungi provide host plants with nitrogen and phosphorus absorbed from the soil and, in return, they receive carbon (sugar and lipids) fixed by their host plants through photosynthesis (Smith & Read, 2008). In addition, mycorrhizal fungi protect their host plants from pathogens as well as various environmental stress such as drought, heat, and soil salinization (Duchesne et al., 1988; Sylvia DM & Sinclair WA, 1983; Tedersoo & Bahram, 2019). Among these mycorrhizal fungi, ectomycorrhizal fungi commonly occur in temperate and boreal forest ecosystems, forming symbiotic relationships with dominant plant families, such as Fagaceae and Pinaceae (Smith & Read, 2008). Because of their strong impacts on host plants’ fitness, they are considered to drive positive plant–soil feedback, in which seedlings of dominant plants are benefitted by below-ground hyphal networks and buried spores of compatible mycorrhizal fungi (Nara, 2006; Pec et al., 2020). Therefore, ecological processes of mycorrhizal fungi in fine roots, such as processes driven by preference for host plants and spatially dynamic recruitment of symbionts from the soil, are keys to understand forest ecosystem dynamics.

While ectomycorrhizal fungi are known as important components of temperate and boreal forests, recent studies based on high throughput DNA sequencing have revealed an unexpected diversity and prevalence of endophytic fungi in plant root systems (Nilsson et al., 2019; Toju et al., 2018). Among these root-endophytic fungi, dark septate endophytes have been reported from more than 600 plant species across 100 families, forming symbiotic relationship regardless of plants’ mycorrhizal type (Jumpponen & Trappe, 1998; Terhonen et al., 2019). Intriguingly, some dark septate endophytes are known to enhance host plants’ growth as well as plants’ tolerance to environmental and pathogen stress (Mandyam & Jumpponen, 2005, 2014; Usuki & Narisawa, 2007). Thus, not only ectomycorrhizal fungi but also root-endophytic fungi possibly play pivotal roles in feedback between plant communities and below-ground biota in temperate and boreal forests (Hewitt et al., 2023; Schittko & Wurst, 2014). Nonetheless, the basic ecology of root-endophytic fungi in forest ecosystems have remained poorly explored. Although root-endophytic fungi have been detected from diverse plant taxa, the degree of each species’ host preference in real ecosystems has been poorly examined (but see Põlme et al., 2018). In addition, while ectomycorrhizal fungi are known to form intensive extraradical hyphal networks for transporting soil nutrients (Smith & Read., 2008), little is known about how endophytic fungi form spatially structured processes of below-ground ecosystems. Therefore, towards comprehensive understanding of plant–soil feedback, it is essential to compare host preference levels and spatially structured ecological processes among major fungal guilds.

In addition to the ecological properties of respective fungi, facilitative and antagonistic interactions between fungal species are expected to play important roles in the community processes of root-associated fungi (Maherali & Klironomos., 2007, Pickles et al., 2012, Kennedy, 2010, Wisz et al., 2013). For example, in-vitro co-culture assays have suggested facilitative interactions between ectomycorrhizal fungi, such as *Hebeloma cylindrosporum* and *Paxillus involutus*, and certain fungal strains belonging to an endophytic fungal taxon *Leptodontidium* (Berthelot et al., 2019). Those facilitative interactions are potentially reflected in aggregated (co-occurrence) patterns of fungi in root tips (Berthelot et al., 2019; Kia et al., 2019; Yamamoto et al., 2014). In terms of antagonistic interactions between species, experimental studies have revealed that competition for space and carbon sources within terminal roots can occur between ectomycorrhizal fungi and endophytic fungi (Kennedy et al., 2009; Reininger & Sieber, 2012, 2013). Such competitive interactions are expected to structure fine-scale segregated distribution (Koide et al., 2005; Pickles et al., 2012). Thus, although aggregated and segregated spatial patterns do not always infer direct species interactions (Blanchet et al., 2020; Faust & Raes, 2012; Hirano & Takemoto, 2019), forest-scale analyses of potential facilitative and competitive interactions will provide pivotal insights into the organization of root-associated fungal communities.

The potential impacts of multiple ecological processes on species assembly can be addressed based on emerging statistical approaches (Ovaskainen et al., 2017; Pichler & Hartig, 2021; Warton et al., 2015, Leibold et al., 2022, Pollock et al., 2014). The flexible platform of joint species distribution modeling (jSDM), for example, enables the simultaneous evaluation of the impacts of multiple factors on community compositional patterns (Ovaskainen et al., 2017; Pichler & Hartig, 2021; Pollock et al., 2014). With jSDM, we can examine how biotic/abiotic environmental factors, spatial autocorrelations, and direct/indirect interactions between species collectively structure the distribution of tens or hundreds of species (Abrego et al., 2020; Hartig et al., 2024; Vass et al., 2022). Another promising statistical approach is to infer direct species interactions by controlling implicit (latent) environmental factors that explain distribution patterns of species constituting communities (Kurtz et al., 2019). The framework of sparse inverse covariance estimation for ecological associations (SPIEC-EASI), for example, provide a platform for separating effects of species interactions from those of environmental preference shared between species (Kurtz et al., 2015, 2019). The application of those statistical frameworks is expected to set a starting point for comprehensively understanding how multiple ecological factors and mechanisms drive the symbiosis linking below-ground and above-ground ecosystems.

In this study, we examine how multiple ecological factors could organize the assembly of fungi associated with plant roots. We sampled fine roots from diverse ectomycorrhizal and arbuscular mycorrhizal plant species at > 100 positions within a cool temperate forest in Japan, obtaining the data of fungal community structure for 1,615 fine root samples. We then performed a series of statistical modeling to evaluate how host plant preference, fungal community structure of background soil, spatial autocorrelation, and fungus–fungus interactions (or shared environmental preference) could collectively structure the spatial distribution of root-associated fungi. The analysis based on jSDM allowed us to examine how functional guilds of root-associated fungi (e.g., ectomycorrhizal, arbuscular mycorrhizal, and endophytic fungi) could vary in basic community-assembly rules. Furthermore, a SPIEC-EASI network inference with latent environmental variables enabled the exploration of species and interactions that potentially played key roles in the community processes of below-ground plant–fungal symbiosis. Overall, the statistical modeling of the entire root-associated fungal community provides a thorough platform for deciphering the diversity of basic assembly rules within the kingdom Fungi, suggesting functional complementarity of fungal guilds in below-ground ecosystem processes.

## MATERIALS AND METHODS

### Sampling

Sampling was conducted in the research forest of Sugadaira Research Station, Mountain Science Center, University of Tsukuba, Sugadaira, Ueda, Nagano Prefecture, Japan (36.524 °N; 138.349 °E; 1340 m a.s.l.). The forest was dominated by ectomycorrhizal tree species such as *Betula platyphylla* var. *japonica* and *Pinus densiflora*. Within the eight-ha secondary forest, we set 126 sampling positions, which were at least 4-m apart from each other (Figure S1). At each sampling point, ca. 30 cm^2^ soil was excavated from a depth of 0 to 20 cm in order to collect woody plant roots and bulk soil. Samples obtained here were immediately refrigerated in the field and frozen at −20 °C after being brought back to the laboratory. The location of each sampling site and the tree species occurring within 3-m range from each sampling site (> 4-cm diameter at breast height) were recorded. The obtained plant roots were classified according to their morphology at each sampling site, and eight terminal root fragments (1-cm in length) were collected for each morphological type. The roots were stored at −20 °C until DNA extraction.

At 30 sampling positions randomly selected from the 126 positions, additional soil samples were collected for the measurement of pH. After sieving, 10 g of each soil sample was shaken at 1.12 g for 30 minutes in 25 ml Milli-Q water. The pH of the soil samples was measured within a week after sampling by a portable pH meter LAQUAact D-74 (HORIBA).

For the identification of plant root samples based on DNA sequences, the leaves of the major component species of the forest were collected as a reference. The target species were *Pinus densiflora* (Pinaceae), *Betula platyphylla* var. *japonica* (Betulaceae), *Quercus mongolica* var. *grosseserrata* (Fagaceae), *Populus sieboldii* (Salicaceae), *Castanea crenata* (Fagaceae), *Corylus heterophylla* (Betulaceae), *Acer mono* (Sapindaceae), *Larix leptolepis* (Pinaceae), *Juglans ailanthifolia* (Juglandaceae), *Fagus crenata* (Fagaceae), and *Picea jezoensis* var. *hondoensis* (Pinaceae). For all these major component species other than *A. mono*, no congeneric species were found in the study forest: hereafter, we refer to these plant species with their genus names: for *A. mono*, some congeneric species (*A. crataegifolium* and *A. rufinerve*) were observed at low frequency in the research site. The leaf samples were immediately refrigerated in the field and then frozen at −20 °C after being brought back to the laboratory.

### DNA extraction and purification

The root samples were washed with sonication at 45 kHz for 3 min in 1 ml of 0.5% tween20. The samples were then surface sterilized with 3 ml of NaClO (Nakalaitesque, Sodium Hypochlorite Solution; effective chlorine concentration ca. 11%) diluted to 1% (v/v), rinsed with 3 ml of sterile distilled water, and immersed in 99.5% ethanol for 30 seconds.

DNA extraction was performed with a cetyltrimethylammonium bromide (CTAB) method followed by purification by Phenol/Chloroform/Isoamyl alcohol (25:24:1). The roots were frozen at −20 °C, freeze-dried overnight, and then pulverized with 1- and 4-mm zirconium ball mixture at 30 Hz for 3 minutes using a TissueLyser II (Qiagen). The pulverized root samples were centrifuged at 3,010 g for 3 minutes at 20 °C, then 600 μl of CTAB buffer (100 mM Tris; 1.4 M NaCl; 20 mM EDTA; 2% CTAB) was added, subsequently kept at 60 °C for 1 hour. The DNA extract was centrifuged at 3,010 g for 30 minutes at 20 °C, and 200 μl of the supernatant was collected. The supernatant was mixed with 200 μl of Phenol/Chloroform/Isoamyl alcohol (25:24:1) for 10 minutes and then centrifuged at 3,010 g for 30 minutes at 20 °C. DNA in 80 μl of the supernatant was then precipitated with 80 μl of ice-cold 100% 2-propanol 17,800 g for 10 minutes at 4 °C and subsequently washed with 100 μl of ice-cold 70% ethanol. After centrifugation at 17,800 g for 10 minutes at 4 °C, the supernatant was removed and the precipitate was air-dried for 15 minutes. To this was added 40 μl of 1 × tris-EDTA buffer [10 mM tris-hydroxymethylaminomethane solution (pH 8.0), 1 mM ethylenediaminetetraacetic acid (EDTA; pH 8.0)] to obtain purified DNA extract. The DNA extraction protocol was applied as well to leaf samples for plant species identification.

### PCR amplification

The fungal community compositions of the root and soil samples were analyzed by targeting the internal transcribed spacer 1 (ITS1) region. In the PCR of fungal ITS1 region, we used the forward primer ITS1F-KYO1 (Toju et al., 2012) fused with 3–6-mer Ns for improved Illumina sequencing quality (Lundberg et al., 2013) and the forward Illumina sequencing primer (5′-TCG TCGGCA GCG TCA GAT GTG TAT AAG AGA CAG-[3–6-merNs]–[ITS1F-KYO1]−3′) and the reverse primer ITS2-KYO2 (Toju et al., 2012) fused with 3–6-mer Ns and the reverse sequencing primer (5′-GTC TCG TGG GCT CGG AGA TGT GTA TAAGAG ACA G [3–6-mer Ns]—[ITS2-KYO2]−3′). The buffer and DNA polymerase kit of KOD One PCR Master Mix (Toyobo) was used with a temperature profile of 35 cycles at 98 °C for 10 seconds, 52 °C for 5 seconds, and 68 °C for 20 seconds, followed by a final extension step at 68 °C for 2 minutes. The ramp rate through the thermal cycles was set to 1 °C/second in order to prevent generation of chimeric sequences (Stevens et al., 2013). To add Illumina sequencing adaptors to respective samples, supplemental PCR was performed using the forward fusion primers consisting of the P5 Illumina adaptor, 8-mer indexes for sample identification (Hamady et al., 2008), and a partial sequence of the sequencing primer (5′-AAT GAT ACG GCG ACC ACC GAG ATC TAC AC – [8-mer index] -TCG TCG GCA GCG TC−3′) and the reverse fusion primers consisting of the P7 adaptor, 8-mer indices, and a partial sequence of the sequencing primer (5′-CAA GCA GAA GAC GGC ATA CGA GAT –[8-mer index] – GTC TCG TGG GCT CGG−3′). KOD One PCR Master Mix was used with a temperature profile of 8 cycles at 98 °C for 10 s, 55 °C for 5 seconds, 68 °C for 5 seconds (ramp rate = 1 °C/second), and a final extension at 68 °C for 2 minutes. The PCR amplicons of the samples were pooled with equal volume after a purification/equalization process with AMPureXP Kit (Beckman Coulter). The ratio of AMPureXP reagent to amplicons was set to 0.6 (v/v) in order to remove primer dimers (i.e., sequences shorter than 200 bp).

In addition to the PCR of the fungal ITS1 region, the internal transcribed spacer 2 (ITS2) region was amplified for the molecular identification of plant species, targeting root and reference leaf samples. In the PCR of plant ITS2 region, we used the forward primer ITS-3p62plF1 (Kolter & Gemeinholzer, 2021) and the forward Illumina sequencing primer (5′-TCG TCGGCA GCG TCA GAT GTG TAT AAG AGA CAG-[6-merNs]–[ITS-3p62plF1]-3′) and the reverse primer ITS-4unR1 (Kolter & Gemeinholzer, 2021) and the reverse sequencing primer (5′-GTC TCG TGG GCT CGG AGA TGT GTA TAAGAG ACA G-[6-merNs]– [ITS-4unR1]-3′). The buffer and DNA polymerase kit of KOD One PCR Master Mix (Toyobo) was used with a temperature profile of 35 cycles at 98 °C for 10 seconds, 52 °C for 5 seconds, and 68 °C for 20 seconds, followed by a final extension step at 68 °C for 2 minutes. The ramp rate through the thermal cycles was set to 1 °C/ seconds in order to prevent generation of chimeric sequences (Stevens et al., 2013). Illumina sequencing adaptors and 8-mer index sequences were then added in the second PCR as described above. The amplicons were purified and pooled as described above.

The sequencing libraries of the fungal ITS1 region and plant ITS2 region were processed in 12 runs of an Illumina MiSeq sequencer (15% PhiX spike-in). Because the quality of forward sequences is generally higher than that of reverse sequences in Illumina sequencing, we optimized the MiSeq run setting in order to use only forward sequences. Specifically, the run length was set 271 forward and 31 reverse cycles to enhance forward sequencing data: the reverse sequences were used only for screening ITS1 sequences within the output data.

### Bioinformatics

In total, 134,066,687 sequencing reads were obtained in the Illumina sequencing. The raw sequencing data were converted into FASTQ files using the program bcl2fastq 1.8.4 distributed by Illumina. The output FASTQ files were demultiplexed with the program Claident v0.9.2022.01.26 (Tanabe & Toju, 2013, Tanabe, 2022). Sequencing reads whose 8-mer index positions included nucleotides with low (< 30) quality scores were removed in this process.

For each dataset of the ITS1 and ITS2 regions, filtering of the sequencing reads was performed with the program Cutadapt (Martin, 2011) v.3.7 and DADA2 (Callahan et al., 2016) v.1.18.0 of R 4.2.2 (R core team, 2022) to trim target sequences and remove low-quality data. We then obtained amplicon sequence variants (ASVs) using DADA2. Detection of contaminat ASVs were performed with ‘decontam’ function (method = “prevalence”) of the “decontam” package (Davis et al., 2018). To merge all the sequencing results, we performed “mergeSequenceTables” function of the DADA2 package. Taking into account potential intragenomic variation of fungal ITS1 & plant ITS2 sequences, the ASVs were re-clustered into operational taxonomic units (OTUs) with a 97% similarity threshold using the program VSEARCH v2.21.1 (Rognes et al., 2016). To evaluate potential effects of the cutoff similarity setting on the downstream statistical results, we performed an additional re-clustering of ASVs into OTUs at 93 % sequence similarity threshold for the fungal ITS1 dataset. For each cutoff similarity setting (97% or 93%) of the ITS1 region, the taxonomic assignment of fungal OTUs was performed based on the UNITE General FASTA release database version 9.0 for Fungi (Abarenkov et al. 2023) using the “AssignTaxnomy” function of the DADA2 package (Callahan et al., 2016). For the plant ITS2 region, taxonomic assignment was performed with the five-nearest-neighbor method (Tanabe & Toju, 2013) using the reference sequences identified to genus in the BLAST nt sequence database (version 2022-07-15).

For the identification of plant roots, a molecular phylogenetic analysis was performed with the neighbor-joining method using MEGA X (Glen al. 2020) by targeting the ITS2 sequences identified as Viridiplantae (Figure S2). The ITS2 sequences that formed monophyletic groups (≥ 90% bootstrapping probabilities; 1,000 iterations) with those from reference leaf samples were identified to species. Additional BLAST search of plant ITS2 sequences were performed to supplement the molecular identification. Samples from which sequences from multiple plant genera were detected were not included in the following analyses.

For the fungal ITS1 dataset of the root and soil samples, functional guilds (guilds) of fungal OTUs were inferred using FungalTraits database (Põlme et al., 2020) with reference to the identification results at the genus level. In the FungalTraits database, the following “primary_lifestyles” associated with plant roots were targeted: mycoparasites (“mycoparasite”), root endophytes (“root_endophyte”), ectomycorrhizal fungi (“ectomycorrhizal”), and arbuscular mycorrhizal fungi (“arbuscular_mycorrhizal”), and plant root pathogens (“plant_pathogen” in “primary_lifestyle” and “root_pathogen” in “plant_pathogenic_capacity_template”). Likewise, the following “secondary_lifestyles” were explored within the dataset: mycoparasites (“mycoparasite”), root-associated fungi (“root-associated”), root endophytes (“root_endophyte” or “root_endophyte_dark_septate”), ectomycorrhizal fungi (“ectomycorrhizal”), arbuscular mycorrhizal fungi (“arbuscular_mycorrhizal”), nematophagous (“nematophagous” in “secondary_lifestyle” or “animal_parasite” in “secondary_lifestyle” and “nematophagous” in “animal_biotrophic_capacity_template”), plant root pathogens (“plant_pathogen” in “secondary_lifestyle” and “root_pathogen” in “plant_pathogenic_capacity_template”). If a different functional guild was inferred for primary_lifestyle and secondary_lifestyle within FungalTrait, the one for primary_lifestyle was used as the functional guild information of a target OTU. The guilds of some fungal genera whose lifestyle in the root were well characterized in previous studies but different guilds were inferred by the above automatic guild assignment were corrected manually (Table S1). Ericoid mycorrhizal fungi were not considered as a guilds because no root samples in our dataset was identified as Ericaceae. In addition to these guilds inference, fungal OTUs whose phylum were annotated as Glomeromycota by the above taxonomic assignment were also regarded as arbuscular mycorrhizal fungi. In the following analyses, ectomycorrhizal fungi were denoted as “EcMF,” arbuscular mycorrhizal fungi as “AMF,” mycoparasites as “Mycoparasite,” root endophytes as “Endophyte”, plant root pathogen as “Pathogen”, nematophagous fungi as “Nematophagous”. Fungal OTUs inferred as root-associated fungi were denoted as “other_RAF” (other root associated fungi), because they were not regarded as any other specific guilds in the above criteria. Fungal OTUs unidentified at the genus level were denoted as “Unidentified”, while those identified at the genus level but unassigned to any functional guild were denoted as “Unassigned”.

Coverage-based rarefaction was performed, respectively, for root and soil fungal community matrices, in which rows represented samples and columns denoted fungal OTUs. Before the rarefaction, we discarded root samples with less than 1,000 reads and soil samples with less than 5,000 reads. Root samples for which genus-level plant information was unavailable were discarded as well. For each of the root and soil dataset, rarefaction was performed with the lowest coverage score among the samples (root, > 99.9%; soil, > 99.9%) using the “vegan” package v.2.6-4 (Oksanen et al., 2022) of R. Relationships between the number of the sequencing reads and that of detected fungal OTUs were examined for both root and soil fungal datasets with the “rarecurve” function of the “vegan” package. Likewise, relationship between the number of the samples and that of detected fungal OTUs was examined with the “specaccum” function (10,000 trials). In the following analyses, the rarefied datasets were used unless otherwise noted. After a series of the data filtering detailed above, fungal community data of background soil were unavailable for two out of the 126 sampling positions examined: the two sampling positions were excluded from the following statistical analyses. In total, fungal community data were obtained for 1,615 root samples collected from the 124 sampling positions (Figure S3). In total, 2,351 OTU and 2,834 OTU were obtained from those root and soil samples, respectively (Figure S4).

### Joint species distribution modeling

The distribution of fungal OTUs in root samples was modeled based on the framework of jSDM (Ovaskainen et al., 2017; Pichler & Hartig, 2021; Pollock et al., 2014) using the “sjSDM” package (version 1.0.1; Pichler & Hartig, 2021) of R. The factors included in the analysis were host plant species (P), the fungal community structure of background soil (S), spatial autocorrelation (Sp), and fungus–fungus covariance (co-occurrence) patterns (Cov). In addition to the full model including all the four variables, null model and 14 partial models including one, two, or three of the factor(s) were constructed (Table 1). Before the modeling, the read-count data of fungal OTUs were converted into binary data (i.e., presence or absence). Among the explanatory variables (factors), soil fungal community structure was included as the principal coordinates of the fungal OTU compositions between soil samples. Bray-Curtis dissimilarity (*β*-diversity) of fungal OTU compositions were used in the principal coordinate analysis (pCoA): values of pCoA axes 1 to 68 were used so that the cumulative contribution exceeded 90%. As spatial autocorrelation, the eigenvectors of spatial coordination of the sampling points were used. The “generateSpatialEV” function of the “sjsdm” package was used to convert the spatial coordination into eigenvectors. The root samples of infrequently occurred host plants (< 30 root samples) were excluded from the analysis. To evaluate the validity of the full model including all the variables, a receiver operating characteristic (ROC) curve was drawn and then area under the curve (AUC) was calculated. The statistical analysis was performed for both 97% and 93% OTU-cutoff criteria to confirm the stability of the results.

**Table 1:** Models examined in the jSDM framework. The explanatory variables examined in each of the full and partial models are presented. The repertoires of the explanatory variables are host plant species (P), the fungal community structure of background soil (S), spatial autocorrelation (Sp), and fungus–fungus covariance (co-occurrence) patterns (Cov).

|  | Model name | Factors in the models |  |  |  |
| --- | --- | --- | --- | --- | --- |
|  |  | Host plant | Soil fungal community | Spatial autocorrelation | Inter-OTU covariance |
| Model 1 | Full ( P + S + Sp + Cov ) | ○ | ○ | ○ | ○ |
| Model 2 | P + S + Sp | ○ | ○ | ○ |  |
| Model 3 | P + S + Cov | ○ | ○ |  | ○ |
| Model 4 | P + Sp + Cov | ○ |  | ○ | ○ |
| Model 5 | S + Sp + Cov |  | ○ | ○ | ○ |
| Model 6 | P + S | ○ | ○ |  |  |
| Model 7 | P + Sp | ○ |  | ○ |  |
| Model 8 | P + Cov | ○ |  |  | ○ |
| Model 9 | S + Sp |  | ○ | ○ |  |
| Model 10 | S + Cov |  | ○ |  | ○ |
| Model 11 | Sp + Cov |  |  | ○ | ○ |
| Model 12 | P | ○ |  |  |  |
| Model 13 | S |  | ○ |  |  |
| Model 14 | Sp |  |  | ○ |  |
| Model 15 | Cov |  |  |  | ○ |
| Model 16 | Null |  |  |  |  |

To evaluate the explanatory power for the spatial distribution of the fungal OTUs, the log-likelihood of each of the 16 models (full model + 14 partial models + null model) was calculated. Moreover, to evaluate the influence of each factor on the distribution of each guild of fungi (Leibold et al., 2022), contribution of each fungal OTU to the log-likelihood ratio between models was calculated. In the evaluation of the effects of fungus–fungus covariance, we used the log-likelihood ratio between the full model and the model lacking fungus–fungus covariance (P + S + Sp). Meanwhile, in the evaluation of the effects of host plants, soil fungal community, or spatial autocorrelation, the model lacking fungus–fungus covariance (P + S + Sp), but not the full model, was used as the model to be compared because fungus–fungus covariance might reflect shared responses to unaccounted environmental factors. Consequently, a model with soil fungal community and spatial autocorrelation (S + Sp), that with plant identity and spatial autocorrelation (P + Sp), and that with plant identity and soil fungal community (P + S) were compared with the model lacking fungus–fungus covariance (P + S + Sp) in the evaluation of the effects of host plants, soil fungal community, and spatial autocorrelation, respectively. To compare the log-likelihood ratios among fungal guilds, Steel-Dwass tests were performed: fungal functional guilds including five or more OTUs were examined in the tests.

In a supplementary analysis, potential impacts of soil environmental conditions were evaluated. Specifically, by targeting the 29 positions at which soil pH information was available, a joint species distribution model including soil pH and the abovementioned four variables was constructed. The root samples of infrequently occurred host plants (< 15 root samples) were excluded from the analysis. To evaluate the explanatory power of the full model, AUC was calculated. In addition to the full model, null model and 30 partial models including one, two, three, or four of the five factors were constructed: log-likelihoods of these model were calculated (Table S2).

### Host-plant preference

The host-plant preference of each fungal OTU was evaluated based on the *d* ′ metric of interaction specificity (Blüthgen et al., 2006). To evaluate the statistical significance of the host preference patterns, a randomization test was performed. Specifically, the host plant information was shuffled among root samples collected at the same sampling positions (10,000 permutations) and *d* ′ values calculated based on the randomized community matrices were compared with the *d* ′ value calculated from the original data matrix. For each fungal OTU, a standardized host preference score was obtained as follows:

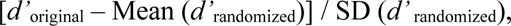

where *d’*_original_ was the *d’* estimate of the original data, and Mean (*d’* _randomized_) and SD (*d’* _randomized_) were the mean and standard deviation of the *d’* scores of randomized data matrices. False discovery rates [FDR; (Benjamini & Hochberg, 1995)] were then calculated to evaluate the statistical significance of the preferences (two-tallied tests). Using the original and randomized matrices, plants’ preferences for fungi were evaluated as well based on the standardized *d* ′ metric. Furthermore, specificity of each plant-fungus associations (two-dimensional preference) was evaluated with the randomization approach (Toju et al., 2016). Specifically, the extent to which focal fungus–plant associations were observed more frequently than that expected by chance was calculated for a pair of a fungal OTU *i* and plant species *j* as follows:

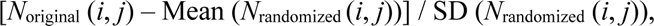

where *N*_original_ (*i*, *j*) denoted the number of root samples from which a focal combination of a fungal OTU and a plant species was observed in the original data, and the Mean (*N*_randomized_ (*i*, *j*)) and SD (*N*_randomized_ (*i*, *j*)) were the mean and standard deviation of the number of samples for the focal fungus-plant pair across randomized matrices. Prior to the above preference analyses, the data matrices were converted into a binary format (i.e., presence/absence of fungal OTUs). The fungal OTUs that occurred in less than 30 root samples among were discarded. Likewise, as in the jSDM, the root samples of infrequently occurred host plants (< 30 root samples) and samples collected at the two positions lacking soil fungal community information was excluded.

### Habitat preference: roots vs. soil

To gain an overview of fungal habitat preference, each fungal OTU was plotted on a two-dimensional surface representing occurrences in root and soil samples. In addition, each fungal OTU was plotted on another surface illustrating mean relative abundance (log-transformed proportion of sequence read counts) in root and soil samples.

The habitat preference of each fungal OTU was statistically examined in a generalized linear model (GLM) of occurrences in the roots. At each sampling position, the number of root samples from which a focal fungal OTU was detected was counted for each host plant species. In parallel, for the soil sample of each sampling site, the relative abundance (log-transformed proportion of sequence read counts) of the focal fungal OTU was calculated. A GLM of each fungal OTU occurrence in root samples was then constructed by including relative abundance of the OTU in soil samples and plant species identity as explanatory variables, and the total sample number of each host plant genus at each site as an offset term (log link-function and Poisson errors). A Steel-Dwass test was performed to evaluate differences in the standardized partial regression coefficients of the soil relative abundance between fungal guilds. The fungal OTUs that occurred in at least 30 root samples and 15 soil samples were subjected to the analysis.

### Spatial autocorrelation

To determine the extent of spatial autocorrelation in the root-associated fungal community, Mantel’s correlogram analysis was performed. The effects of geographic distances between sampling sites on dissimilarity in fungal community structure were estimated by “mantel.correlog” function of “vegan” package (10,000 permutations) with Jaccard *β*-diversity calculated based on the binary fungal community data (presence/absence). The Mantel’s correlogram analysis was performed as well for subset datasets that included specific fungal guilds [arbusculr mycorrhizal fungi (“AMF”), ectomycorrhizal fungi (“EcMF”), endophytes (“Endophyte”), plant pathogens ““Pathogen”), mycoparasites ““Mycoparasite”), nematophagous fungi ““Nematophagous”) or unclassified root-associated fungi (“Other_RAF”)].

### Fungus–fungus network analysis

We examined how direct fungus–fungus interactions could influence the assembly of root-associated fungi based on the framework of sparse inverse covariance estimation for ecological associations (SPIEC-EASI) (Kurtz et al., 2015, 2019). To extract direct fungus–fungus interactions from the patterns in the coexistence (co-occurrence), we constructed the model including latent variable as implemented with the “sparse and low rank” (SLR) method of SPIEC-EASI (Kurtz et al., 2019) with un-rarefied datasets (as recommended in SPIEC-EASI). Based on models with latent (implicit) environmental variables, the SLR method allows us to distinguish co-occurrence associations resulting from direct fungus–fungus interactions from patterns caused by shared environmental preferences. We first constructed models including 0 to 20 latent variables and then selected the optimal number of latent variables in light of Bayesian information criterion (BIC): i.e., the model with the smallest BIC score was selected. The selected models was used to infer the architecture of positive/negative association network. For the positive association network, modules of densely associated sets of fungal OTUs were inferred with the Louvain algorithm as implemented in “cluster_louvain” function of “igarph” package (version 1.4.2; Csardi and Nepusz, 2006).

Within the fungus–fungus association networks, fungal OTUs with high degree centrality (i.e., the number of connected nodes) was explored. For each of the OTUs highlighted, we quantified the extent to which its spatial distribution was determined by fungus–fungus covariance in the jSDM. Specifically, the log-likelihood ratio between the model including all factors (Full (P + S + Sp + Cov)) and the model from which fungus–fungus covariance was removed (P + S + Sp) was calculated for each fungal OTU. Thus, fungal OTUs with strongest signs of interactions with other fungal species were explored based on two lines of statistical approaches. Consistency between the results of SPIEC-EASI and those of jSDM was examined with Kendall rank correlation test.

## RESULTS

### Fungal diversity and community compositions

Within the fungal community of the root samples, Russulales (the propotion of sequencing read counts = 27.3%), Helotiales (22.9%), Agaricales (14.1%), and Thelephorales (11.3%) were most abundant (Figure 1a, Figure S5b). At the genus level, *Russula* (22.6 %), *Tomentella* (8.9%), *Hyaloscypha* (8.5%), and *Phialocephala* (5.1%) showed the highest relative abundance (Figure 1b, Figure S5a). In terms of functional guilds, ectomycorrhizal fungi were the most abundant (52.1%), followed by endophytic fungi (12.4%), unclassified root-associated fungi (3.3%), arbuscular mycorrhizal fungi (2.5%), and rare functional guilds (mycoparasite, root pathogen and nematophagous fungi) (Figure 1c, Figure S5c).

**Figure 1:**
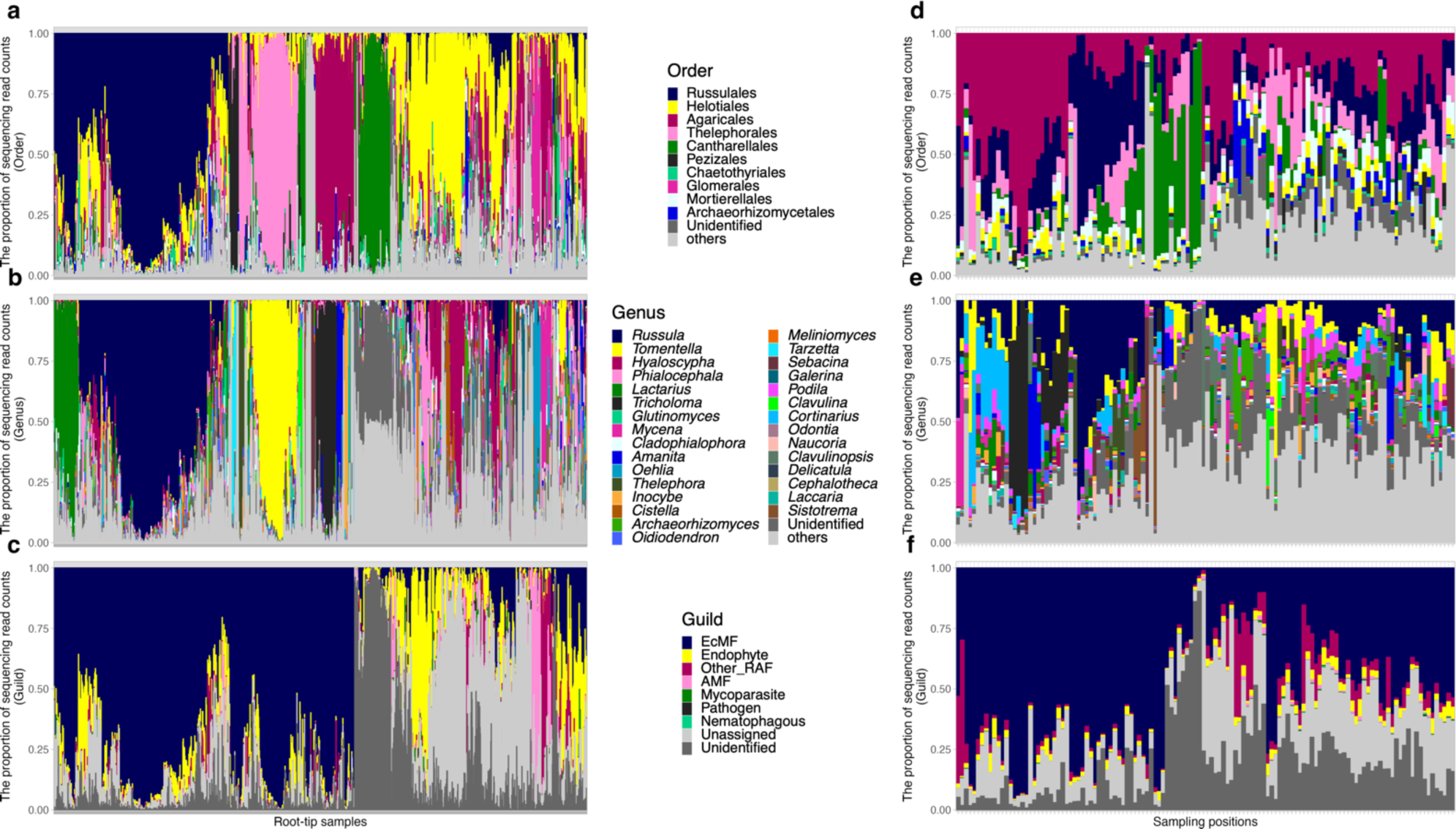
Root and soil fungal community compositions. (a) Order-level taxonomic compositions (root). The proportions of fungal sequencing reads are shown for each of the 1,615 root-tip samples. (b) Genus-level taxonomic compositions (root). (c) Functional guild compositions (root). The functional guilds of fungal OTUs were inferred with the Fungaltraits database. (d) Order-level taxonomic compositions (soil). The proportions of fungal sequencing reads are shown for each of the 124 soil samples. (e) Genus-level taxonomic compositions (soil). (f) Functional guild compositions (soil).

Within the fungal community of the soil samples, Agaricales (27.1%), Russulales (17.8%), Thelephorales (10.5%), and Cantharellales (10.3%) were most abundant (Figure 1d). At the genus level, *Russula* (15.9%), *Tomentella* (6.4%), *Tricholoma* (5.9%), and *Cortinarius* (5.0%) showed the highest relative abundance (Figure 1e). Fungal functional guilds were inferred for 62.2% of the total read counts [ectomycorrhizal fungi (53.4%), unclassified root-associated fungi (5.8%), endophytic fungi (2.5%), arbuscular mycorrhizal fungi (0.29%); Figure 1f].

### Ecological factors of fungal community assembly

In jSDM, the full model (Table 1), which included host plant identity, soil fungal community structure, spatial autocorrelation, and fungus–fungus covariance as explanatory variables, best predicted the spatial distribution of fungi at both 97% and 93% thresholds of fungal OTU definition (97%, AUC = 0.889; 93%, AUC = 0.873; Figure S6). Specifically, at the 97% cutoff similarity, the full model involving all four factors showed the highest log-likelihood, followed by a model involving host plant identity, soil fungal community structure, and fungus–fungus covariance (P + S + Cov; Figure 2). At both cutoff similarities of fungal OTU definition, the top four models with the highest log-likelihoods involved fungus–fungus covariance (Figure 2, Figure S7). On the other hand, models with only host plant identity (P) and those with only soil fungal community structure (S) as well as models with both factors (P + S) showed lowest log-likelihoods comparable to those of null models (Figure 2, Figure S7).

**Figure 2:**
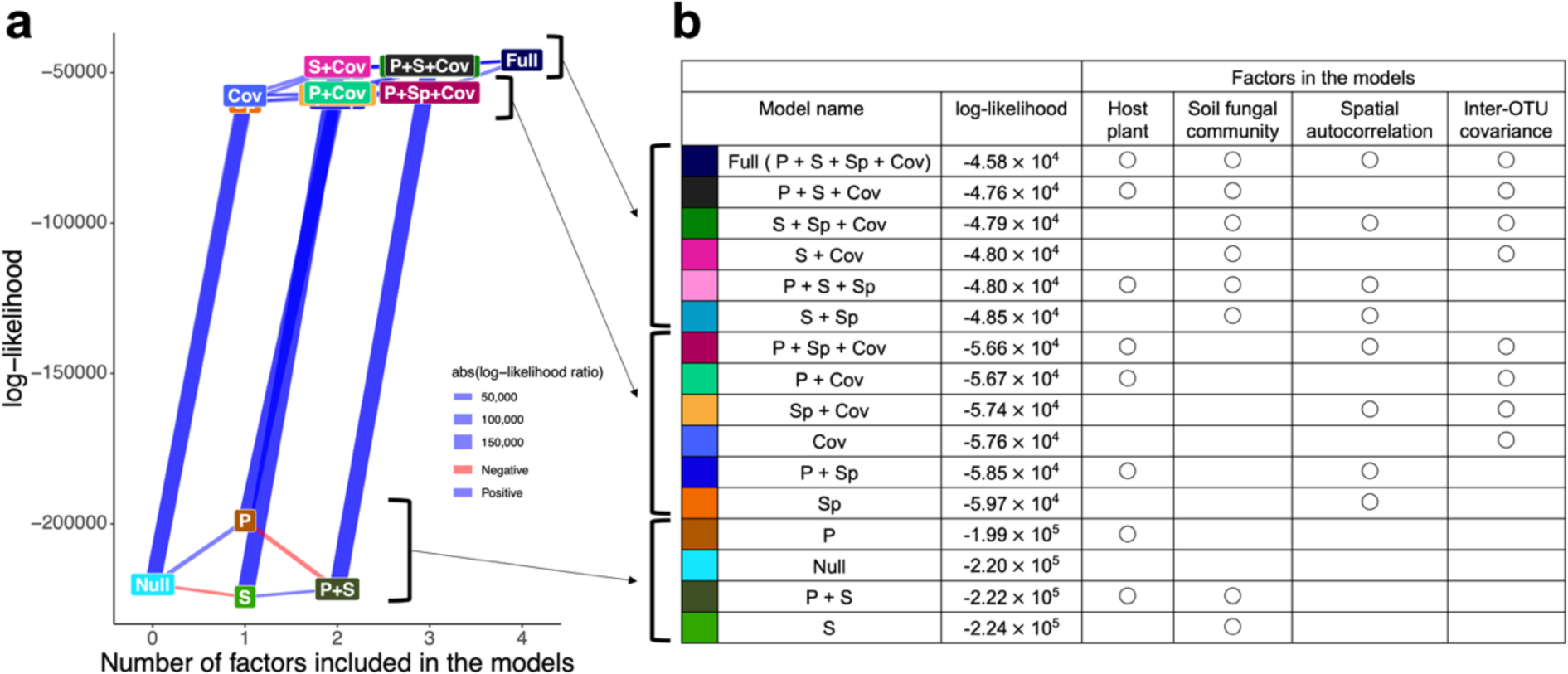
Comparison of the full and partial models in jSDM. (a) Performance of the models. The log-likelihoods of the models (Table 1) are shown. The pairs of full/partial models differing only in the presence/absence of a single explanatory variable are linked with edges, whose thickness represent log-likelihood ratios of the paired models. The OTUs in the input data were defined at the 97%. See results at the 93% OTU cutoff similarity for Figure S7. (b) Models aligned in decreasing order of log-likelihoods. The examined models were roughly classified into three groups in terms of their log-likelihoods.

These patterns were consistent in the supplemental jSDM that involved soil pH as an additional factor (Figure S6, Figure S8). In the supplementary analysis, the full model consisting of all factors showed the highest log-likelihood, followed by seven partial models involving fungus–fungus covariance (Figure S8). Meanwhile, the model that considered only soil pH showed much lower log-likelihoods than the null model (Figure S8; Table S2).

We next evaluated the impacts of each ecological factor on the distribution of each fungal OTU in root samples. The effects of each ecological factor, which was evaluated in terms of log-likelihood ratios between models involving a focal factor and models without it, varied among the functional guilds of root-associated fungi (Figure 3; Table S3-6). At the 97% threshold of fungal OTU definition, for example, host plant identity had the strongest impacts on the distribution of arbuscular mycorrhizal and unclassified root-associated fungi, and the lowest impact on ectomycorrhizal fungi within the study forest dominated by ectomycorrhizal plant species (Figure 3a). Meanwhile, endophytic fungi showed intermediate levels of influence by host plant identity (Figure 3a). In contrast to the patterns observed for host plant identity, the impacts of soil fungal community structure were the highest for ectomycorrhizal fungi (Figure 3b). Likewise, spatial structure had the strongest impacts on the distribution of ectomycorrhizal fungi, while spatial autocorrelation was less evident in endophytic and unclassified root-associated fungi (Figure 3c). Fungus–fungus covariance had greater impacts on the distribution of endophytic fungi in root samples than on ectomycorrhizal and arbuscular mycorrhizal fungi (Figure 3d). The overall patterns were consistent between 97% and 93% cutoff similarity analyses (Figure 3), although the differences of the factors’ impact among fungal functional guilds were weaker at the 93% cutoff similarity than at the 97% cutoff similarity possibly due to reduced number of fungal OTUs (Figure 3e-f). At the 93% threshold analysis, the impacts of host plant identity, spatial structure, and fungus–fungus covariance differed significantly among some fungal functional guilds, while no significant difference was observed for spatial structure (Figure 3e-f).

**Figure 3:**
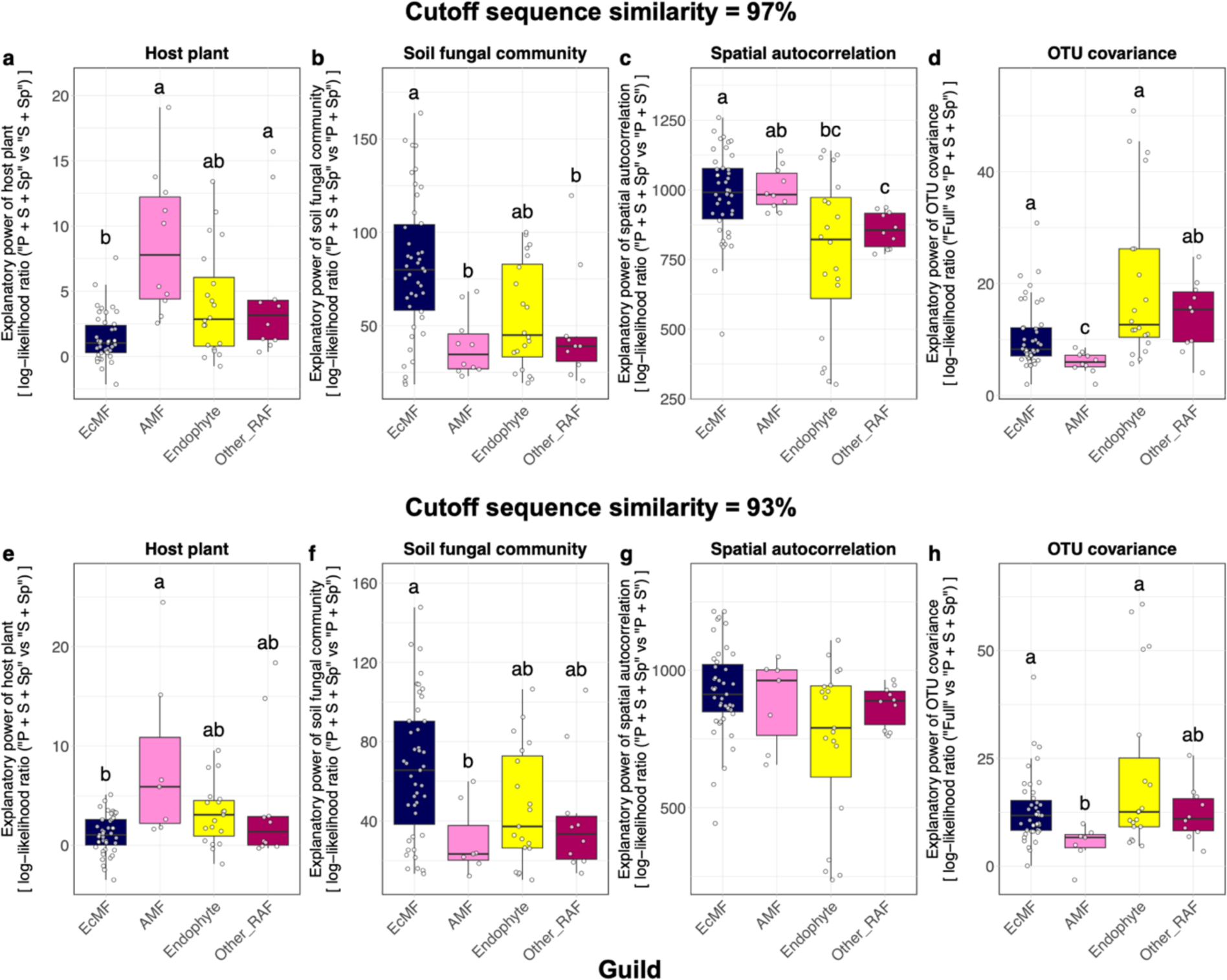
Comparison of the importance of ecological factors/processes among fungal functional guilds. (a) Host plant identity. The spatial distribution of each OTU was modeled in the jSDM framework. Subsequently, the two models differing in the presence/absence of host plant identity (“P + S + Sp” vs. “S + Sp”; see Table 1) was compared to evaluate the contribution of the explanatory variable. The log-likelihood ratios of the compared models were calculated for respective OTUs, which were grouped into four categories [ectomycorrhizal fungi (EcMF), arbuscular mycorrhizal fungi (AMF), endophytic fungi (Endophyte), and other root-associated fungi (Other_RAF)]. Difference in the log-likelihood ratios among fungal functional guilds were examined with Steel-Dwass test (significant differences were indicated by different letters). The OTUs in the input data were defined at the 97% cutoff similarity. (e-h) Results at the 93% OTU cutoff similarity.

### Host plant preference

Among the 185 fungal OTUs targeted in the randomization analysis, statistically significant host preferences were detected for 32 OTUs in standardized host preference (*d′* metrics) or randomization analysis (two-dimensional preference) (Figure 4, Table S7, Table S8). Except for arbuscular mycorrhizal fungi (e.g. X_00511, X_02153), which inevitably showed high host preferences in the study forest dominated by ectomycorrhizal plants, Venturiales sp. (X_07951; *d′* = 8.15, FDR < 0.001), *Pezicula ericae* (X_01277; *d′* = 6.82, FDR < 0.001), *Phialocephala fortinii* (X_00176; *d′* = 5.46, FDR < 0.001), *Archaeorhizomyces* sp. (X_09467; *d′* = 5.34, FDR < 0.001), *Cladophialophora* sp. (X_00235; *d′* = 3.18, FDR < 0.05), *Russula ematica* (X_00700; *d′* = 2.84, FDR < 0.05), and *Hyaloscypha* sp. (X_00682; *d′* = 5.06, FDR < 0.001), for instance, showed significant host plant preferences (Figure 4, Table S7). Among 10 plant species (genera) analyzed, *Pinus* (*d′* = 10.5, FDR < 0.001), *Acer* (*d′* = 7.52, FDR < 0.001), *Betula* (*d′* = 6.05, FDR < 0.001), *Toxicodendron* (*d′* = 4.05, FDR = 0.001), *Populus* (*d′* = 3.18, FDR = 0.004), *Juglans* (*d′* = 2.65, FDR = 0.010), and *Larix* (*d′* = 2.64, FDR = 0.010) showed statistically significant preferences for root-associated fungal OTUs (Figure 4).

**Figure 4:**
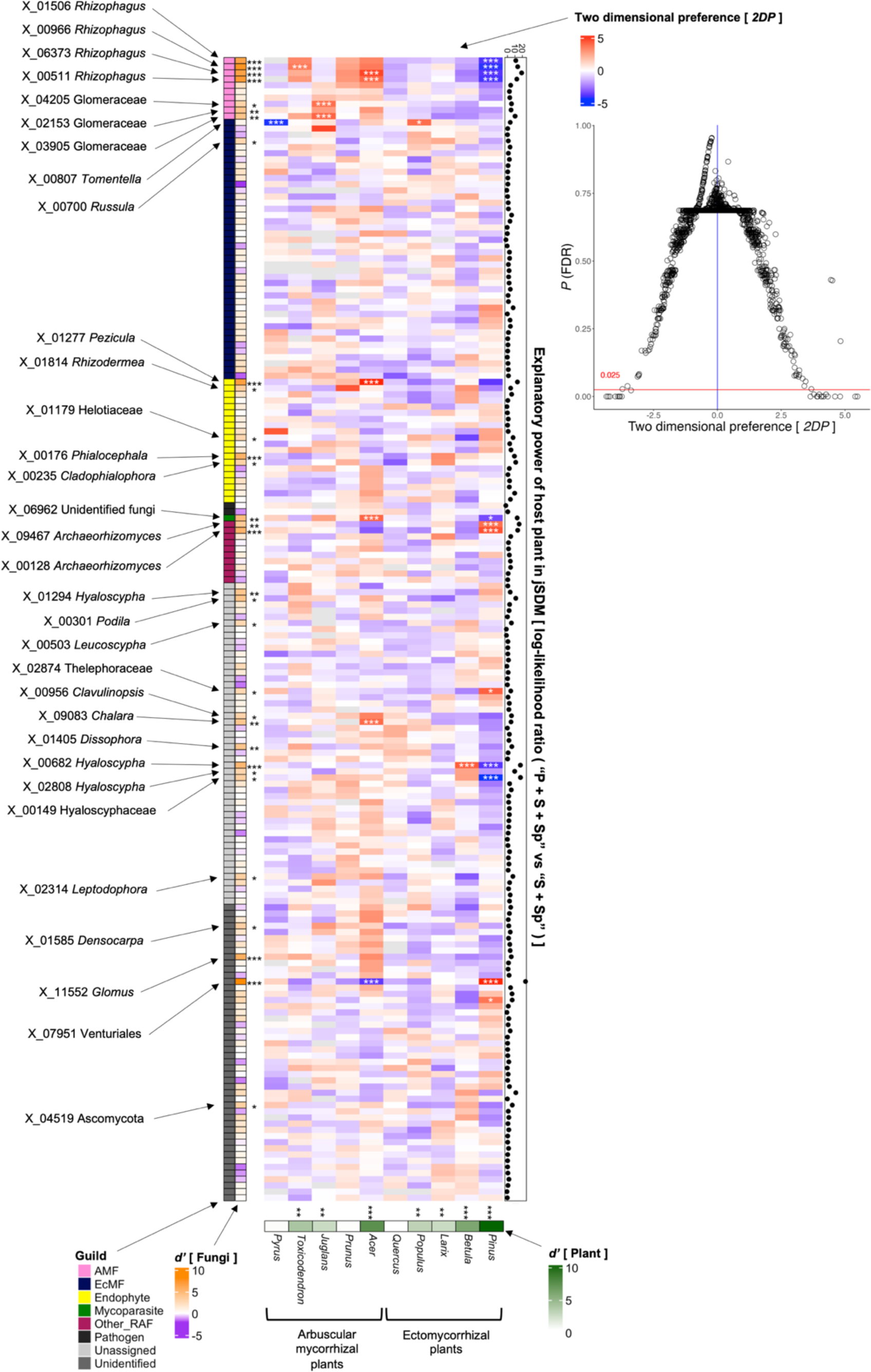
Preferences in plant–fungus associations. The standardized *d’* estimate of preference for host plant genus is shown for each of the 185 fungal OTUs (defined at the 97% cutoff sequence similarity) analyzed in the jSDM (rows). Likewise, the standardized *d’* estimate of preference for fungal OTUs is shown for each plant (columns). A cell in the matrix heatmap represents a two-dimensional preference (*2DP*) estimate, which indicate the extent to which the association of a target plant–fungal pair is observed more/less frequently than expected by chance. The relationship between *2DP* estimates and FDR-corrected *P* values is shown in the right panel. The log-likelihood ratio of the two models differing in the presence/absence of host plant identity (“P + S + Sp” vs. “S + Sp”; see Table 1) is shown for each fungal OTU on the right side of the heatmap. Significant preferences estimated in the standardized *d’* (one-tailed test) and *2DP* (two-tailed test) metrics are indicated by asterisks (***, *P* (FDR) < 0.001; **, *P* (FDR) < 0.01; *, *P* (FDR) < 0.05). The OTU ID and taxonomy of the fungal OTUs with significant preferences for plants are shown.

In the randomization analysis, statistically significant preferences were observed in several pairs of fungal OTUs and plant species such as *Tomentella terrestris* [X_00807] and *Populus*, *Pezicula ericae* [X_01277] and *Acer*, *Archaeorhizomyces borealis* [X_00128] and *Pinus*, *Hyaloscypha* sp. [X_00682] and *Betula*, and Venturiales sp. [X_07951] and *Pinus* (Figure 4, Table 2). Many of the fungal OTUs exhibiting significant associations with specific plant species also showed strong signs of host preferences in the jSDM (Figure 4, Table S3). In contrast, several fungus–plant associations were observed much less frequently than expected by chance: e.g., *Tomentella terrestris* [X_00807] and *Pyrus*, Venturiales sp. [X_07951] and *Acer*, Hyaloscyphaceae sp. [X_00149] and *Pinus* (Figure 4, Table S8).

**Table 2:**
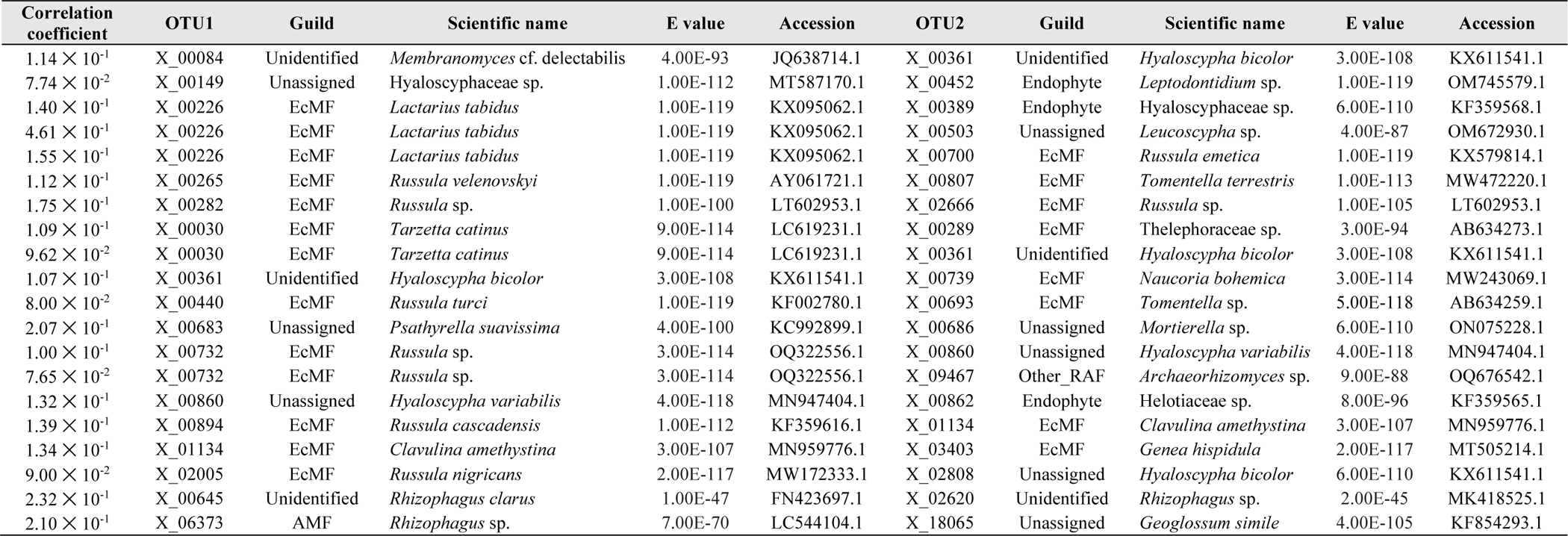
Within-root positive associations between fungi. With the SLR method of SPIEC-EASI analysis, fungus–fungus associations were inferred by controlling the potential effects of shared environmental preference (Figure 8a). This list includes the top-20 fungal OTU pairs in which positive associations were suggested based on the analysis. The OTUs were defined at the 97% cutoff similarity. For each fungal OTU pair, the correlation coefficient estimated with SPIEC-EASI was shown with the OTU ID, the functional guild automatically assigned by the Fungaltraits database, and BLAST top-hit results (the scientific name of the matched sequence, E-value, and NCBI accession number) of each fungus. Pairs of arbuscular mycorrhizal fungal OTUs are excluded from the list because they can involve erroneous inference of interactions resulting from the peculiar reproductive mechanisms of arbuscular mycorrhizal fungi (the presence of multiple genomes in single individuals).

### Habitat preference: roots vs. soil

Among the 138 OTUs analyzed, some endophytic fungi appeared frequently in the root samples (e.g., *Phialocephala fortinii* [X_00176], *Leptodontidium* sp. [X_00452], Figure 5a). While these fungi were also detected with high frequencies in the soil, they did not have large relative abundance in soil fungal communities (Figure 5a,b). Notably, *Phialocephala fortinii* (X_00176) appeared most frequently in root samples (62.2% of analysed root samples) and had the second largest relative abundance in root fungal communities. In contrast to these fungi, some ectomycorrhizal fungi, which had large relative abundance in root, also had large abundance in the soil (e.g. *Russula* sp. [X_00120], *Russula vesca* [X_00218]; Figure 5a,b). In addition to these fungi, *Membranomyces* sp. [X_00084] and Hyaloscyphaceae sp. [X_00149], respectively, showed similar patterns of habitat preference as endophytic and ectomycorrhizal fungi (Figure 5).

**Figure 5:**
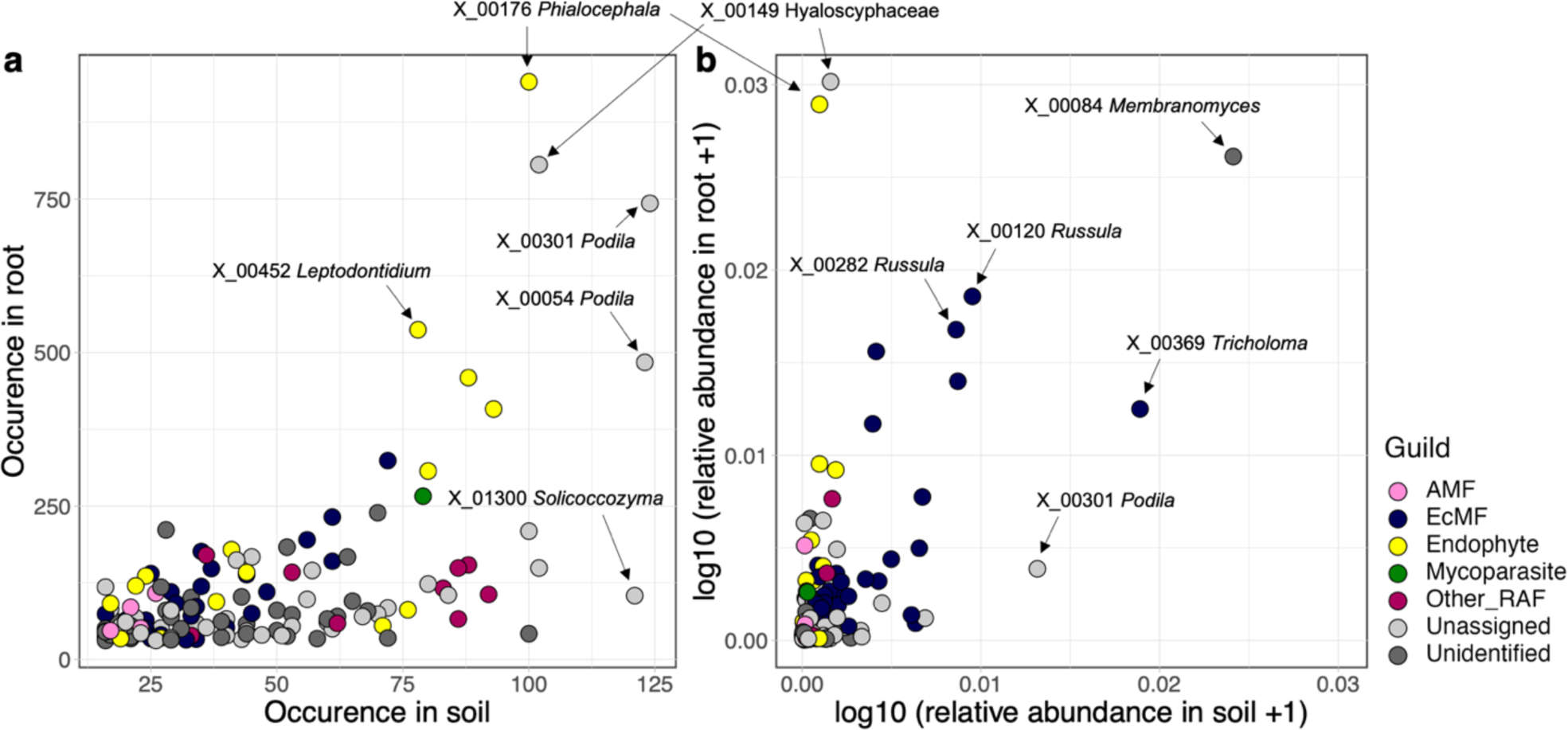
Habitat preference. (a) Two-dimensional surface of fungal OTU occurrence. The numbers of root and soil samples from which target fungal OTUs (defined at the 97% cutoff sequence similarity) are shown, respectively, along the vertical and horizontal axes. Fungal OTUs that appeared in ≥ 30 root samples and ≥ 15 soil samples were targeted in the analysis. (b) Two-dimensional surface of fungal OTU relative abundance. The proportions of sequence read counts in root and soil samples are shown for each fungal OTUs along the vertical and horizontal axes, respectively.

A series of GLMs indicated that positive relationship between occurrence in roots and abundance in background soil were more conspicuous for ectomycorrhizal fungi than for endophytic and other root-associated fungi (Figure 6a). The list of fungal OTUs whose distribution in roots was significantly explained by distribution patterns in the soil involved 30 ectomycorrhizal and seven endophytic fungi (Table S9). For the ectomycorrhizal OTUs, inclusion of soil fungal community structure improved prediction skills in jSDM (Figure 6b).

**Figure 6:**
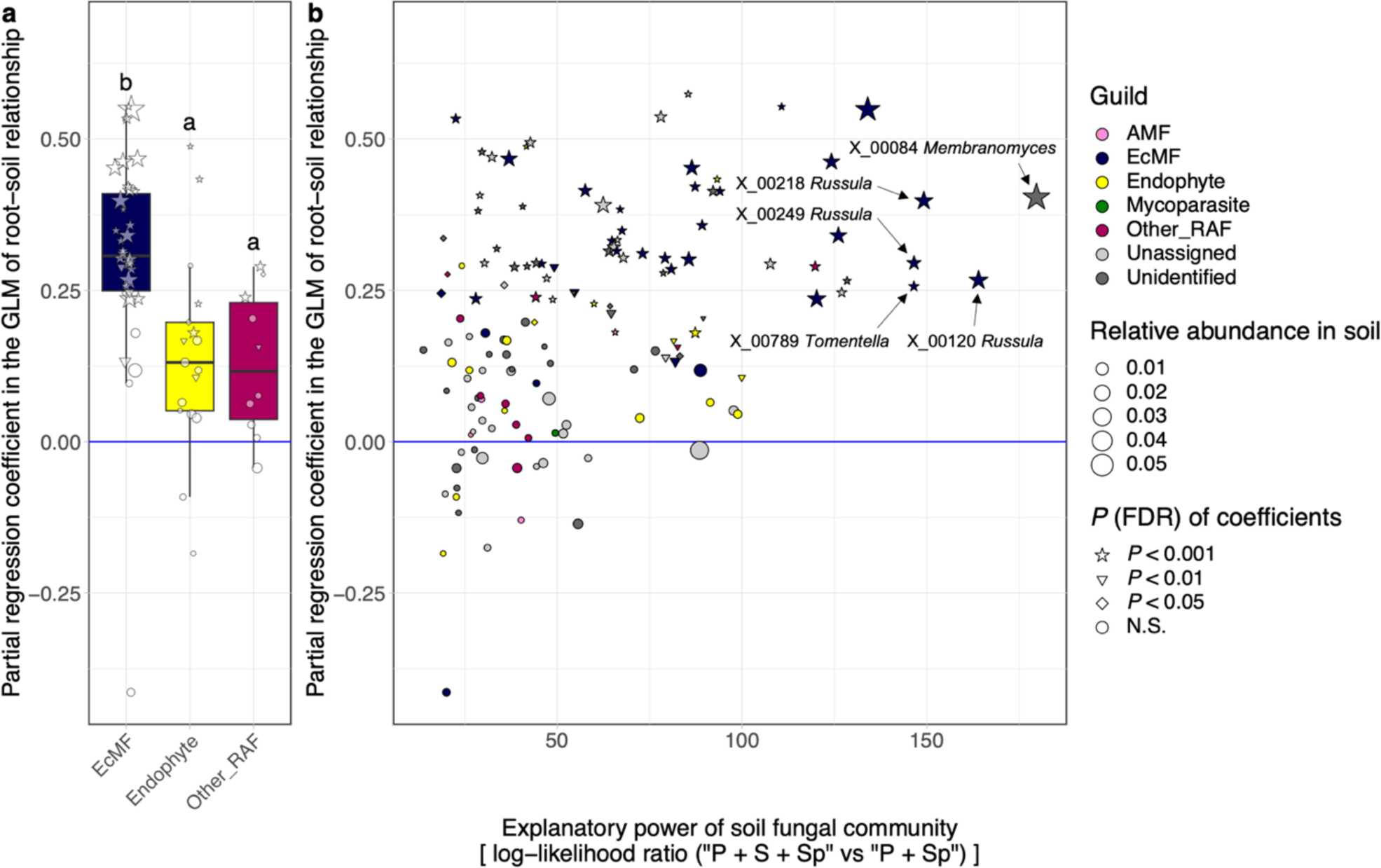
Relationship between fungal OTU occurrence in root and relative abundance in the soil. For each fungal OTU (defined at the 97% cutoff sequence similarity), a generalized liner model (GLM) of occurrence in root samples (the presence/absence of a target OTU in root-tip samples collected at each of the 124 sampling sites) was constructed by setting the relative abundance of the OTU in soil samples (the proportion of sequence reads at each of the 124 sites) as the explanatory variable. The standardized regression coefficients representing the influence of background soil population structure on endosphere fungal distribution were then obtained for respective fungal OTUs. (a) Variation in the partial regression coefficients among fungal functional guilds. The guild of arbuscular mycorrhizal fungi is not shown due to the small number of OTUs examined in the analysis. Difference in the log-likelihood ratios among fungal functional guilds were examined with Steel-Dwass test (significant differences were indicated by different letters). Symbols represent statistical significance of the partial regression coefficients in the GLMs. Symbol size indicates the average relative abundance (proportion of sequence reads) of each fungal OTU across 124 soil samples. (b) Relationship between the partial regression coefficients and jSDM results. The log-likelihood ratios of the two jSDM models differing in the presence/absence of background soil community structure (“P + S + Sp” vs. “P + Sp”; see Table 1) are shown along the horizontal axis. The OTU ID and taxonomy of the top-5 fungal OTUs with the greatest log-likelihood ratios are shown.

### Spatial autocorrelation

Mantel’s correlogram analysis detected statistically significant spatial autocorrelation for the entire root-associated fungal community within ca. 10 m, but the degree of spatial autocorrelation was weak (Mantel’s *r* = 0.024, *P* < 0.001; Figure 7). In the supplemental analysis conducted for each fungal functional guild, the root fungal community structure of endophytic, mycoparasitic, and unclassified root-associated fungi showed weak spatial autocorrelation (Mantel’s *r* < 0.05) in all the distance class (Figure 7). On the other hand, ectomycorrhizal and arbuscular mycorrhizal fungi had statistically significant and relatively strong spatial autocorrelations within ca. 10 m (Mantel’s *r* = 0.08 for ectomycorrhizal fungi and 0.069 for arbuscular mycorrhizal fungi; Figure 7).

**Figure 7:**
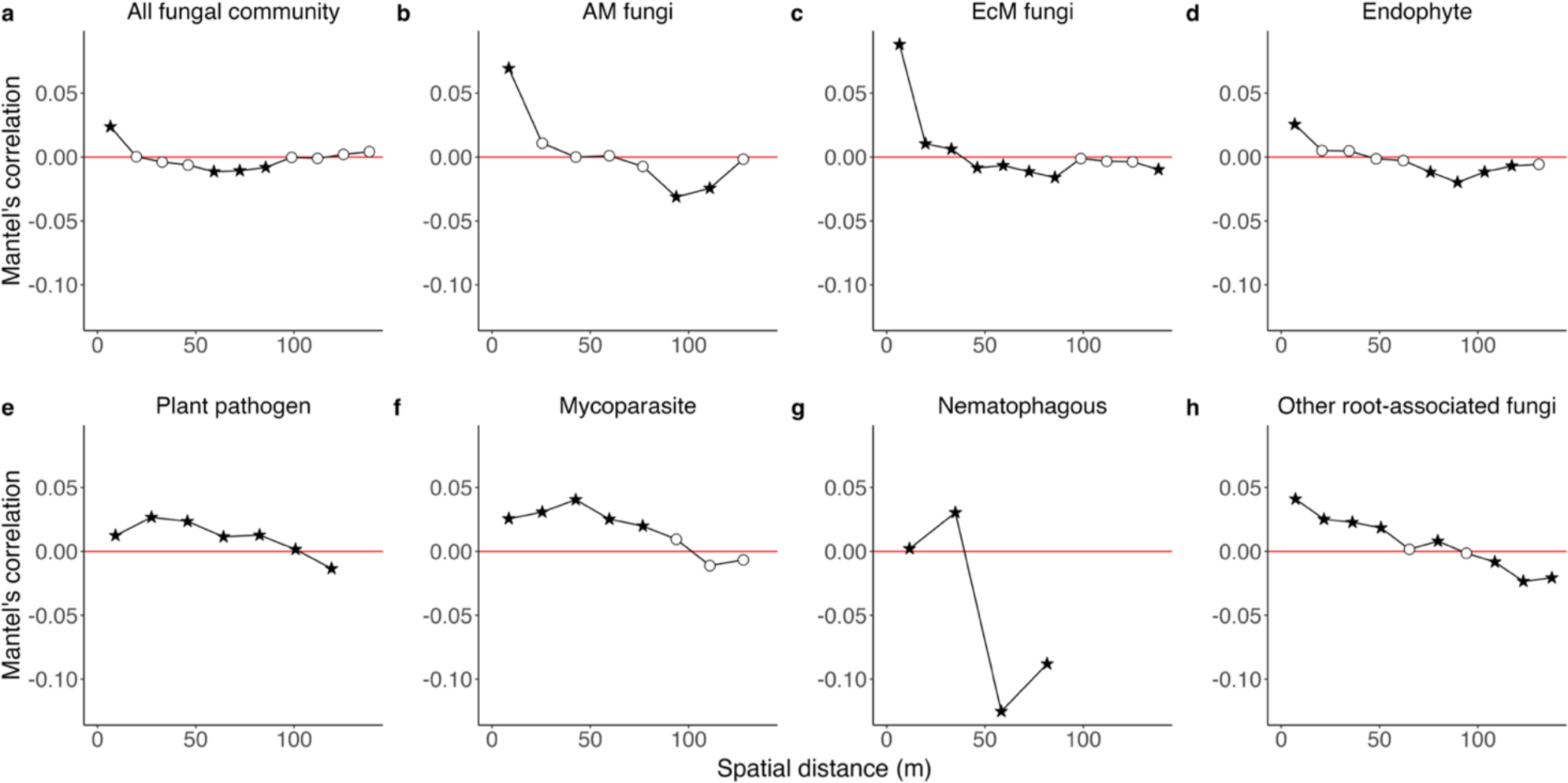
Scales of spatially-autocorrelated community structure. (a) The magnitude of the spatial autocorrelation of fungal community structure is shown along the axis of spatial distance based on a Mantel’s correlogram analysis. Specifically, Mantel’s correlation between Jaccard dissimilarity (*β*-diversity) and geographic distance was examined for each distance class. A filled and star shaped symbol represented statistically significant correlation in the Mantel’s test [*P* (FDR) < 0.05]. In addition to the analysis including all functional guilds of fungi, results on the sub-datasets including only arbuscular mycorrhizal fungi (b), ectomycorrhizal fungi (c), endophytic fungi (d), plant-pathogenic fungi (e), mycoparasitic fungi (f), nematophagous fungi (g), or other root-associated fungi (h) are presented.

### Fungus–fungus network analysis

In the network analysis, a SPIEC-EASI model with six latent variables was selected based on BIC (Figure 8; see also Figure S9). Within the inferred network of fungus–fungus associations, positive associations were inferred in 305 pairs of fungal OTUs: e.g., *Membranomyces* sp. (X_00084) – *Hyaloscypha bicolor* (X_00361), *Lactarius tabidus* (X_00226) *–* Hyaloscyphaceae sp. (X_00149), *Leptodontidium* sp. (X_00452) *–* Hyaloscyphaceae sp. (X_00149), *Lactarius tabidus* (X_00226) – *Russula emetica* (X_00700), *Tarzetta catinus* (X_00030) – Thelephoraceae sp. (X_00289), *Russula turci* (X_00440) – *Tomentella* sp. (X_00693) and *Russula cascadensis* (X_00894) *– Clavulina amethystine* (X_18065) pairs (Table 2). Negative associations were also inferred in 19 pairs of fungal OTUs: e.g., *Russula* sp. (X_00282) – *Russula* sp. (X_00120), *Russula velenovskyi* (X_00265) – *Phialocephala fortinii* (X_00176), *Tarzetta catinus* (X_00030) – *Phialocephala fortinii* (X_00176), *Cladophialophora* sp. (X_00235) – *Russula vesca* (X_00218), Hyaloscyphaceae sp. (X_00149) – *Phialocephala fortinii* (X_00176) and *Leptodontidium* sp. (X_00452) – *Phialocephala fortinii* (X_00176), although these associations were relatively weak (Table 3). The network of positive fungus–fungus associations was compartmentalized into 13 modules, each of which consisted of sets of fungi densely linked with each other (Figure 8a,b). Among the modules, nine were consisted of multiple functional guilds of fungi; in particular, ectomycorrhizal and endomycorrhizal fungi co-occurred in six modules (Table 5).

**Figure 8:**
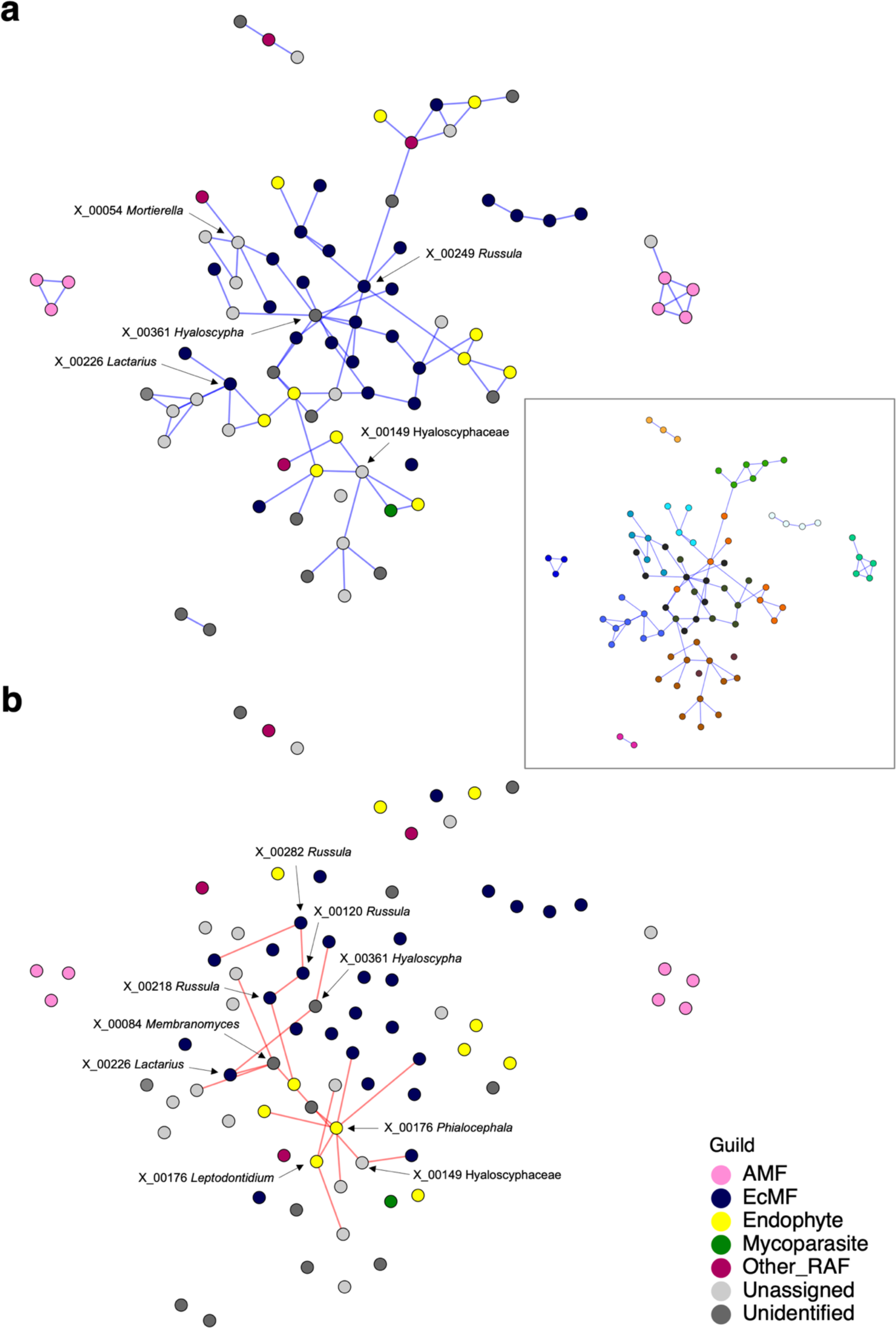
Networks of fungus–fungus associations. Potential interactions between fungal OTUs (defined at the 97% cutoff sequence similarity) were inferred based on the SLR method of SPIEC-EASI. Potential effects of environmental preferences shared between fungal OTUs were controlled in the BIC-selected best model, which included six latent variables. The inferred network architecture was shown separately for positive (a) and negative (b) associations between fungal OTUs. For the positive association network (a), modules of densely associated sets of fungal OTUs were inferred with the Louvain algorithm (see the box in the right side and Table S10). Node colors represent fungal functional guilds. The OTU ID and taxonomy of the OTUs with ≥ 5 positive association links or ≥ 2 negative association links are presented.

**Table 3:**
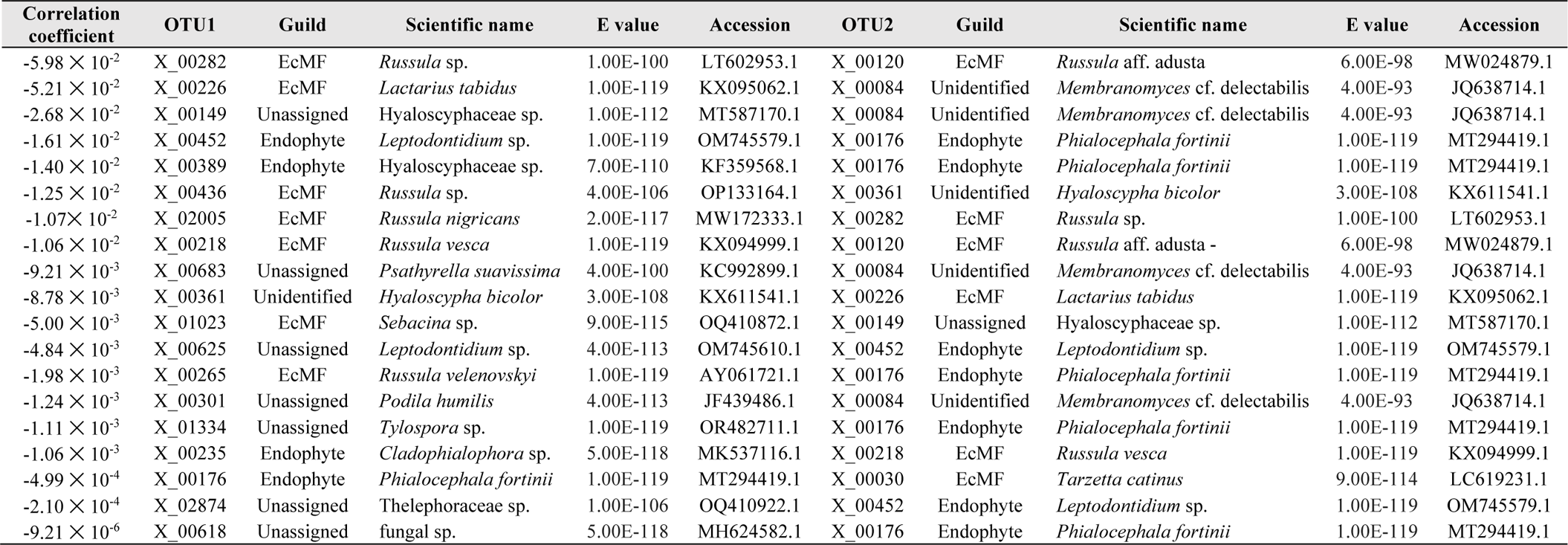
Within-root negative associations between fungi. With the SLR method of SPIEC-EASI analysis, fungus–fungus associations were inferred by controlling the potential effects of shared environmental preference (Figure 8b). This list includes the top-20 fungal OTU pairs in which negative associations were suggested based on the analysis. The OTUs were defined at the 97% cutoff similarity. For each fungal OTU pair, the correlation coefficient estimated with SPIEC-EASI was shown with the OTU ID, the functional guild automatically assigned by the Fungaltraits database, and BLAST top-hit results (the scientific name of the matched sequence, E-value, and NCBI accession number) of each fungus.

We then compared the results on fungus–fungus association patterns between SPIEC-EASI and jSDM. Across fungal OTUs, the degree centrality in the positive and negative association network inferred in the SPIEC-EASI analysis was significantly correlated with log-likelihood ratios comparing the full model (P + S + Sp + Cov) and the model from which fungus–fungus covariance was removed (P + S + Sp) in jSDM (Positive: *τ* = 0.283, *P* < 0.001, Negative: *τ* = 0.256, *P* < 0.001; Figure 9). The analysis highlighted some endophytic fungi whose distribution patterns were inferred to be more dependent on fungus–fungus associations than those of other fungi (e.g., Hyaloscyphaceae sp. [X_00149], *Phialocephala fortinii* [X_00176] and *Leptodontidium* sp. [X_00452]) (Table S11, Table S12, Figure 9).

**Figure 9:**
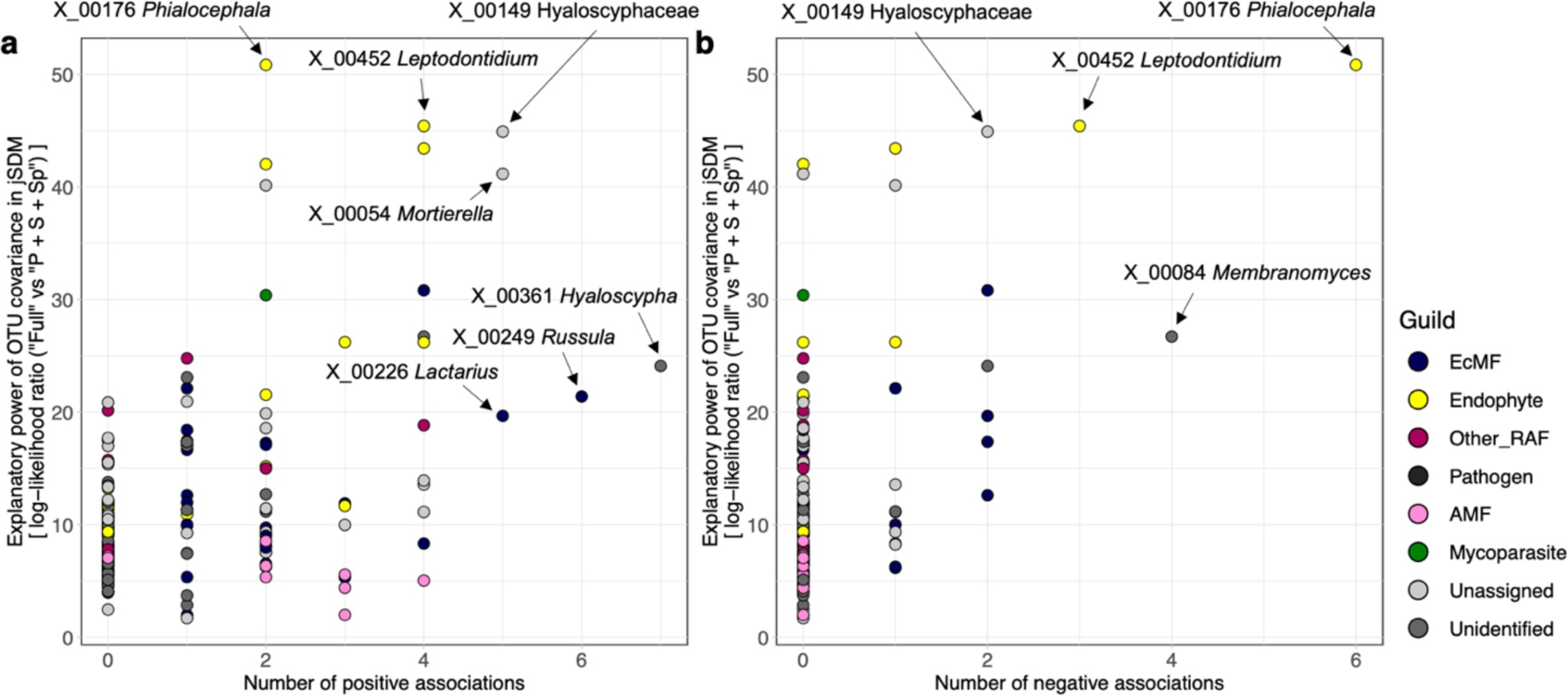
Fungal OTUs (defined at the 97% cutoff sequence similarity) with the strongest signs of species interactions. Fungi potentially playing key roles within the community were explored based on both the jSDM (Figure 2) and SPIEC-EASI (Figure 8) frameworks. Based on the jSDM, the log-likelihood ratios of the two models differing in the presence/absence of fungus–fungus covariance [“Full (P + S + SP + Cov)” vs. “P + S + SP”; see Table 1] are shown for respective fungal OTUs along the vertical axis. Meanwhile, the number of positive (a) or negative (b) associations (i.e., degree centrality) within the SPIEC-EASI networks is shown along the horizontal axis. The degree centrality was significantly correlated with the log-likelihood ratio of the jSDM for both analyses of positive (Kendal’s correlation coefficient = 0.283, *P* < 0.001) and negative (Kendal’s correlation coefficient = 0.256, *P* < 0.001) associations. Colors represent fungal functional guilds. The OTU ID and taxonomy of the OTUs showing the highest levels of log-likelihood ratios or network degree centralities are presented.

Qualitatively similar results were obtained for the analysis at the 93% OTU threshold in fugal aggregation patterns, although the number of fungal OTUs analyzed was reduced in the supplemental analysis (Figure S10, Figure S11). Few fungus–fungus associations were inferred in the analysis of segregated patterns at the 93% cutoff similarity (Figure S10, Figure S11).

## DISCUSSION

By the fungal community profiling of 1,615 root samples collected in a cool-temperate forest, this study examined how multiple ecological factors collectively organize the community structure of fungi in root systems. Our field sampling across more than 100 positions allowed us to apply spatial statistical analyses based on jSDM, by which host plant identity, background soil fungal community structure, spatial autocorrelation, and potential fungus–fungus associations were simultaneously considered. The general statistical platform provided a novel opportunity for comparing basic ecological processes among major root-associated fungal guilds. In particular, the fact that ectomycorrhizal and endophytic fungal guilds systematically differ in the balance of basic assembly factors is expected to reorganize our understanding of forest-scale dynamics of plant–fungus symbioses as discussed below.

### Multiple assembly factors

Concomitantly evaluating the impacts of multiple ecological factors on species distributions is a prerequisite for comprehensively understanding the assembly processes of fungi associated with plant roots. Our statistical analysis based on jSDM revealed that filtering by host plant species, background soil fungal community structure, spatial autocorrelation, and fungus–fungus associations (covariance) collectively determine the fine-scale community structure of root-associated fungi (Figure 2; Table S3-6). Among the factors, filtering by host plants has been regarded as stongly deterministic factor, while background community structure and spatial autocorrelation are influenced by stochasticity in soil ecosystem processes (Fierer, 2017; Munoz-Ucros et al, 2021). Thus, by extending discussion in previous studies examining roles of single ecological factors (e.g. Goldmann et al., 2016; Li et al., 2021; Shakoor et al., 2018; Taniguchi et al., 2023), this study indicates how deterministic and stochastic processes collectively operate in the assembly of root-associated fungi.

When combinations of ecological factors were examined in the framework of jSDM, partial models involving fungus–fungus covariance and background soil fungal community structure best explained the distribution of root-associated fungi (Figure 2). This result does not necessarily mean that direct facilitative and/or competitive interactions between fungal species within root systems or species interactions at the endosphere–rhizosphere interface (Kennedy, 2010; Martínez-Medina et al., 2011; Raaijmakers et al., 2009; Werner & Kiers, 2015) are the major drivers of fungal community assembly. Nonetheless, the statistical analysis based on jSDM illuminates the importance of evaluating potential roles of fungus–fungus associations in understanding the assembly of root-associated fungal communities. Regarding fungus–fungus covariance, not only direct fungus–fungus interactions but also sharing of environmental preferences between fungal species can underly the observed patterns in fungal distributions. Because our jSDM included a limited number of environmental factors (e.g., host plant identity), large parts of variation could remain unexplained by the explicit variables. Thus, in the jSDM framework, such variation in observed patterns would be attributed to fungus–fungus covariance, which implicitly represent not only direct species interactions but also shared niche preference. For further understanding how direct fungus–fungus interactions organize root-associated fungal community assembly, the jSDM approach needs to be expanded by incorporating more background environmental variables such as phosphorus/nitrogen concentrations in the soil and host-plant physiochemical states (Gong et al., 2022; Nguyen et al., 2020; Pang et al., 2021).

### Fungal guilds and assembly rules

Although relative contributions of community ecological factors should be carefully discussed, jSDM allows us to examine whether basic assembly rules could differ among fungal functional guilds. By making the most of the flexibility of likelihood-based analyses in jSDM, we were able to compare the effects of each ecological factor on species distribution patterns among fungal OTUs. We then found that the magnitude of impacts by each factor differed greatly among fungal guilds (Figure 3). For example, host plant identity showed less impacts on the distribution of ectomycorrhizal fungi than on the distribution of other fungi in the cool-temperate forest dominated by ectomycorrhizal plants (Figure 3a). In contrast, ectomycorrhizal fungal distribution was more influenced by background fungal community structure in the soil than root-endophytic, arbuscular mycorrhizal, and unclassified root-associated fungi (Figure 3b). Likewise, the patterns observed for ectomycorrhizal fungi reflected spatial autocorrelations more intensively than endophytic and unclassified root-associated fungi (Figure 3c). The large contributions of the two ecological factors are possibly explained by the spatial expansion strategies of ectomycorrhizal fungi. Specifically, as ectomycorrhizal fungi are known to form intensive below-ground networks of extraradical hyphae (Selosse et al., 2006, Smith and Read, 2008, Klein et al., 2016), their spatial patterns in plant roots are described as spatially structured colonization from such hyphal networks.

In terms of variation in ecological properties among fungal guilds, the jSDM and a series of supplementary analyses illuminated the uniqueness of root-endophytic fungi. Albeit statistically non-significant, relationship between root-associated and background soil community structure was weaker for endophytic fungi than for ectomycorrhizal fungi (Figure 3b). A GLM-based analysis further indicated that high occurrence in root samples was strongly associated with high abundance in background soil for ectomycorrhizal fungi but not for endophytic fungi (Figure 6). Our data also suggested that some endophytic fungi occurred very frequently in root samples (Figure 5a) and their relative abundance can be exceptionally high in the roots (Figure 5b). Moreover, signs of spatially autocorrelated distribution were weaker in endophytic fungi than in mycorrhizal fungi (Figures 3c and 7).

These results provide an opportunity for inferring the ecology of endophytic fungi in below-ground ecosystems. Many of root-endophytic fungi, especially those called dark septate endophytes (Grünig et al., 2011; Jumpponen, 2001; Jumpponen & Trappe, 1998; Rodriguez et al., 2009), has been known to support the growth and survival of their host plants (Akhtar et al., 2022; Newsham 2011; Mandyam & Jumpponen, 2005). Thus, in terms of overall effects on host plants, root-endophytic and ectomycorrhizal fungi play similar roles in forest ecosystems. However, the statistical results outlined above indicate that the two fungal guilds differ in the basic nature of symbiosis with plants. The relatively weak sign of spatial structuring suggests that root-endophytic fungi do not construct rigorous hyphal networks such as those formed by ectomycorrhizal fungi. Thus, while ectomycorrhizal fungi transport soil phosphorus and nitrogen to their host plants through extensive extraradical hyphae (Smith and Read, 2008), root-endophytic fungi possibly contribute to plants’ physiology through different mechanisms. Interestingly, a meta-analysis of physiological experiments on plant–fungal interactions has suggested that root-endophytic fungi mineralize organic nitrogen-containing compounds in the rhizosphere, thereby provisioning plants with inorganic forms of nitrogen (Newsham 2011). In addition, some dark septate endophytes could have abilities to decompose complex polymeric forms of organic nutrients including diverse organic phosphorus compounds (Della Monica et al., 2015; Surono & Narisawa, 2017) using their metabolic toolboxes (Caldwell et al., 2000; Knapp et al., 2018; Knapp & Kovács, 2016; Schlegel et al., 2016). Therefore, ectomycorrhizal fungi, which transport nitrogen/phosphorus from distal parts, and endophytic fungi, which support plants’ nitrogen acquisition from diverse and complex substrates around roots, possibly play complementary functions in below-ground nutrient cycles.

The difference in the level of spatial structuring highlights potential difference in dispersal strategies between ectomycorrhizal and root-endophytic fungi. Ectomycorrhizal fungi disperse their spores by wind from above-ground fruiting bodies. In addition, they rigorously extend their extraradical hyphae in the soil (Smith and Read, 2008) as mentioned above. In contrast, little information is available for the dispersal strategies of root endophytic fungi. Because most endophytic fungi are known exclusively as anamorphs (i.e., asexual stages) (Grünig et al., 2008; Hambleton & Sigler, 2005; Jumpponen & Trappe, 1998), they possibly lack mechanisms of wind dispersal by macroscopic fruiting bodies, resulting in the decay of spatial autocorrelation in community structure within short distance (Figure 7d). Nonetheless, some root-endophytic fungal OTUs occurred at surprisingly high frequencies in root samples (Figure 5a). Given that background soil community structure had weaker impacts on the distribution patterns of root-endophytic fungi than on those of ectomycorrhizal fungi (Figure 3b), the exceptionally high occurrence of endophytic fungi in plant root systems is enigmatic. Although population genetic analyses have begun to provide the evidences of gene flows between local populations of root-endophytic fungi (Nakamura et al., 2019), dispersal mechanisms of most endophytic fungi, including the most prevalent lineage of dark septate endophytes (i.e., *Phialocephala fortinii s.l.–Acephala19pplanatea* species complex), have remained to be explored (Grünig et al., 2008; Hambleton & Sigler, 2005; Jumpponen & Trappe, 1998; Sieber, 2002). An attractive but poorly explored hypothesis is that conidia or mycelia of root-endophytic fungi are efficiently dispersed by specific animal vectors (e.g., mites, springtails, earthworm and rodent) that move between rhizosphere patches within forests (Fracchia et al., 2011).

Overall, statistical analyses targeting the entire fungal taxa can provide a basis for comparing basic ecology among fungal functional guilds, deepening our understanding of below-ground ecosystem processes. In particular, the fact that mycorrhizal and root-endophytic fungi fundamentally differ in the level of host preference and spatial structuring patterns gives essential viewpoints for considering the dynamics of seedling establishment in forest ecosystems. By incorporating accumulated knowledge of mycorrhizal symbiosis, analyses of the entire plant–fungus associations will help us understand how feedback between plant and fungal community processes (i.e., plant–soil feedback) is organized in terrestrial ecosystems.

### Filtering by host plants

Among the ecological factors/processes examined in jSDM, filtering by host plants has been investigated the most intensively in previous studies (Ishida et al., 2007; Kernaghan & Patriquin, 2011; Molina and Trappe, 1982; Põlme et al., 2018). In our analysis, filtering by host plant identity was inferred to be stronger for arbuscular mycorrhizal fungi than for ectomycorrhizal fungi (Figure 3a). This result is seemingly inconsistent with the general belief that arbuscular mycorrhizal fungi have greater host ranges than ectomycorrhizal fungi (Brundrett & Tedersoo, 2018). The point is that having a broad list of potential host plant taxa does not guarantee that a focal fungal species can interact indistinguishably with diverse plant species in a local ecosystem. In other words, an arbuscular mycorrhizal fungal species can interact with a limited repertoire of plant species in a forest dominated by ectomycorrhizal plant species. Thus, it is crucial to acknowledge that statistical analyses in real ecosystems help us clarify patterns of “realized” plant–fungus associations irrespective of *a priori* knowledge of “fundamental” host ranges of fungi.

A randomization analysis of fungal OTU-level preference further provided insights into within-guild variation in host preference. Several arbuscular mycorrhizal fungi, for example, showed strong preferences for specific arbuscular mycorrhizal plants within the forest (e.g., OTUs with significant preferences on *Toxicodendron*, *Juglans*, or *Acer*; Figure 4). This result highlights the previous findings that DNA metabarcoding-based analyses can uncover overlooked specificity between arbuscular mycorrhizal fungal species and host plant taxa (Öpik et al., 2009; Yang et al., 2012). In contrast, all ectomycorrhizal fungi, but some *Tomentella* and *Russula* OTUs, did not show statistically significant preferences for hosts within the cool-temperate forest dominated by ectomycorrhizal plant species (Figure 4). This pattern is consistent with the previous observations that ectomycorrhizal fungi are often detected from non-ectomycorrhizal plant species (Maciá-Vicente & Popa, 2022; Schneider-Maunoury et al., 2020; Toju & Sato, 2018). Intriguingly, some ectomycorrhizal fungi have been known to damage the root tissues of arbuscular mycorrhizal plant species, thereby increasing competitive advantage of their host ectomycorrhizal plant species (Plattner & Hall, 1995; Taschen et al., 2020).

In the forest, root-endophytic fungi exhibited moderate levels of host preferences when compared to other root-associated fungal guilds (Figure 3a). For example, *Rhizodermea* and *Cladophialophora*, which were commonly detected from co-occurring arbuscular mycorrhizal and ectomycorrhizal plant species in a previous study of a warm-temperate forest (Toju & Sato, 2018), displayed significant but weak signs of host preferences (Figure 4). This result on the presence of endophytic fungi with relatively weak host preferences coincides with the increasing reports of endophytic fungi with broad potential host ranges (Jumpponen & Trappe, 1998; Terhonen et al., 2019). In contrast, *Pezicula*, which has been reported to appear more frequently in arbuscular mycorrhizal plant roots than in co-occurring ectomycorrhizal plant roots (Toju & Sato, 2018), showed a statistically significant preference for an arbuscular mycorrhizal plant (*Acer*) in the OTU-level analysis (Figure 4). An OTU belonging to the genus *Phialocephala*, a well-known lineage of dark septate endophytes (Grünig et al., 2008, Jumpponen & Trappe, 1998), showed a high host preference as well (Figure 4). Albeit informative, these observed patterns may not be attributed exclusively to filtering by plants. Tripartite interactions involving not only endophytic fungi and host plants but also fungi in other guilds (Kennedy et al., 2009; Reininger & Sieber, 2012, 2013) can be strong determinants of the realized host–symbiont associations as discussed below.

### Exploration of key interactions and species

The framework of jSDM provides an opportunity for inferring potential contributions of symbiont–symbiont interactions in the assembly of root-associated fungi. However, in the jSDM approach, estimated fungus– fungus covariance can result not only from the effects of direct fungus–fungus interactions but also from those of environmental preferences (niches) shared between fungal species. Therefore, we took an alternative statistical approach (SPIEC-EASI), by which impacts of unobserved environmental conditions can be controlled in the inference of species interactions (Kurtz et al., 2019). The BIC-selected model that included six latent environmental variables then highlighted potential facilitative and competitive interactions between pairs of fungal OTUs (Figure 8). The strongest facilitative interaction was estimated between a *Membranomyces* fungus and a *Hyaloscypha* fungus (Table 2). Given that *Membranomyces* is a possibly ectomycorrhizal genus (Uehling et al., 2012) and that *Hyaloscypha* is20pplanate20icallyy allied to root endophytes and a well-known ericoid mycorrhizal fungus *Rhizoscyphus ericae* (Fehrer et al., 2019), this result confirms the presence of tight positive interactions between ectomycorrhizal and root-endophytic fungi (Berthelot et al., 2019). Such positive interactions between endophytic and ectomycorrhizal fungi were inferred as well in a set of Hyaloscyphaceae sp. and an ectomycorrhizal *Lactarius* OTU (Table 2). In addition, our results also suggested facilitative interactions between possibly endophytic fungi (e.g., *Leptodontidium*–Hyaloschyphaceae) and those between ectomycorrhizal fungi (e.g., *Russula*–*Lactarius*, *Tarzetta*–*Thelephoraceae*, *Tomentella*–*Russula*, and *Clavulina*–*Russula* pairs; Table 2). The inference of positive interactions between ectomycorrhizal fungi are of particular interest because ectomycorrhizal fungi have been known to compete with each other within host root systems (Kennedy, 2010; Kennedy et al., 2009; Pickles et al., 2012). In fact, negative interactions were observed in some pairs of ectomycorrhizal fungi (e.g., *Russula*–*Russula* pairs) as well as in pairs of ectomycorrhizal fungi and endophytic fungi (e.g., *Russula*–*Phialocephala*, *Cladophialophora*–*Russula*, and *Tarzetta*–*Phialocephala* pairs) and those of endophytic fungi (e.g., *Leptodontidium*–*Phialocephala* and Hyaloscyphaceae–*Phialocephala* pairs; Table 3).

These positive and negative interactions between fungi are expected to drive the assembly of root-associated fungal communities through historically contingent processes (Fukami, 2015; Kennedy, 2010; Sikes et al., 2016). The colonization of an endophytic or mycorrhizal fungus may change the micro-environment within host-plant root systems (i.e., niche construction; Muthukumar & Sulaiman, 2021; Sasan & Bidochka, 2012; Berta et al., 1993), thereby promoting the secondary colorization of specific sets of compatible fungi (Toju et al., 2020; Toju et al., 2018). Meanwhile, the presence of a fungal species in roots can work as a barrier to following colonizers through the preemption of space/resources or specific blocking mechanisms (Kennedy, 2010). Previous experimental studies on mycorrhizal fungi have shown that early colonizers can prevent the followers’ entry into plant root systems and that the order of arrivals, rather than individual species’ competitive abilities, would be the essential factor of assembly patterns (Kennedy et al., 2009, Werner and Kiers, 2015). Such priority effects (Fukami, 2015) may be stronger in ectomycorrhizal fungal communities (Kennedy, 2010) than in arbuscular mycorrhizal fungal communities (Werner and Kiers, 2015) potentially because ectomycorrhizal fungi can defend their territories by forming dense hyphal structures (mantles) surrounding host roots (Smith and Read, 2008). Potentially due to the defensive abilities of ectomycorrhizal fungi, the inferred negative interations between ectomycorrhizal fungi and fungi in other ecological guilds (Table 3) may be asymmetric (Sun et al., 2023). Indeed, an experimental study has shown that ectomycorrhizal fungi can impose asymmetric suppressive impacts on root-endophytic fungi (Reininger & Sieber, 2012).

When the information of potential pairwise interactions was integrated, some potentially endophytic fungi (e.g., Hyaloscyphaceae, *Leptodontidium*, and *Phialocephala*) were located at the core positions within each of the positive and negative interaction networks (Figures 8-9). With respect to positive symbiont– symbiont association networks, such hub species may play pivotal roles in promoting the coexistence of functionally variable fungi within plant root systems (Abrego et al., 2020; Toju et al., 2018). Hubs within negative interaction networks are also expected to control the entire structure of root-associated fungal communities (Netherway et al., 2024; Toju et al., 2018). Thus, in addition to ectomycorrhizal fungi, which possess clear physical defensive mechanisms, root-endophytic fungi could be strong determinants of the entire patterns of below-ground plant–fungus symbioses in cool-temperate forests, driving plant–soil feedback processes. The ecological and physiological mechanisms by which root-endophytic fungi interact with many other fungi deserve future intensive investigations.

### Caveats and methodological challenges

Although our forest-wide analyses provide a novel opportunity for evaluating how multiple ecological factors/processes drive the entire structure of complex associations between plants and their numerous symbionts, the results need to be interpreted with caution in light of potential pitfalls and methodological limitations. First, because this study was based on data from a single time point in a single forest dominated by ectomycorrhizal trees, we are unable to conclude that similar ecological patterns (e.g., variation in the balance of assembly factors among fungal guilds) can be observed in other forest ecosystems. Comparative research across a wide range of latitudes from the tropics to boreal regions will help us understand how forests with different dominant mycorrhizal types vary in their plant–soil feedback processes (Bennett et al., 2017; Kadowaki et al., 2018). Second, it should be kept in mind that DNA-based data do not provide direct evidences of interactions between fungal species. Even with analyses incorporating latent environmental variables, effects of direct species interactions would not be completely separated from those of shared environmental preferences (Blanchet et al., 2020; Hirano & Takemoto, 2019; Kurtz et al., 2015, 2019). Therefore, microscopic observation of hyphal interactions inside fine roots, genes expression analyses of fungal physiological states in host root tissues, and experimental co-inoculation studies are necessary for deepening our knowledge of fungus–fungus interactions. Third, the jSDM approach need to be extended to partition variances in distribution patterns more comprehensively. With the current framework, the covariance term of species distributions needs to be carefully interpreted because it can contain impacts of ecological factors other than direct species interactions (Blanchet et al., 2020; Leibold et al., 2022). The incorporation of latent variables, for example, may broaden the application of jSDM to ecological community datasets with limited information of background environmental conditions.

## CONCLUSIONS

By applying emerging statistical frameworks to massive datasets of root-associated fungal communities, we here examined how multiple ecological factors/processes organize spatial distribution patterns of below-ground plant–fungus symbioses. The results then suggested that fungal community structure in the background soil and fungus–fungus associations within roots, as well as filtering by host plants and spatially autocorrelation in ecological processes, could collectively drive the assembly of root-associated fungi. We also found that the relative importance of the assembly factors/processes could differ systematically between mycorrhizal and root-endophytic fungi. Our analysis further suggested that root-endophytic fungi, whose diversity and physiological functions have been largely unknown, potentially played hidden ecological roles within networks of symbiont–symbiont interactions. All these insights are crucial for understanding how below-ground ecological processes organize plant community regeneration and succession. Towards further integrative knowledge, comparative studies need to be conducted in various types of terrestrial ecosystems. Studies on temperate or tropical forests co-dominated by arbuscular mycorrhizal and ectomycorrhizal plant species (Bennett et al., 2017; Kadowaki et al., 2018; Segnitz et al., 2020; Ushio et al., 2017), for example, will help us understand how root-endophytic fungi buffer or promote positive/negative feedback between below-ground and above-ground community dynamics.

## AUTHOR CONTRIBUTIONS

Mikihito Noguchi and Hirokazu Toju designed the research and collected data. Mikihito Noguchi analyzed the data and wrote the manuscript, advised by Hirokazu Toju.

## ACKNOWLEDGMENTS

We thank Tanaka Kenta for his advice on fieldwork. We are also grateful to Sugadaira Research Station, Mountain Science Center, University of Tsukuba for the permission of the fieldwork. This work was financially supported by JST FOREST (JPMJFR2048), Human Frontier Science Program (RGP0029/2019), and JST CREST (JPMJCR23N5) to H.T. as well as by JSPS Research Fellowships for Young Scientists (JP23KJ1380) to M.N.

## DATA AVAILABILITY STATEMENT

The DNA sequence data have been deposited to the DNA Data Bank of Japan (DDBJ accession: PRJDB17520) [to be released after acceptance of the paper]. The computer codes are available from the GitHub repository (https://github.com/ngchngch/contrasting-but-interdependent-assembly-processes-of-mycorrhizal-and-endophytic-fungi) [to be released after acceptance of the paper].

