## Supplementary Figures and Tables for "Mycorrhizal and endophytic fungi structure contrasting but interdependent assembly processes in forest below-ground symbiosis"

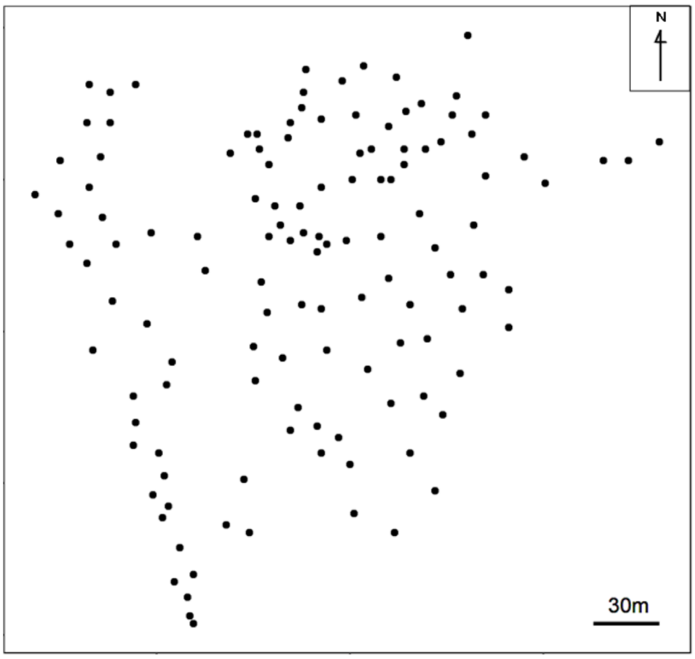


Figure S1：Sampling points. The 126 points from which plant root and soil were sampled are indicated by dots.

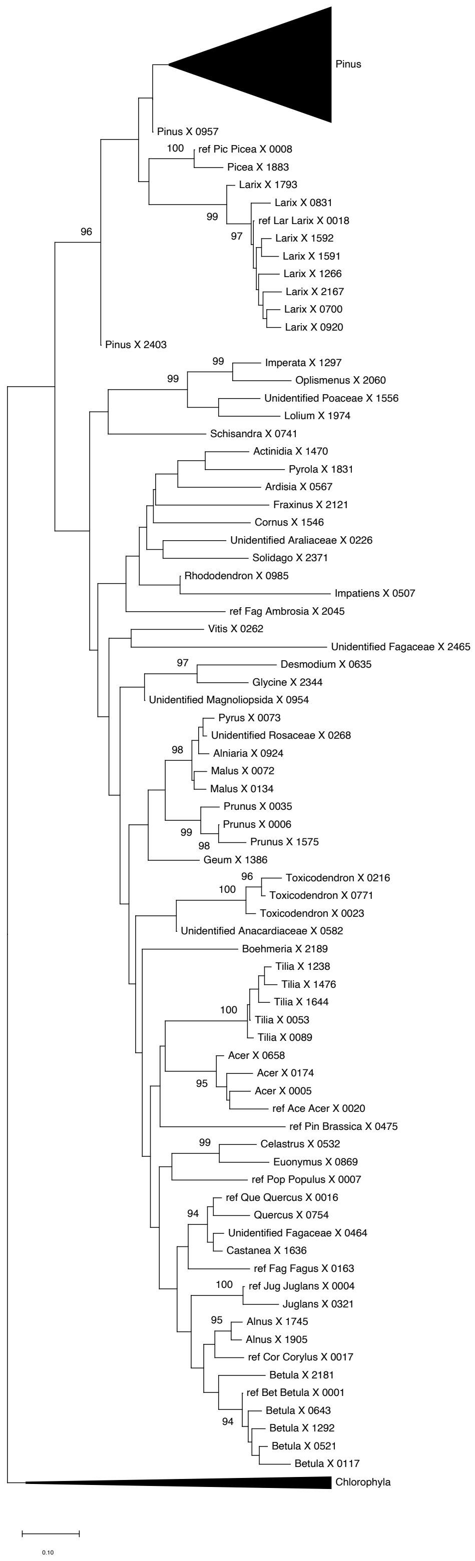


Figure S2: Phylogenetic tree of the host plants. A phylogenetic tree of streptophyta and chlorophyta based on the neighbor joining method using partial sequences of the plant ITS2 region. Sequences of chlorophyta are grouped together as an outgroup ("Chlorophyta"). In addition, because a number of homologous sequences were identified as the genus *Pinus*, they are shown as a group ("Pinus"). For the other sequences, the OTU name and the BLAST taxonomic assignment at the genus level are shown at the terminal of each branch.

Sequences derived from reference plant leaf samples were represented as "ref – [Plant name] – [OTU name]”. Branches with > 90% bootstrap values (1,000 times) are indicated.

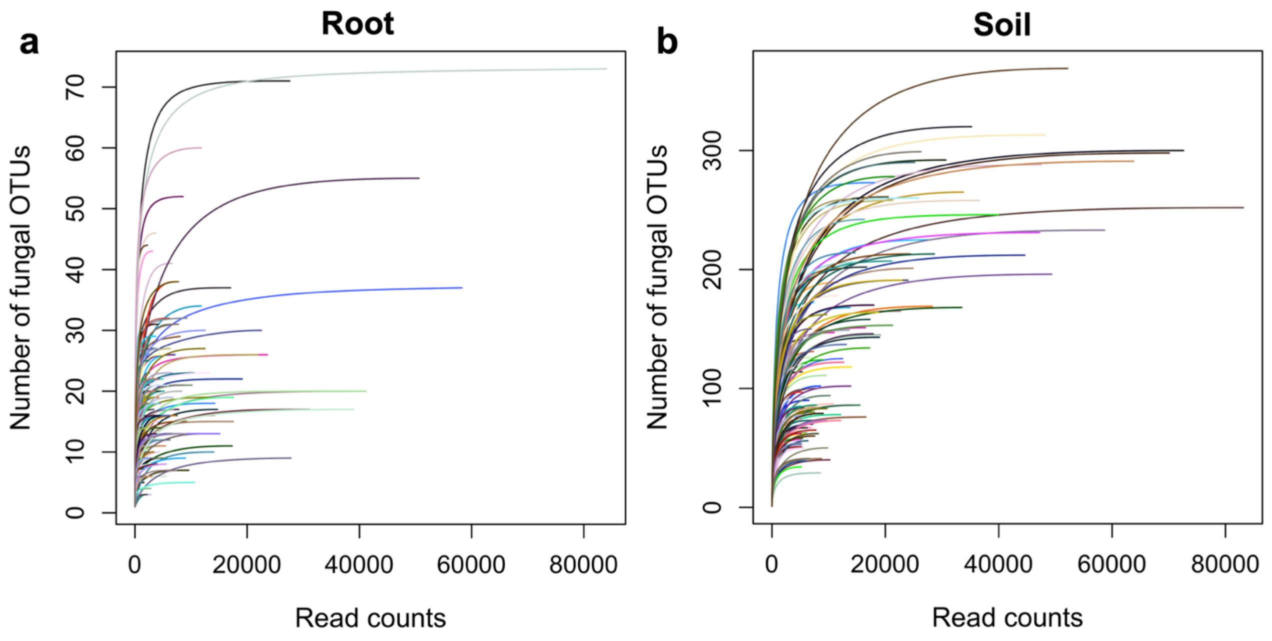


Figure S3: Relationships between the number of sequencing reads and that of fungal OTUs (cutoff sequence similarity = 97%). (a) Number of fungal OTUs detected in plant root samples. For visualization, the relationship between the number of sequencing reads and the number of fungal OTUs detected in 200 randomly selected samples is shown. (b) Number of fungal OTUs detected in soil samples. The relationship between the number of sequencing reads and the number of fungi detected in 124 samples whose reads were successfully obtained.

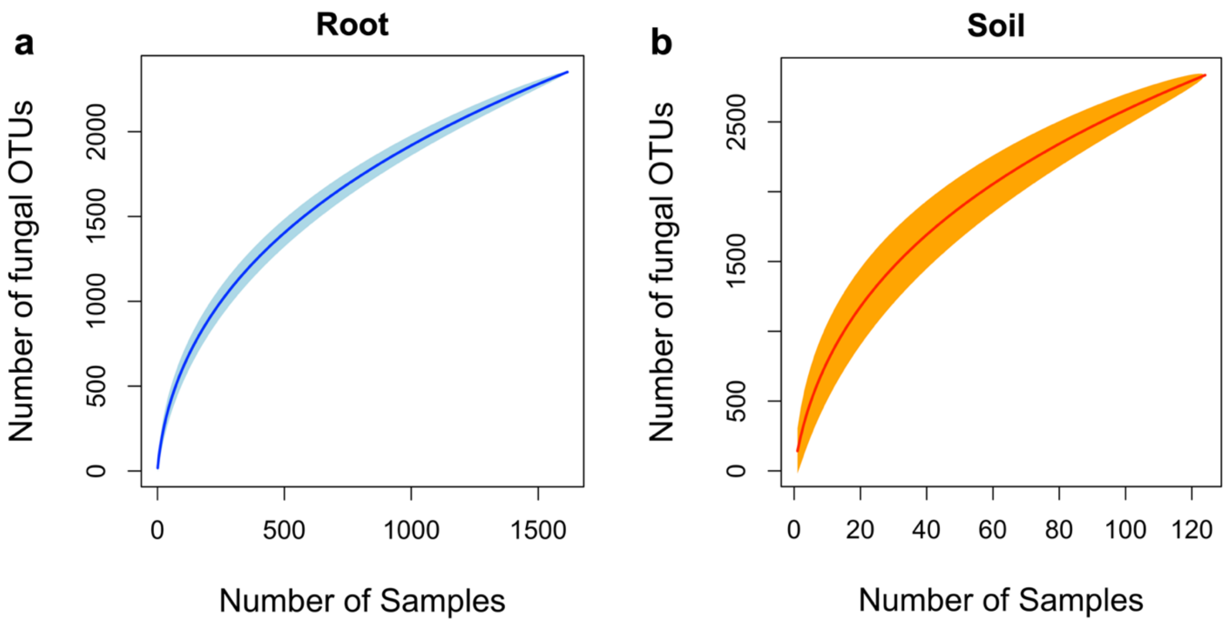


Figure S4: Relationship between the number of root/soil samples and that of fungal OTUs (cutoff sequence similarity = 97%) observed. (a) Number of fungal OTUs in root samples with plant genus identified. In total, sequencing data were successfully obtained from 1,615 samples, and 2,351 OTUs were detected across the samples. (b) Number of fungal OTUs in soil samples. In total, sequencing data were successfully obtained from 124 samples, and 2,834 OTUs were detected across the samples.

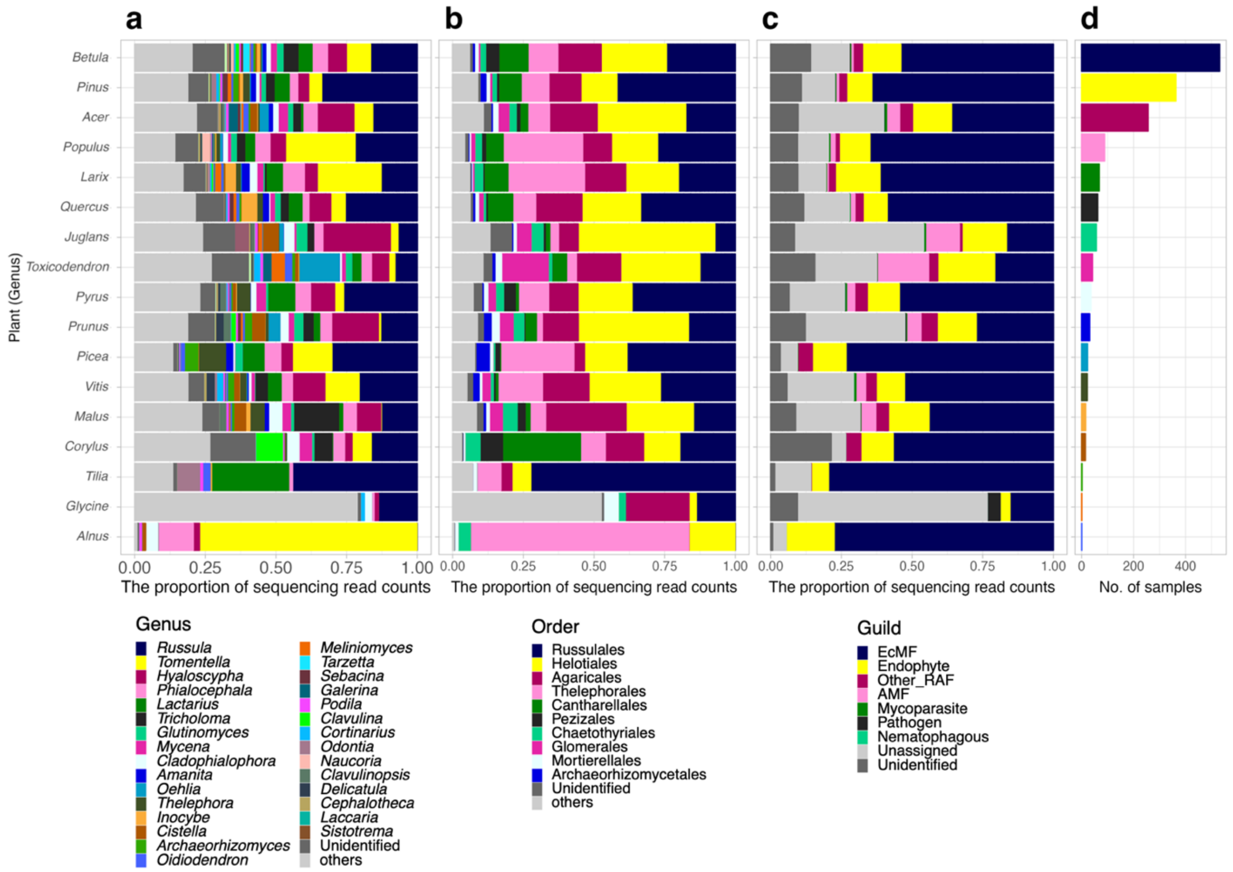


Figure S5: Root fungal community compositions. (a) Genus-level taxonomic compositions. The mean proportions of fungal sequencing reads are shown for each plant genus. (b) Order-level taxonomic compositions. (c) Guild compositions. The guilds (functional groups) of fungal OTUs were inferred with the Fungaltraits database. (d) Number of plant root samples.

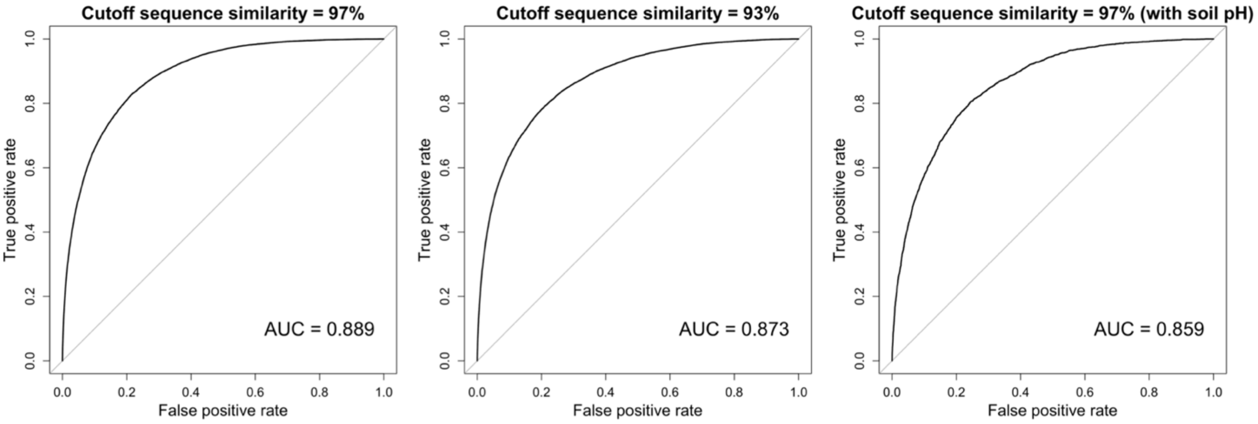


Figure S6: Predictive accuracy of the jSDM models for the distribution of fungal OTU (cutoff sequence similarity = 93%) and fungal OTU (cutoff threshold = 97%) considering the four or five factors. The relationship between the false positive rate and the true positive rate for each model (OTU (93%) and OTU (97%): Model 1 in Table 1, OTU (97%) with soil pH: Model 1 in Table S1) is shown as the prediction accuracy for the distribution of fungal OTU (93%) and fungal OTU (97%). Area under the curve (AUC) is also shown in the figures, respectively.

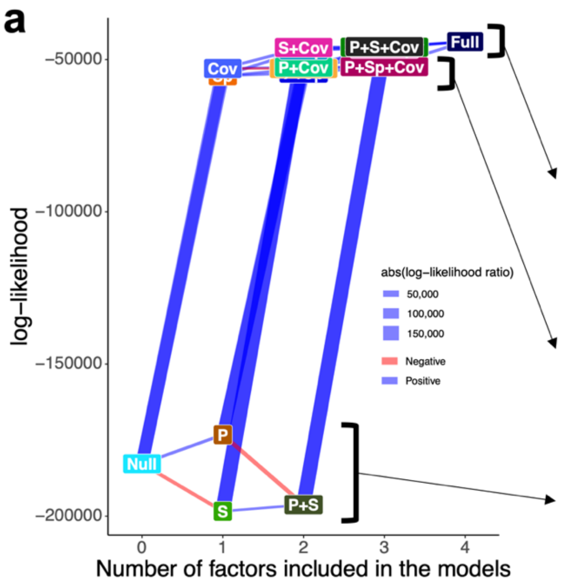


b

| | | | Factors in the models | | | |
|---|---|---|---|---|---|---|
| | Model name | log-likelihood | Host plant | Soil fungal community | Spatial autocorrelation | Inter-OTU covariance |
| | Full ( P + S + Sp + Cov) | $-4.41 \times 10^4$ | ○ | ○ | ○ | ○ |
| | P + S + Cov | $-4.58 \times 10^4$ | ○ | ○ | | ○ |
| | S + Sp + Cov | $-4.59 \times 10^4$ | | ○ | ○ | ○ |
| | S + Cov | $-4.60 \times 10^4$ | | ○ | | ○ |
| | P + S + Sp | $-4.64 \times 10^4$ | ○ | ○ | ○ | |
| | S + Sp | $-4.68 \times 10^4$ | | ○ | ○ | |
| | P + Sp + Cov | $-5.23 \times 10^4$ | ○ | | ○ | ○ |
| | P + Cov | $-5.24 \times 10^4$ | ○ | | | ○ |
| | Sp + Cov | $-5.29 \times 10^4$ | | | ○ | ○ |
| | Cov | $-5.29 \times 10^4$ | | | | ○ |
| | P + Sp | $-5.44 \times 10^4$ | ○ | | ○ | |
| | Sp | $-5.53 \times 10^4$ | | | ○ | |
| | P | $-1.73 \times 10^5$ | ○ | | | |
| | Null | $-1.83 \times 10^5$ | | | | |
| | P + S | $-1.96 \times 10^5$ | ○ | ○ | | |
| | S | $-1.98 \times 10^5$ | | ○ | | |

Figure S7: Comparison of the full and partial models in jSDM. (a) Performance of the models. The loglikelihoods of the models (Table 1) are shown. The pairs of full/partial models differing only in the presence/absence of a single explanatory variable are linked with edges, whose thickness represent log-likelihood ratios of the paired models. The OTUs in the input data were defined at the 93%. (b) Models aligned in decreasing order of log-likelihoods. The examined models were roughly classified into three groups in terms of their log-likelihoods.

a

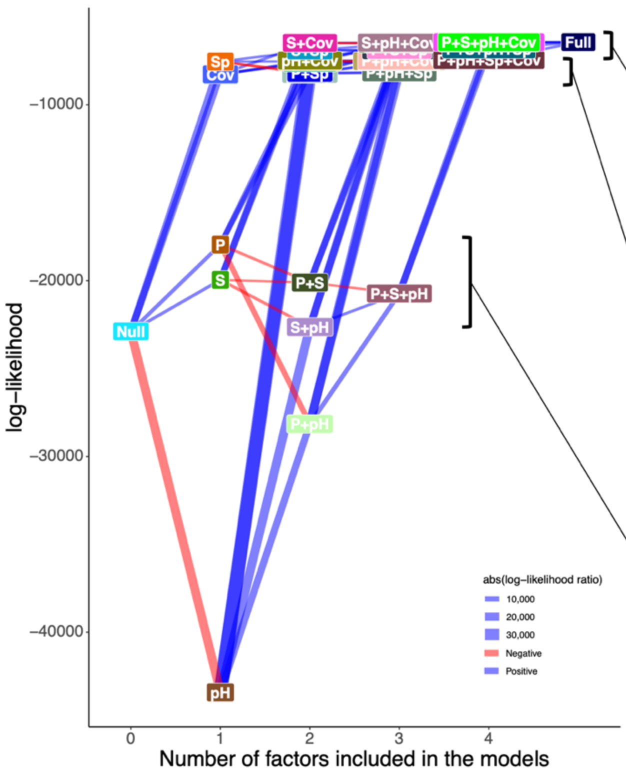


b

| Model name | log-likelihood | Factors in the models: Host plant | Soil fungal community | Soil pH | Spatial autocorrelation | Inter-OTU covariance |
|---|---|---|---|---|---|---|
| Full ( P + S + Sp + Cov) | $-6.43 \times 10^4$ | ○ | ○ | ○ | ○ | ○ |
| P + S + pH + Cov | $-6.44 \times 10^4$ | ○ | ○ | ○ | | ○ |
| S + pH + Sp + Cov | $-6.44 \times 10^4$ | | ○ | ○ | ○ | ○ |
| S + pH + Cov | $-6.46 \times 10^4$ | | ○ | ○ | | ○ |
| P + S + Sp + Cov | $-6.47 \times 10^4$ | ○ | ○ | | ○ | ○ |
| S + Cov | $-6.48 \times 10^4$ | | ○ | | | ○ |
| P + S + Cov | $-6.48 \times 10^4$ | ○ | ○ | | | ○ |
| S + Sp + Cov | $-6.52 \times 10^4$ | | ○ | | ○ | ○ |
| P + S + pH + Sp | $-6.84 \times 10^4$ | ○ | ○ | ○ | ○ | |
| P + S + Sp | $-6.87 \times 10^4$ | ○ | ○ | | ○ | |
| S + pH + Sp | $-6.89 \times 10^4$ | | ○ | ○ | ○ | |
| S + Sp | $-6.91 \times 10^4$ | | ○ | | ○ | |
| P + pH + Sp + Cov | $-7.45 \times 10^4$ | ○ | | ○ | ○ | ○ |
| P + pH + Cov | $-7.48 \times 10^4$ | ○ | | ○ | | ○ |
| pH + Cov | $-7.48 \times 10^4$ | | | ○ | | ○ |
| pH + Sp + Cov | $-7.51 \times 10^4$ | | | ○ | ○ | ○ |
| Sp + Cov | $-7.53 \times 10^4$ | | | | ○ | ○ |
| P + Sp + Cov | $-7.54 \times 10^4$ | ○ | | | ○ | ○ |
| Sp | $-7.54 \times 10^4$ | | | | ○ | |
| P + Cov | $-7.60 \times 10^4$ | ○ | | | | ○ |
| P + pH + Sp | $-8.15 \times 10^4$ | ○ | | ○ | ○ | |
| P + Sp | $-8.17 \times 10^4$ | ○ | | | ○ | |
| pH + Sp | $-8.24 \times 10^4$ | | | ○ | ○ | |
| Cov | $-8.27 \times 10^4$ | | | | | ○ |
| P | $-1.80 \times 10^5$ | ○ | | | | |
| S | $-2.00 \times 10^5$ | | ○ | | | |
| P + S | $-2.01 \times 10^5$ | ○ | ○ | | | |
| P + S + pH | $-2.07 \times 10^5$ | ○ | ○ | ○ | | |
| S + pH | $-2.26 \times 10^5$ | | ○ | ○ | | |
| Null | $-2.29 \times 10^5$ | | | | | |
| P + pH | $-2.82 \times 10^5$ | ○ | | ○ | | |
| pH | $-4.34 \times 10^5$ | | | ○ | | |

Figure S8: Comparison of the full and partial models in jSDM with soil pH. (a) Performance of the models. The loglikelihoods of the models (Table 1) are shown. The pairs of full/partial models differing only in the presence/absence of a single explanatory variable are linked with edges, whose thickness represent log-likelihood ratios of the paired models. The OTUs in the input data were defined at the 97%. (b) Models aligned in decreasing order of log-likelihoods. The examined models were roughly classified into three groups in terms of their log-likelihoods.

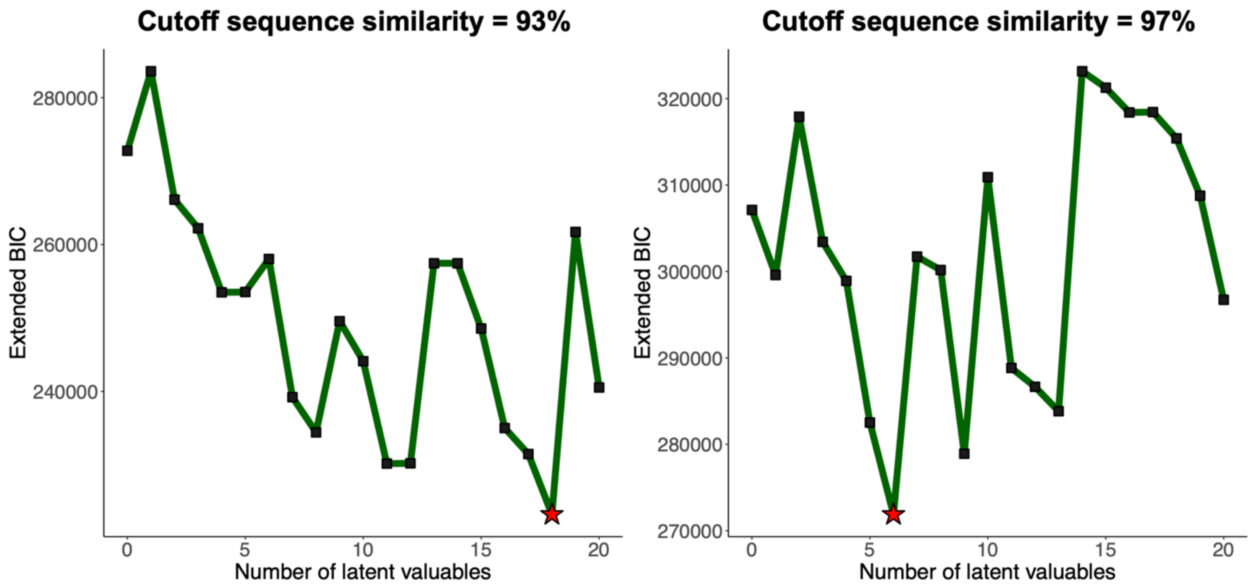


Figure S9: Relationship between the number of assumed latent variables and extended BIC in the co-occurrence network analysis. Co-occurrence network analysis with 0-20 latent variables assumed was conducted for the fungal OTUs included in the analysis in the jSDM analysis. The extended BIC of each estimated network is shown on the vertical axis. The point with the smallest extended BIC is indicated by stars. Results based for the two cutoff sequence similarity thresholds for defining OTUs (93% and 97%) are respectively shown.

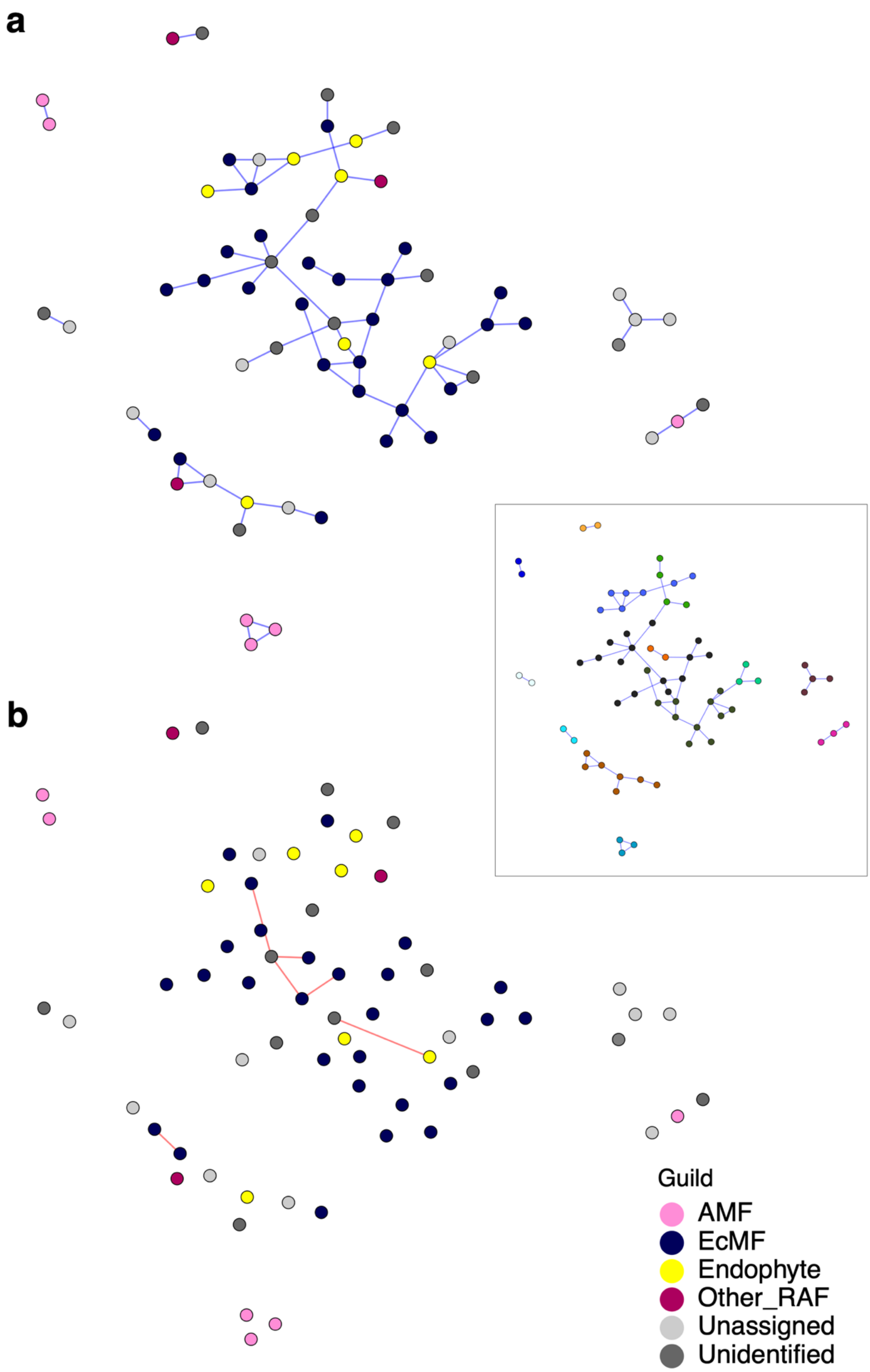
a
b
Guild
AMF
EcMF
Endophyte
Other_RAF
Unassigned
Unidentified

77

Figure S10: Networks of fungus–fungus associations. Potential interactions between fungal OTUs (defined at the 93% cutoff sequence similarity) were inferred based on the SLR method of SPIEC-EASI. Potential effects of environmental preferences shared between fungal OTUs were controlled in the BIC-selected best model, which included six latent variables. The inferred network architecture was shown separately for positive (a) and negative (b) associations between fungal OTUs. For the positive association network (a), modules of densely associated sets of fungal OTUs were inferred with the Louvain algorithm (see the box in the right side).

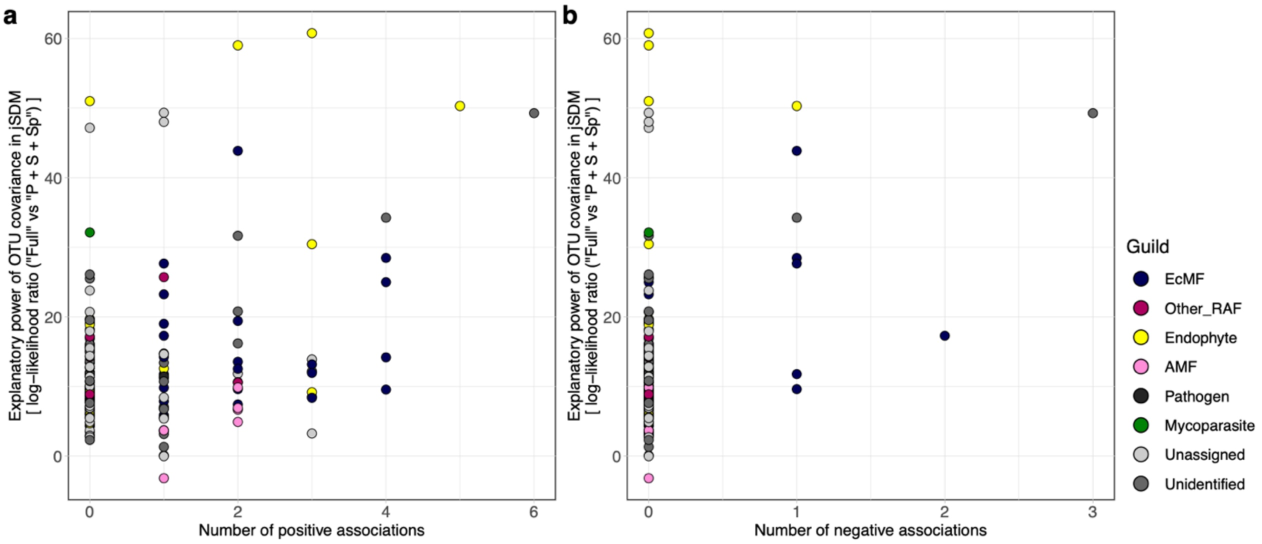


Figure S11: Fungal OTUs (cutoff sequence similarity = 93% similarity) with the strongest signs of species interactions. Fungi potentially playing key roles within the community were explored based on both the jSDM (Figure 2) and SPIEC-EASI (Figure 8) frameworks. Based on the jSDM, the log-likelihood ratios of the two models differing in the presence/absence of fungus–fungus covariance [“Full (P + S + SP + Cov)” vs. “P + S + SP”; see Table 1] are shown for respective fungal OTUs along the vertical axis. Meanwhile, the number of positive (a) or negative (b) associations (i.e., degree centrality) within the SPIEC-EASI networks is shown along the horizontal axis. The degree centrality was significantly correlated with the log-likelihood ratio of the jSDM for both analyses of positive (Kendal’s correlation coefficient = 0.283, $P < 0.001$) and negative (Kendal’s correlation coefficient = 0.256, $P < 0.001$) associations. Colors represent fungal functional guilds.

| Phylum | Class | Order | Family | Genus | Guild (manual) | Reference | Reference DOI |
|---|---|---|---|---|---|---|---|
| Basidiomycota | Agaricomycetes | Cantharellales | Hydnaceae | *Sistotrema* | EcMF | Marino et al., 2008 | https://doi.org/10.1007/s11557-008-0562-4 |
| Ascomycota | Leotiomycetes | Helotiales | Helotiaceae | *Pseudoclathrosphaerina* | Endophyte | Takashima et al., 2014 | https://doi.org/10.11178/jdsa.9.81 |
| Ascomycota | Dothideomycetes | Pleosporales | Pleosporaceae | *Curvularia* | Pathogen | Falloon, 1975 | https://doi.org/10.1080/00288233.1976.10426773 |
| Ascomycota | Eurotiomycetes | Chaetothyriales | Herpotrichiellaceae | *Exophiala* | Endophyte | Li et al., 2011 | https://doi.org/10.1016/j.scitotenv.2010.12.012 |
| Ascomycota | Leotiomycetes | Helotiales | Leptodontidiaceae | *Leptodontidium* | Endophyte | Upson et al., 2009 | https://doi.org/10.1016/j.funeco.2009.02.004 |
| Ascomycota | Sordariomycetes | Diaporthales | Diaporthaceae | *Diaporthe* | Pathogen | Yang et al., 2018 | https://doi.org/10.3897/mycokeys.39.26914 |
| Ascomycota | Sordariomycetes | Hypocreales | Nectriaceae | *Fusarium* | Pathogen | Gordon and Martyn, 1997 | https://doi.org/10.1146/annurev.phyto.35.1.111 |

Table S1: Fungal genera whose lifestyle in plant roots were well characterized in previous studies but different functional guilds were inferred with the automatic assignment based on the Fungaltraits database. The table shows their taxonomic information and manually corrected guild with references.

| | | **Factors in the models** | | | | |
|---|---|---|---|---|---|---|
| | Model name | Host plant | Soil fungal community | Soil pH | Spatial autocorrelation | inter-OTU covariance |
| Model 1 | Full ( P + S + pH + Sp + Cov ) | ○ | ○ | ○ | ○ | ○ |
| Model 2 | P + S + pH + Sp | ○ | ○ | ○ | ○ | |
| Model 3 | P + S + pH + Cov | ○ | ○ | ○ | | ○ |
| Model 4 | P + S + Sp + Cov | ○ | ○ | | ○ | ○ |
| Model 5 | P + pH + Sp + Cov | ○ | | ○ | ○ | ○ |
| Model 6 | S + pH + Sp + Cov | | ○ | ○ | ○ | ○ |
| Model 7 | P + S + pH | ○ | ○ | ○ | | |
| Model 8 | P + S + Sp | ○ | ○ | | ○ | |
| Model 9 | P + S + Cov | ○ | ○ | | | ○ |
| Model 10 | P + pH + Sp | ○ | | ○ | ○ | |
| Model 11 | P + pH + Cov | ○ | | ○ | | ○ |
| Model 12 | P + Sp + Cov | ○ | | | ○ | ○ |
| Model 13 | S + pH + Sp | | ○ | ○ | ○ | |
| Model 14 | S + pH + Cov | | ○ | ○ | | ○ |
| Model 15 | S + Sp + Cov | | ○ | | ○ | ○ |
| Model 16 | pH + Sp + Cov | | | ○ | ○ | ○ |
| Model 17 | P + S | ○ | ○ | | | |
| Model 18 | P + pH | ○ | | ○ | | |
| Model 19 | P + Sp | ○ | | | ○ | |
| Model 20 | P + Cov | ○ | | | | ○ |
| Model 21 | S + pH | | ○ | ○ | | |
| Model 22 | S + Sp | | ○ | | ○ | |
| Model 23 | S + Cov | | ○ | | | ○ |
| Model 24 | pH + Sp | | | ○ | ○ | |
| Model 25 | pH + Cov | | | ○ | | ○ |
| Model 26 | Sp + Cov | | | | ○ | ○ |
| Model 27 | P | ○ | | | | |
| Model 28 | S | | ○ | | | |
| Model 29 | pH | | | ○ | | |
| Model 30 | Sp | | | | ○ | |
| Model 31 | Cov | | | | | ○ |
| Model 32 | Null | | | | | |

Table S2：Models examined in the jSDM framework. The explanatory variables examined in each of the full and partial models are presented. The repertoires of the explanatory variables are host plant species (P), the fungal community structure of background soil (S), soil pH (pH), spatial autocorrelation (Sp), and fungus–fungus covariance (co-occurrence) patterns (Cov).

| OTU | Log-likelihood ratio | Guild | Scientific Name | Query Cover | E value | Per. Ident | Accession |
|---|---|---|---|---|---|---|---|
| X_07951 | 24.3 | Unidentified | Venturiales sp. | 100% | 1.00E-113 | 98.77% | MW471960.1 |
| X_06373 | 19.1 | AMF | *Rhizophagus* sp. | 100% | 7.00E-70 | 93.72% | LC544104.1 |
| X_00149 | 17.6 | Unassigned | Hyaloscyphaceae sp. | 100% | 1.00E-112 | 97.96% | MT587170.1 |
| X_00682 | 17.5 | Unassigned | *Hyaloscypha* cf. *hepaticicola* | 100% | 6.00E-104 | 95.95% | AY579413.1 |
| X_00128 | 15.7 | Other_RAF | *Archaeorhizomyces borealis* | 100% | 2.00E-96 | 100.00% | NR_126144.2 |
| X_00966 | 13.8 | AMF | *Rhizophagus* sp. | 100% | 6.00E-77 | 96.32% | LC544104.1 |
| X_09467 | 13.8 | Other_RAF | *Archaeorhizomyces* sp. 'victor nom. seq.' | 100% | 8.00E-88 | 97.98% | OQ676542.1 |
| X_01277 | 13.4 | Endophyte | *Pezicula ericae* | 100% | 1.00E-119 | 100.00% | MT294405.1 |
| X_06962 | 13.1 | Mycoparasite | fungal sp. 25 GM 18-01 | 100% | 2.00E-96 | 95.74% | KF359663.1 |
| X_00511 | 12.6 | AMF | *Rhizophagus* sp. | 100% | 3.00E-67 | 93.16% | LC544104.1 |
| X_00361 | 11.2 | Unidentified | *Hyaloscypha bicolor* | 100% | 3.00E-108 | 96.75% | KX611541.1 |
| X_01506 | 11.2 | AMF | *Rhizophagus* sp. | 100% | 1.00E-72 | 94.24% | LC544104.1 |
| X_00301 | 11.2 | Unassigned | Mortierellales sp. | 100% | 3.00E-113 | 99.58% | MN385503.1 |
| X_00235 | 11.1 | Endophyte | *Cladophialophora* sp. | 100% | 4.00E-118 | 99.59% | MK537116.1 |
| X_03905 | 10.2 | AMF | Glomeraceae sp. | 100% | 2.00E-77 | 96.81% | MT765311.1 |
| X_02808 | 9.71 | Unassigned | *Hyaloscypha bicolor* | 100% | 6.00E-110 | 97.14% | KX611541.1 |
| X_00467 | 9.67 | Endophyte | *Cladophialophora* sp. | 100% | 3.00E-113 | 98.38% | OM744986.1 |
| X_00176 | 9.35 | Endophyte | *Phialocephala fortinii* | 100% | 1.00E-119 | 100.00% | MT294419.1 |
| X_00249 | 7.56 | EcMF | *Russula* sp. | 100% | 2.00E-117 | 99.59% | OQ421803.1 |
| X_01179 | 7.48 | Endophyte | Helotiaceae sp. II GK-2010 | 93% | 6.00E-104 | 97.82% | HQ157864.1 |

Table S3: Effects of host plant identity on fungal OTUs' distribution. For each fungal OTU (cutoff sequence similarity = 97%), impacts of host plants on root-associated fungal distributions was inferred by the log-likelihood ratio between the models with and without the explanatory variable ("P + S + Sp" vs. "S + Sp"; Table 1). The top-20 OTUs with the greatest log-likelihood ratios are shown in the list. For these OTUs, the log-likelihood ratio scores are shown with the OTU IDs, the functional guilds automatically assigned by the Fungaltraits database, and BLAST top-hit results [the scientific name of the matched sequence, query cover, E-value, percent of identity ("Per. Ident"), and NCBI accession number].

| OTU | Log-likelihood ratio | Guild | Scientific Name | Query Cover | E value | Per. Ident | Accession |
|---|---|---|---|---|---|---|---|
| X_00084 | $1.80 \times 10^{2}$ | Unidentified | *Membranomyces* cf. delectabilis GB0090523 | 80% | 4.00E-93 | 99.49% | JQ638714.1 |
| X_00683 | $1.66 \times 10^{2}$ | Unassigned | *Psathyrella suavissima* | 100% | 4.00E-100 | 94.35% | KC992899.1 |
| X_00120 | $1.64 \times 10^{2}$ | EcMF | *Russula* aff. adusta aff. 1 ST-2020 | 100% | 5.00E-98 | 94.29% | MW024879.1 |
| X_00218 | $1.49 \times 10^{2}$ | EcMF | *Russula* sp. | 100% | 1.00E-119 | 100.00% | KU924614.1 |
| X_00249 | $1.47 \times 10^{2}$ | EcMF | *Russula* sp. | 100% | 2.00E-117 | 99.59% | OQ421803.1 |
| X_00789 | $1.46 \times 10^{2}$ | EcMF | *Tomentella* sp. | 98% | 4.00E-100 | 95.90% | KY686241.1 |
| X_00369 | $1.34 \times 10^{2}$ | EcMF | *Tricholoma* sp. 22 XXD-2021 | 99% | 3.00E-107 | 97.12% | MW724366.1 |
| X_00265 | $1.32 \times 10^{2}$ | EcMF | *Russula velenovskyi* | 100% | 1.00E-119 | 100.00% | AY061721.1 |
| X_00361 | $1.29 \times 10^{2}$ | Unidentified | *Hyaloscypha bicolor* | 100% | 3.00E-108 | 96.75% | KX611541.1 |
| X_00625 | $1.27 \times 10^{2}$ | Unassigned | *Leptodontidium* sp. | 100% | 3.00E-113 | 98.37% | OM745610.1 |
| X_00686 | $1.26 \times 10^{2}$ | Unassigned | *Mortierella* sp. | 100% | 6.00E-110 | 100.00% | ON075228.1 |
| X_00436 | $1.26 \times 10^{2}$ | EcMF | *Russula* sp. | 93% | 4.00E-106 | 98.70% | OP133164.1 |
| X_00807 | $1.24 \times 10^{2}$ | EcMF | *Tomentella terrestris* | 100% | 1.00E-113 | 98.39% | MW472220.1 |
| X_00282 | $1.20 \times 10^{2}$ | EcMF | *Russula* sp. HMAS 276805 | 100% | 1.00E-100 | 94.40% | LT602953.1 |
| X_09467 | $1.20 \times 10^{2}$ | Other_RAF | *Archaeorhizomyces* sp. 'victor nom. seq.' | 100% | 8.00E-88 | 97.98% | OQ676542.1 |
| X_01102 | $1.11 \times 10^{2}$ | EcMF | *Amanita orientifulva* | 97% | 5.00E-117 | 100.00% | LC098745.1 |
| X_18065 | $1.08 \times 10^{2}$ | Unassigned | *Geoglossum simile* | 100% | 4.00E-105 | 98.68% | KF854293.1 |
| X_00289 | $1.05 \times 10^{2}$ | EcMF | Thelephoraceae sp. H217 | 100% | 3.00E-94 | 92.65% | AB634273.1 |
| X_00732 | $1.03 \times 10^{2}$ | EcMF | *Russula* sp. | 100% | 3.00E-114 | 98.78% | OQ322556.1 |
| X_00452 | $1.00 \times 10^{2}$ | Endophyte | *Leptodontidium* sp. | 100% | 1.00E-119 | 100.00% | OM745579.1 |

Table S4: Effects of background soil fungal community structure on fungal OTUs' distribution. For each fungal OTU (cutoff sequence similarity = 97%), impacts of background soil community structure on root-associated fungal distributions was inferred by the log-likelihood ratio between the models with and without the explanatory variable ("P + S + Sp" vs. "P + Sp"; Table 1). The top-20 OTUs with the greatest log-likelihood ratios are shown in the list. For these OTUs, the log-likelihood ratio scores are shown with the OTU IDs, the functional guilds automatically assigned by the Fungaltraits database, and BLAST top-hit results [the scientific name of the matched sequence, query cover, E-value, percent of identity ("Per. Ident"), and NCBI accession number].

| OTU | Log-likelihood ratio | Guild | Scientific Name | Query Cover | E value | Per. Ident | Accession |
|---|---|---|---|---|---|---|---|
| X_01617 | $1.26 \times 10^3$ | EcMF | *Amphinema* sp. 2 YC-2021 | 100% | 1.00E-119 | 100.00% | MW546513.1 |
| X_01750 | $1.24 \times 10^3$ | Unidentified | fungal sp. 15 GM 14-03 | 100% | 2.00E-65 | 90.45% | KF359653.1 |
| X_00741 | $1.21 \times 10^3$ | EcMF | *Laccaria* sp. 'PA01' | 100% | 4.00E-118 | 99.59% | OP749048.1 |
| X_01704 | $1.21 \times 10^3$ | Unassigned | *Ramariopsis* sp. | 100% | 5.00E-111 | 97.56% | MT644620.1 |
| X_02651 | $1.20 \times 10^3$ | Unidentified | Ascomycota sp. | 100% | 3.00E-114 | 98.78% | OM744898.1 |
| X_04225 | $1.20 \times 10^3$ | Unidentified | *Ramicandelaber taiwanensis* | 25% | 5.00E-16 | 95.16% | NR_165200.1 |
| X_01329 | $1.20 \times 10^3$ | Unidentified | *Clavulinopsis luteoalba* | 100% | 2.00E-79 | 90.28% | OP538704.1 |
| X_11552 | $1.19 \times 10^3$ | Unidentified | *Glomus macrocarpum* | 100% | 3.00E-80 | 97.33% | LC544108.1 |
| X_01327 | $1.19 \times 10^3$ | Unidentified | fungal sp. | 100% | 1.00E-99 | 94.29% | MW471758.1 |
| X_00739 | $1.19 \times 10^3$ | EcMF | *Naucoria bohemica* | 100% | 3.00E-114 | 98.78% | MW243069.1 |
| X_00883 | $1.18 \times 10^3$ | Unassigned | *Galerina* sp. | 100% | 1.00E-93 | 92.24% | OK161020.1 |
| X_01353 | $1.18 \times 10^3$ | EcMF | *Tomentella* sp. 280 VM-2015 | 100% | 5.00E-117 | 99.18% | KP783478.1 |
| X_02352 | $1.18 \times 10^3$ | Unassigned | *Ramariopsis avellaneo-inversa* | 100% | 1.00E-112 | 98.37% | MT055954.1 |
| X_01875 | $1.18 \times 10^3$ | EcMF | *Sebacina* sp. | 96% | 2.00E-84 | 90.79% | OQ410802.1 |
| X_00880 | $1.17 \times 10^3$ | EcMF | *Russula burlinghamiae* | 88% | 4.00E-99 | 98.61% | MZ157246.1 |
| X_01821 | $1.16 \times 10^3$ | Unidentified | Helotiales sp. | 100% | 5.00E-111 | 98.33% | OM745587.1 |
| X_00030 | $1.15 \times 10^3$ | EcMF | *Tarzetta catinus* | 100% | 9.00E-114 | 100.00% | LC619231.1 |
| X_04800 | $1.15 \times 10^3$ | Unidentified | Mortierellales sp. RB-2011 | 100% | 4.00E-99 | 95.18% | JQ272348.1 |
| X_00289 | $1.15 \times 10^3$ | EcMF | Thelephoraceae sp. H217 | 100% | 3.00E-94 | 92.65% | AB634273.1 |
| X_00718 | $1.14 \times 10^3$ | Unidentified | Sordariomycetes sp. SM13-17-5-3 | 100% | 3.00E-39 | 98.06% | MN905840.1 |

Table S5: Effects of spatial autocorrelation on fungal OTUs' distribution. For each fungal OTU (cutoff sequence similarity = 97%), impacts of spatial autocorrelation on root-associated fungal distributions was inferred by the log-likelihood ratio between the models with and without the explanatory variable ("P + S + Sp" vs. "P + S"; Table 1). The top-20 OTUs with the greatest log-likelihood ratios are shown in the list. For these OTUs, the log-likelihood ratio scores are shown with the OTU IDs, the functional guilds automatically assigned by the Fungaltraits database, and BLAST top-hit results [the scientific name of the matched sequence, query cover, E-value, percent of identity ("Per. Ident"), and NCBI accession number].

| OTU | Log-likelihood ratio | Guild | Scientific Name | Query Cover | E value | Per. Ident | Accession |
|---|---|---|---|---|---|---|---|
| X_00176 | 50.8 | Endophyte | *Phialocephala fortinii* | 100% | 1.00E-119 | 100.00% | MT294419.1 |
| X_00452 | 45.4 | Endophyte | *Leptodontidium* sp. | 100% | 1.00E-119 | 100.00% | OM745579.1 |
| X_00149 | 44.9 | Unassigned | Hyaloscyphaceae sp. | 100% | 1.00E-112 | 97.96% | MT587170.1 |
| X_00235 | 43.4 | Endophyte | *Cladophialophora* sp. | 100% | 4.00E-118 | 99.59% | MK537116.1 |
| X_00409 | 42.0 | Endophyte | *Oidiodendron maius* | 100% | 1.00E-119 | 100.00% | LC206669.1 |
| X_00054 | 41.2 | Unassigned | *Mortierella* sp. SM13-9-10-3 | 100% | 7.00E-115 | 100.00% | MN905910.1 |
| X_00301 | 40.1 | Unassigned | Mortierellales sp. | 100% | 3.00E-113 | 99.58% | MN385503.1 |
| X_00282 | 30.8 | EcMF | *Russula* sp. HMAS 276805 | 100% | 1.00E-100 | 94.40% | LT602953.1 |
| X_06962 | 30.4 | Mycoparasite | fungal sp. 25 GM 18-01 | 100% | 2.00E-96 | 95.74% | KF359663.1 |
| X_00084 | 26.7 | Unidentified | *Membranomyces* cf. delectabilis GB0090523 | 80% | 4.00E-93 | 99.49% | JQ638714.1 |
| X_00389 | 26.2 | Endophyte | Hyaloscyphaceae sp. CC 14-35 | 100% | 6.00E-110 | 97.58% | KF359568.1 |
| X_00467 | 26.2 | Endophyte | *Cladophialophora* sp. | 100% | 3.00E-113 | 98.38% | OM744986.1 |
| X_00286 | 24.8 | Other_RAF | *Mycena pradensis* | 100% | 1.00E-119 | 100.00% | MT196392.1 |
| X_00361 | 24.1 | Unidentified | *Hyaloscypha bicolor* | 100% | 3.00E-108 | 96.75% | KX611541.1 |
| X_00900 | 23.1 | Unidentified | Herpotrichiellaceae sp. | 100% | 1.00E-119 | 100.00% | OK584496.1 |
| X_00436 | 22.1 | EcMF | *Russula* sp. | 93% | 4.00E-106 | 98.70% | OP133164.1 |
| X_01961 | 21.5 | Endophyte | *Cladophialophora chaetospira* | 100% | 1.00E-119 | 100.00% | HQ871875.1 |
| X_00249 | 21.4 | EcMF | *Russula* sp. | 100% | 2.00E-117 | 99.59% | OQ421803.1 |
| X_00682 | 20.9 | Unassigned | *Hyaloscypha* cf. hepaticicola | 100% | 6.00E-104 | 95.95% | AY579413.1 |
| X_01621 | 20.9 | Unassigned | *Mortierella* sp. | 100% | 6.00E-110 | 100.00% | MN385496.1 |

Table S6: Effects of fungus–fungus covariance on fungal OTUs' distribution. For each fungal OTU (cutoff sequence similarity = 97%), impacts of fungus–fungus covariance on root-associated fungal distributions was inferred by the log-likelihood ratio between the models with and without the explanatory variable ("Full (P + S + Sp + Cov)" vs. "P + S + Sp"; Table 1). The top-20 OTUs with the greatest log-likelihood ratios are shown in the list. For these OTUs, the log-likelihood ratio scores are shown with the OTU IDs, the functional guilds automatically assigned by the Fungaltraits database, and BLAST top-hit results [the scientific name of the matched sequence, query cover, E-value, percent of identity ("Per. Ident"), and NCBI accession number].

| OTU | z-standardized *d'* (fungi) | *P* (FDR) | Guild | Scientific Name | Query Cover | E value | Per. Ident | Accession |
|---|---|---|---|---|---|---|---|---|
| X_00966 | 7.60 | $P < 0.001$ | AMF | *Rhizophagus sp.* | 100% | 6.00E-77 | 96.32% | LC544104.1 |
| X_00511 | 7.12 | $P < 0.001$ | AMF | *Rhizophagus sp.* | 100% | 4.00E-67 | 93.16% | LC544104.1 |
| X_06373 | 6.60 | $P < 0.001$ | AMF | *Rhizophagus sp.* | 100% | 7.00E-70 | 93.72% | LC544104.1 |
| X_01506 | 6.56 | $P < 0.001$ | AMF | *Rhizophagus sp.* | 100% | 1.00E-72 | 94.24% | LC544104.1 |
| X_03905 | 4.19 | $P < 0.01$ | AMF | Glomeraceae sp. | 100% | 2.00E-77 | 96.81% | MT765311.1 |
| X_02153 | 4.02 | $P < 0.01$ | AMF | Glomeraceae sp. | 100% | 5.00E-78 | 96.32% | MT765456.1 |
| X_04205 | 3.10 | $P < 0.05$ | AMF | Glomeraceae sp. | 100% | 4.00E-79 | 95.85% | MT765456.1 |
| X_00700 | 2.84 | $P < 0.05$ | EcMF | *Russula emetica* | 100% | 1.00E-119 | 100.00% | KX579814.1 |
| X_01277 | 6.82 | $P < 0.001$ | Endophyte | *Pezicula ericae* | 100% | 1.00E-119 | 100.00% | MT294405.1 |
| X_00176 | 5.46 | $P < 0.001$ | Endophyte | *Phialocephala fortinii* | 100% | 1.00E-119 | 100.00% | MT294419.1 |
| X_01814 | 3.47 | $P < 0.05$ | Endophyte | *Oidiodendron* sp. | 100% | 8.00E-109 | 96.73% | OP720876.1 |
| X_00235 | 3.18 | $P < 0.05$ | Endophyte | *Cladophialophora* sp. | 100% | 5.00E-118 | 99.59% | MK537116.1 |
| X_01179 | 2.96 | $P < 0.05$ | Endophyte | Helotiaceae sp. II GK-2010 | 93% | 6.00E-104 | 97.82% | HQ157864.1 |
| X_06962 | 4.73 | $P < 0.01$ | Mycoparasite | fungal sp. 25 GM 18-01 | 100% | 2.00E-96 | 95.74% | KF359663.1 |
| X_09467 | 5.34 | $P < 0.001$ | Other_RAF | *Archaeorhizomyces* sp. 'victor nom. seq.' | 100% | 9.00E-88 | 97.98% | OQ676542.1 |
| X_00128 | 3.73 | $P < 0.01$ | Other_RAF | *Archaeorhizomyces borealis* | 100% | 2.00E-96 | 100.00% | NR_126144.2 |
| X_00682 | 5.06 | $P < 0.001$ | Unassigned | *Hyaloscypha* cf. hepaticicola | 100% | 6.00E-104 | 95.95% | AY579413.1 |
| X_00301 | 4.19 | $P < 0.05$ | Unassigned | *Podila humilis* | 100% | 4.00E-113 | 99.58% | JF439486.1 |
| X_01405 | 4.05 | $P < 0.01$ | Unassigned | *Dissophora globulifera* | 100% | 5.00E-118 | 99.59% | MW393876.1 |
| X_01294 | 4.00 | $P < 0.01$ | Unassigned | *Hyaloscypha daedaleae* | 100% | 1.00E-107 | 97.14% | MF161321.1 |
| X_09083 | 3.96 | $P < 0.01$ | Unassigned | *Chalara angustata* | 83% | 7.00E-97 | 100.00% | NR_159786.1 |
| X_00149 | 3.35 | $P < 0.05$ | Unassigned | Hyaloscyphaceae sp. | 100% | 1.00E-112 | 97.96% | MT587170.1 |
| X_02314 | 3.33 | $P < 0.05$ | Unassigned | *Leptodophora orchidicola* | 100% | 1.00E-119 | 100.00% | MT294412.1 |
| X_00956 | 3.08 | $P < 0.05$ | Unassigned | *Clavulinopsis* sp. 'laeticolor-IN03' | 100% | 3.00E-114 | 98.78% | ON650114.1 |
| X_02808 | 2.95 | $P < 0.05$ | Unassigned | *Hyaloscypha bicolor* | 100% | 7.00E-110 | 97.14% | KX611541.1 |
| X_02874 | 2.82 | $P < 0.05$ | Unassigned | Thelephoraceae sp. | 96% | 1.00E-106 | 97.46% | OQ410922.1 |
| X_00503 | 2.76 | $P < 0.05$ | Unassigned | *Leucoscypha* sp. | 82% | 4.00E-87 | 97.03% | OM672930.1 |
| X_07951 | 8.15 | $P < 0.001$ | Unidentified | Venturiales sp. | 100% | 1.00E-113 | 98.77% | MW471960.1 |
| X_11552 | 5.70 | $P < 0.001$ | Unidentified | *Glomus macrocarpum* | 100% | 3.00E-80 | 97.33% | LC544108.1 |
| X_04519 | 3.30 | $P < 0.05$ | Unidentified | Ascomycota sp. | 85% | 4.00E-62 | 89.76% | MN898690.1 |
| X_01585 | 3.27 | $P < 0.05$ | Unidentified | *Densocarpa shanorii* | 22% | 2.00E-16 | 100.00% | KT361850.1 |

Table S7: Host preference of fungal OTUs. With randomization test of *d'* metric, fungal OTUs (cutoff sequence similarity = 97%) with significant host preference were screened. Fungal OTUs showing significant host preference patterns are shown with preference metrics (*z*-standardized *d'* ), FDR-corrected *P*-values, and BLAST top-hit results [the scientific name of the matched sequence, query cover, E-value, percent of identity ("Per. Ident"), and NCBI accession number].

| OTU | Host plant | *2DP* (z-standardized) | *P* (FDR) | Scientific Name | Query Cover | E value | Per. Ident | Accession |
|---|---|---|---|---|---|---|---|---|
| X_01277 | *Acer* | 5.49 | $P < 0.001$ | *Pezicula ericae* | 100% | 1.00E-119 | 100.00% | NR_155653.1 |
| X_06373 | *Acer* | 4.81 | $P < 0.001$ | *Rhizophagus* sp. | 100% | 7.00E-70 | 93.72% | LC544104.1 |
| X_06962 | *Acer* | 4.16 | $P < 0.001$ | fungal sp. 25 GM 18-01 | 100% | 2.00E-96 | 95.74% | KF359663.1 |
| X_00511 | *Acer* | 4.08 | $P < 0.001$ | *Rhizophagus* sp. | 100% | 3.00E-67 | 93.16% | LC544104.1 |
| X_09083 | *Acer* | 4.07 | $P < 0.001$ | *Chalara angustata* | 83% | 6.00E-97 | 100.00% | NR_159786.1 |
| X_07951 | *Acer* | -3.75 | $P < 0.001$ | Venturiales sp. | 100% | 1.00E-113 | 98.77% | MW471960.1 |
| X_00682 | *Betula* | 4.33 | $P < 0.001$ | *Hyaloscypha* cf. hepaticicola | 100% | 6.00E-104 | 95.95% | AY579413.1 |
| X_03905 | *Juglans* | 4.16 | $P < 0.001$ | Glomeraceae sp. | 100% | 2.00E-77 | 96.81% | MT765311.1 |
| X_04205 | *Juglans* | 3.90 | $P < 0.001$ | Glomeraceae sp. | 100% | 4.00E-79 | 95.85% | MT765456.1 |
| X_07951 | *Pinus* | 5.40 | $P < 0.001$ | Venturiales sp. | 100% | 1.00E-113 | 98.77% | MW471960.1 |
| X_09467 | *Pinus* | 4.61 | $P < 0.001$ | *Archaeorhizomyces* sp. 'victor nom. seq.' | 100% | 9.00E-88 | 97.98% | OQ676542.1 |
| X_02874 | *Pinus* | 4.26 | $P < 0.01$ | Thelephoraceae sp. | 96% | 1.00E-106 | 97.46% | OQ410922.1 |
| X_00128 | *Pinus* | 3.93 | $P < 0.001$ | *Archaeorhizomyces borealis* | 100% | 2.00E-96 | 100.00% | NR_126144.2 |
| X_01334 | *Pinus* | 3.78 | $P < 0.01$ | *Amphinema* sp. 7 UK-2011 | 100% | 1.00E-119 | 100.00% | JN943925.1 |
| X_06962 | *Pinus* | -3.38 | $P < 0.05$ | fungal sp. 25 GM 18-01 | 100% | 2.00E-96 | 95.74% | KF359663.1 |
| X_06373 | *Pinus* | -3.82 | $P < 0.001$ | *Rhizophagus* sp. | 100% | 7.00E-70 | 93.72% | LC544104.1 |
| X_00682 | *Pinus* | -3.85 | $P < 0.001$ | *Hyaloscypha* cf. hepaticicola | 100% | 6.00E-104 | 95.95% | AY579413.1 |
| X_01506 | *Pinus* | -3.90 | $P < 0.001$ | *Rhizophagus* sp. | 100% | 1.00E-72 | 94.24% | LC544104.1 |
| X_00511 | *Pinus* | -4.00 | $P < 0.001$ | *Rhizophagus* sp. | 100% | 3.00E-67 | 93.16% | LC544104.1 |
| X_00966 | *Pinus* | -4.21 | $P < 0.001$ | *Rhizophagus* sp. | 100% | 6.00E-77 | 96.32% | LC544104.1 |
| X_00149 | *Pinus* | -4.35 | $P < 0.001$ | Hyaloscyphaceae sp. | 100% | 1.00E-112 | 97.96% | MT587170.1 |
| X_00807 | *Populus* | 3.99 | $P < 0.01$ | *Tomentella terrestris* | 100% | 1.00E-113 | 98.39% | MW472220.1 |
| X_00807 | *Pyrus* | -4.28 | $P < 0.001$ | *Tomentella terrestris* | 100% | 1.00E-113 | 98.39% | MW472220.1 |
| X_00966 | *Toxicodendron* | 3.65 | $P < 0.001$ | *Rhizophagus* sp. | 100% | 6.00E-77 | 96.32% | LC544104.1 |

Table S8: Randomization analysis of plant–fungal associations. Based on a randomization test of two-dimensional preference (*2DP*), pairs of plants and fungal OTUs (cutoff sequence similarity = 97%) showing significant signs of preference/avoidance are screened. The plant–fungus pairs listed are shown with preference metrics (*z*-standardized *2DP*), FDR-corrected *P*-values, and BLAST top-hit results of the fungal OTUs [the scientific name of the matched sequence, query cover, E-value, percent of identity ("Per. Ident"), and NCBI accession number].

| OTU | Regression coefficent | *P* (FDR) | Guild | Scientific Name | Query Cover | E value | Per. Ident | Accession |
|---|---|---|---|---|---|---|---|---|
| X_06373 | 0.181 | $P < 0.001$ | AMF | *Rhizophagus* sp. | 100% | 7.00E-70 | 93.72% | LC544104.1 |
| X_01102 | 0.553 | $P < 0.001$ | EcMF | *Amanita orientifulva* | 97% | 5.00E-117 | 100.00% | LC098745.1 |
| X_00369 | 0.549 | $P < 0.001$ | EcMF | *Tricholoma* sp. 22 XXD-2021 | 99% | 3.00E-107 | 97.12% | MW724366.1 |
| X_01875 | 0.533 | $P < 0.01$ | EcMF | *Sebacina* sp. | 96% | 2.00E-84 | 90.79% | OQ410802.1 |
| X_01082 | 0.467 | $P < 0.001$ | EcMF | *Sistotrema flavorhizomorphae* | 100% | 1.00E-119 | 100.00% | NR_178118.1 |
| X_00807 | 0.462 | $P < 0.001$ | EcMF | *Tomentella terrestris* | 100% | 1.00E-113 | 98.39% | MW472220.1 |
| X_02343 | 0.452 | $P < 0.001$ | EcMF | *Cortinarius ferrugineovelatus* | 100% | 4.00E-118 | 99.59% | MH930161.1 |
| X_01023 | 0.421 | $P < 0.001$ | EcMF | *Sebacina* sp. | 96% | 8.00E-115 | 100.00% | OQ410872.1 |
| X_00880 | 0.415 | $P < 0.001$ | EcMF | *Russula burlinghamiae* | 88% | 5.00E-99 | 98.61% | MZ157246.1 |
| X_00693 | 0.414 | $P < 0.001$ | EcMF | *Tomentella* sp. T1228 | 100% | 4.00E-118 | 99.59% | AB634259.1 |
| X_00218 | 0.398 | $P < 0.001$ | EcMF | *Russula vesca* | 100% | 1.00E-119 | 100.00% | KX094999.1 |
| X_00766 | 0.384 | $P < 0.001$ | EcMF | *Tuber* sp. | 83% | 1.00E-94 | 99.02% | LC556138.1 |
| X_00700 | 0.357 | $P < 0.001$ | EcMF | *Russula emetica* | 100% | 1.00E-119 | 100.00% | KX579814.1 |
| X_01551 | 0.349 | $P < 0.001$ | EcMF | *Tomentella* sp. | 100% | 1.00E-112 | 97.96% | OQ418543.1 |
| X_00436 | 0.340 | $P < 0.001$ | EcMF | *Russula* sp. | 93% | 4.00E-106 | 98.70% | OP133164.1 |
| X_00337 | 0.332 | $P < 0.001$ | EcMF | *Lactarius vinaceorufescens* | 99% | 4.00E-112 | 98.35% | OP749847.1 |
| X_01649 | 0.315 | $P < 0.001$ | EcMF | *Inocybe suaveolens* | 100% | 5.00E-117 | 99.18% | HQ604205.1 |
| X_01134 | 0.311 | $P < 0.001$ | EcMF | *Clavulina amethystina* | 100% | 3.00E-107 | 96.33% | MN959776.1 |
| X_00440 | 0.303 | $P < 0.001$ | EcMF | *Russula turci* | 100% | 1.00E-119 | 100.00% | KF002780.1 |
| X_00888 | 0.301 | $P < 0.001$ | EcMF | *Tomentella subtestacea* | 100% | 8.00E-109 | 96.73% | MW546519.1 |
| X_00249 | 0.295 | $P < 0.001$ | EcMF | *Russula* sp. | 100% | 2.00E-117 | 99.59% | OQ421803.1 |
| X_01210 | 0.294 | $P < 0.001$ | EcMF | *Pachyphlodes* sp. | 100% | 5.00E-118 | 99.59% | OM672915.1 |
| X_00800 | 0.288 | $P < 0.01$ | EcMF | *Tomentella sublilacina* | 100% | 1.00E-119 | 100.00% | AJ889976.1 |
| X_00030 | 0.285 | $P < 0.001$ | EcMF | *Tarzetta catinus* | 100% | 9.00E-114 | 100.00% | LC619231.1 |
| X_00120 | 0.267 | $P < 0.001$ | EcMF | *Russula* aff. adusta aff. 1 ST-2020 | 100% | 5.00E-98 | 94.29% | MW024879.1 |
| X_00789 | 0.257 | $P < 0.001$ | EcMF | *Tomentella* sp. | 98% | 4.00E-100 | 95.90% | KY686241.1 |
| X_02666 | 0.247 | $P < 0.01$ | EcMF | *Russula* sp. HMAS 276805 | 100% | 1.00E-105 | 96.34% | LT602953.1 |
| X_01361 | 0.245 | $P < 0.05$ | EcMF | *Laccaria bicolor* | 100% | 6.00E-117 | 99.18% | JX504116.1 |
| X_01617 | 0.236 | $P < 0.001$ | EcMF | *Amphinema* sp. 2 YC-2021 | 100% | 1.00E-119 | 100.00% | MW546513.1 |
| X_00282 | 0.236 | $P < 0.001$ | EcMF | *Russula* sp. HMAS 276805 | 100% | 1.00E-100 | 94.40% | LT602953.1 |
| X_00226 | 0.133 | $P < 0.01$ | EcMF | *Lactarius tabidus* | 100% | 1.00E-119 | 100.00% | KX095062.1 |
| X_01825 | 0.488 | $P < 0.001$ | Endophyte | Helotiaceae sp. 3 GM 13-01 | 100% | 1.00E-112 | 97.96% | KF359565.1 |
| X_00862 | 0.434 | $P < 0.001$ | Endophyte | Helotiaceae sp. 3 GM 13-01 | 100% | 8.00E-96 | 93.47% | KF359565.1 |
| X_00723 | 0.228 | $P < 0.001$ | Endophyte | *Oidiodendron* sp. | 100% | 1.00E-119 | 100.00% | OM745603.1 |
| X_01179 | 0.197 | $P < 0.05$ | Endophyte | Helotiaceae sp. II GK-2010 | 93% | 6.00E-104 | 97.82% | HQ157864.1 |
| X_00235 | 0.180 | $P < 0.001$ | Endophyte | *Cladophialophora chaetospira* | 100% | 4.00E-118 | 99.59% | KF359558.1 |
| X_00389 | 0.167 | $P < 0.01$ | Endophyte | Hyaloscyphaceae sp. CC 14-35 | 100% | 7.00E-110 | 97.58% | KF359568.1 |
| X_00452 | 0.106 | $P < 0.01$ | Endophyte | *Leptodontidium* sp. | 100% | 1.00E-119 | 100.00% | OM745579.1 |
| X_09467 | 0.290 | $P < 0.001$ | Other_RAF | *Archaeorhizomyces* sp. 'victor nom. seq.' | 100% | 9.00E-88 | 97.98% | OQ676542.1 |
| X_01025 | 0.276 | $P < 0.05$ | Other_RAF | *Mortierella* sp. SM13-27-18-1 | 100% | 7.00E-109 | 100.00% | MN905904.1 |
| X_00286 | 0.239 | $P < 0.001$ | Other_RAF | *Mycena pradensis* | 100% | 1.00E-119 | 100.00% | MT196392.1 |
| X_00128 | 0.157 | $P < 0.01$ | Other_RAF | *Archaeorhizomyces borealis* | 100% | 2.00E-96 | 100.00% | NR_126144.2 |
| X_00860 | 0.574 | $P < 0.001$ | Unassigned | *Hyaloscypha variabilis* | 100% | 4.00E-118 | 99.59% | MN947404.1 |
| X_00956 | 0.537 | $P < 0.001$ | Unassigned | *Clavulinopsis* sp. 'laeticolor-IN03' | 100% | 3.00E-114 | 98.78% | ON650114.1 |
| X_03252 | 0.494 | $P < 0.001$ | Unassigned | *Ramariopsis* sp. | 99% | 4.00E-112 | 97.96% | OP896794.1 |
| X_01704 | 0.470 | $P < 0.001$ | Unassigned | *Ramariopsis* sp. | 100% | 5.00E-111 | 97.56% | MT644620.1 |
| X_03788 | 0.407 | $P < 0.001$ | Unassigned | fungal sp. | 100% | 7.00E-109 | 97.14% | MF965332.1 |
| X_01204 | 0.391 | $P < 0.001$ | Unassigned | *Clavulinopsis* sp. 'laeticolor-IN01' | 100% | 2.00E-115 | 98.78% | OR168885.1 |
| X_02874 | 0.333 | $P < 0.001$ | Unassigned | Thelephoraceae sp. | 96% | 1.00E-106 | 97.46% | OQ410922.1 |
| X_01217 | 0.321 | $P < 0.001$ | Unassigned | Clavariaceae sp. | 100% | 2.00E-104 | 95.51% | MT587230.1 |
| X_01356 | 0.304 | $P < 0.001$ | Unassigned | *Penicillium montanense* | 100% | 1.00E-119 | 100.00% | MT514387.1 |
| X_02352 | 0.295 | $P < 0.001$ | Unassigned | *Ramariopsis avellaneo-inversa* | 100% | 1.00E-112 | 98.37% | MT055954.1 |
| X_18065 | 0.294 | $P < 0.001$ | Unassigned | *Geoglossum simile* | 100% | 5.00E-105 | 98.68% | KF854293.1 |
| X_03539 | 0.270 | $P < 0.001$ | Unassigned | *Odontia parvispina* | 100% | 5.00E-92 | 90.51% | MW033328.1 |
| X_01807 | 0.259 | $P < 0.05$ | Unassigned | *Entoloma* aff. rhodocylix | 100% | 5.00E-118 | 99.59% | ON943248.1 |
| X_00625 | 0.246 | $P < 0.001$ | Unassigned | *Leptodontidium* sp. | 100% | 3.00E-113 | 98.37% | OM745610.1 |

| | | | | | | | | |
|---|---|---|---|---|---|---|---|---|
| X_01294 | 0.235 | $P < 0.001$ | Unassigned | *Hyaloscypha daedaleae* | 100% | 1.00E-107 | 97.14% | MF161321.1 |
| X_00682 | 0.139 | $P < 0.01$ | Unassigned | *Hyaloscypha* cf. hepaticicola | 100% | 6.00E-104 | 95.95% | AY579413.1 |
| X_00519 | 0.479 | $P < 0.001$ | Unidentified | *Aestipascuomyces dupliciliberans* | 29% | 1.00E-16 | 98.28% | MW049145.1 |
| X_01750 | 0.414 | $P < 0.001$ | Unidentified | fungal sp. 15 GM 14-03 | 100% | 2.00E-65 | 90.45% | KF359653.1 |
| X_00084 | 0.404 | $P < 0.001$ | Unidentified | *Membranomyces* cf. delectabilis GB0090523 | 80% | 4.00E-93 | 99.49% | JQ638714.1 |
| X_01254 | 0.389 | $P < 0.001$ | Unidentified | *Hyaloscypha* sp. ECRU075 | 100% | 5.00E-117 | 99.18% | KM678388.1 |
| X_02284 | 0.381 | $P < 0.001$ | Unidentified | Basidiomycota sp. CC 01-08 | 84% | 2.00E-28 | 74.01% | KF359631.1 |
| X_01106 | 0.336 | $P < 0.05$ | Unidentified | Mortierellales sp. GD22a | 100% | 7.00E-109 | 99.56% | EF152530.1 |
| X_02651 | 0.319 | $P < 0.001$ | Unidentified | Ascomycota sp. | 100% | 3.00E-114 | 98.78% | OM744898.1 |
| X_00166 | 0.315 | $P < 0.001$ | Unidentified | Cryptomycota sp. | 97% | 3.00E-31 | 76.78% | MT353563.1 |
| X_01819 | 0.295 | $P < 0.001$ | Unidentified | fungal sp. | 81% | 3.00E-57 | 86.26% | MG190403.1 |
| X_00718 | 0.290 | $P < 0.001$ | Unidentified | Sordariomycetes sp. SM13-17-5-3 | 100% | 3.00E-39 | 98.06% | MN905840.1 |
| X_01585 | 0.289 | $P < 0.001$ | Unidentified | *Densocarpa shanorii* | 22% | 1.00E-16 | 100.00% | KT361850.1 |
| X_01976 | 0.279 | $P < 0.001$ | Unidentified | *Cladophialophora* sp. | 100% | 5.00E-117 | 99.59% | MK536958.1 |
| X_00361 | 0.266 | $P < 0.001$ | Unidentified | *Hyaloscypha bicolor* | 100% | 3.00E-108 | 96.75% | KX611541.1 |
| X_17682 | 0.246 | $P < 0.01$ | Unidentified | fungal sp. | 82% | 1.00E-36 | 76.47% | MK564906.1 |
| X_01332 | 0.224 | $P < 0.05$ | Unidentified | *Russula carmesina* | 100% | 2.00E-91 | 92.68% | MT925405.1 |
| X_00211 | 0.212 | $P < 0.01$ | Unidentified | Ascomycota sp. RT-2012 | 100% | 4.00E-35 | 79.60% | JQ711841.1 |
| X_00645 | 0.204 | $P < 0.01$ | Unidentified | *Rhizophagus clarus* | 100% | 1.00E-47 | 84.92% | FN423697.1 |
| X_01413 | 0.141 | $P < 0.05$ | Unidentified | *Rosettozyma petaloides* | 100% | 9.00E-39 | 75.61% | NR_174037.1 |

Table S9: Relationship between fungal OTU occurrence in root and relative abundance in the soil. Fungal OTUs (cutoff sequence similarity = 97%) with significant relationship between occurrence in root samples and relative abundance in soil samples are listed. Fungal OTUs with significant standardized partial regression coefficients in the generalized liner model (GLM) examined in Figure 6 are shown with FDR-corrected *P*-values, inferred guilds (functional groups), and BLAST top-hit results [the scientific name of the matched sequence, query cover, E-value, percent of identity ("Per. Ident"), and NCBI accession number].

| OTU | Module | Guild | Scientific Name | Query Cover | E value | Per. Ident | Accession |
|---|---|---|---|---|---|---|---|
| X_02005 | 1 | EcMF | *Russula nigricans* | 99% | 2.00E-117 | 99.59% | MW172333.1 |
| X_00739 | 1 | EcMF | *Naucoria bohemica* | 100% | 3.00E-114 | 98.78% | MW243069.1 |
| X_00369 | 1 | EcMF | *Tricholoma* sp. 22 XXD-2021 | 99% | 3.00E-107 | 97.12% | MW724366.1 |
| X_00289 | 1 | EcMF | Thelephoraceae sp. H217 | 100% | 4.00E-94 | 92.65% | AB634273.1 |
| X_00030 | 1 | EcMF | *Tarzetta catinus* | 100% | 1.00E-113 | 100.00% | LC619231.1 |
| X_02874 | 1 | Unassigned | Thelephoraceae sp. | 96% | 1.00E-106 | 97.46% | OQ410922.1 |
| X_02808 | 1 | Unassigned | *Hyaloscypha bicolor* | 100% | 7.00E-110 | 97.14% | KX611541.1 |
| X_01334 | 1 | Unidentified | *Tylospora* sp. | 100% | 1.00E-119 | 100.00% | OR482711.1 |
| X_00361 | 1 | Unidentified | *Hyaloscypha bicolor* | 100% | 3.00E-108 | 96.75% | KX611541.1 |
| X_00084 | 1 | Unidentified | *Membranomyces* cf. delectabilis GB0090523 | 80% | 4.00E-93 | 99.49% | JQ638714.1 |
| X_00218 | 2 | EcMF | *Russula vesca* | 100% | 1.00E-119 | 100.00% | KX094999.1 |
| X_01316 | 2 | Other_RAF | Mortierellales sp. | 93% | 5.00E-111 | 100.00% | MT533763.1 |
| X_00301 | 2 | Unassigned | *Podila humilis* | 100% | 4.00E-113 | 99.58% | JF439486.1 |
| X_00300 | 2 | Unassigned | *Saitozyma podzolica* | 100% | 7.00E-89 | 100.00% | KY320605.1 |
| X_00054 | 2 | Unassigned | *Podila minutissima* | 100% | 8.00E-115 | 100.00% | MK513846.1 |
| X_00807 | 3 | EcMF | *Tomentella terrestris* | 100% | 1.00E-113 | 98.39% | MW472220.1 |
| X_00789 | 3 | EcMF | *Tomentella* sp. | 98% | 4.00E-100 | 95.90% | KY686241.1 |
| X_00693 | 3 | EcMF | *Tomentella* sp. T1228 | 100% | 5.00E-118 | 99.59% | AB634259.1 |
| X_00440 | 3 | EcMF | *Russula turci* | 100% | 1.00E-119 | 100.00% | KF002780.1 |
| X_00265 | 3 | EcMF | *Russula velenovskyi* | 100% | 1.00E-119 | 100.00% | AY061721.1 |
| X_00120 | 3 | EcMF | *Russula* aff. adusta aff. 1 ST-2020 | 100% | 6.00E-98 | 94.29% | MW024879.1 |
| X_00235 | 3 | Endophyte | *Cladophialophora* sp. | 100% | 5.00E-118 | 99.59% | MK537116.1 |
| X_00682 | 3 | Unassigned | *Hyaloscypha* cf. hepaticicola | 100% | 6.00E-104 | 95.95% | AY579413.1 |
| X_00128 | 4 | Other_RAF | *Archaeorhizomyces borealis* | 100% | 2.00E-96 | 100.00% | NR_126144.2 |
| X_04645 | 4 | Unassigned | *Pezoloma* sp. | 100% | 1.00E-106 | 96.73% | OM951709.1 |
| X_01413 | 4 | Unidentified | *Rosettozyma petaloides* | 100% | 9.00E-39 | 75.61% | NR_174037.1 |
| X_00337 | 5 | EcMF | *Lactarius cremicolor* | 100% | 1.00E-119 | 100.00% | OR364573.1 |
| X_01277 | 5 | Endophyte | *Pezicula ericae* | 100% | 1.00E-119 | 100.00% | MT294405.1 |
| X_00452 | 5 | Endophyte | *Leptodontidium* sp. | 100% | 1.00E-119 | 100.00% | OM745579.1 |
| X_00176 | 5 | Endophyte | *Phialocephala fortinii* | 100% | 1.00E-119 | 100.00% | MT294419.1 |
| X_06962 | 5 | Mycoparasite | fungal sp. 25 GM 18-01 | 100% | 2.00E-96 | 95.74% | KF359663.1 |
| X_00286 | 5 | Other_RAF | *Mycena pradensis* | 100% | 1.00E-119 | 100.00% | MT196392.1 |
| X_01204 | 5 | Unassigned | *Clavulinopsis* sp. 'laeticolor-IN01' | 100% | 2.00E-115 | 98.78% | OR168885.1 |
| X_00625 | 5 | Unassigned | *Leptodontidium* sp. | 100% | 4.00E-113 | 98.37% | OM745610.1 |
| X_00149 | 5 | Unassigned | Hyaloscyphaceae sp. | 100% | 1.00E-112 | 97.96% | MT587170.1 |
| X_01254 | 5 | Unidentified | *Hyaloscypha* sp. ECRU075 | 100% | 6.00E-117 | 99.18% | KM678388.1 |
| X_00900 | 5 | Unidentified | Herpotrichiellaceae sp. | 100% | 1.00E-119 | 100.00% | OK584496.1 |
| X_00211 | 5 | Unidentified | Ascomycota sp. RT-2012 | 100% | 4.00E-35 | 79.60% | JQ711841.1 |
| X_00732 | 6 | EcMF | *Russula* sp. | 100% | 3.00E-114 | 98.78% | OQ322556.1 |
| X_01825 | 6 | Endophyte | Helotiaceae sp. 3 GM 13-01 | 100% | 1.00E-112 | 97.96% | KF359565.1 |
| X_00862 | 6 | Endophyte | Helotiaceae sp. 3 GM 13-01 | 100% | 9.00E-96 | 93.47% | KF359565.1 |
| X_09467 | 6 | Other_RAF | *Archaeorhizomyces* sp. 'victor nom. seq.' | 100% | 9.00E-88 | 97.98% | OQ676542.1 |
| X_00860 | 6 | Unassigned | *Hyaloscypha variabilis* | 100% | 5.00E-118 | 99.59% | MN947404.1 |
| X_00166 | 6 | Unidentified | Cryptomycota sp. | 97% | 3.00E-31 | 76.78% | MT353563.1 |
| X_00700 | 7 | EcMF | *Russula emetica* | 100% | 1.00E-119 | 100.00% | KX579814.1 |
| X_00226 | 7 | EcMF | *Lactarius tabidus* | 100% | 1.00E-119 | 100.00% | KX095062.1 |
| X_00389 | 7 | Endophyte | Hyaloscyphaceae sp. CC 14-35 | 100% | 7.00E-110 | 97.58% | KF359568.1 |
| X_03607 | 7 | Pathogen | *Fusarium oxysporum* | 100% | 5.00E-111 | 100.00% | MF281350.2 |
| X_00686 | 7 | Unassigned | *Mortierella* sp. | 100% | 6.00E-110 | 100.00% | ON075228.1 |
| X_00683 | 7 | Unassigned | *Psathyrella suavissima* | 100% | 4.00E-100 | 94.35% | KC992899.1 |
| X_00505 | 7 | Unassigned | *Mortierella* sp. S-24 | 100% | 5.00E-104 | 99.10% | KJ735018.1 |
| X_00503 | 7 | Unassigned | *Leucoscypha* sp. | 82% | 4.00E-87 | 97.03% | OM672930.1 |
| X_01649 | 8 | EcMF | *Inocybe umbratica* | 100% | 6.00E-117 | 99.18% | OR364571.1 |
| X_01082 | 8 | EcMF | *Sistotrema flavorhizomorphae* | 100% | 1.00E-119 | 100.00% | NR_178118.1 |
| X_00249 | 8 | EcMF | *Russula* sp. | 100% | 2.00E-117 | 99.59% | OQ421803.1 |
| X_01961 | 8 | Endophyte | *Cladophialophora chaetospira* | 100% | 1.00E-119 | 100.00% | HQ871875.1 |
| X_00467 | 8 | Endophyte | *Cladophialophora* sp. L3-3-NN-2016 | 100% | 4.00E-113 | 98.38% | MN148682.1 |
| X_00409 | 8 | Endophyte | *Oidiodendron maius* | 100% | 1.00E-119 | 100.00% | LC206669.1 |
| X_07951 | 8 | Unidentified | Venturiales sp. | 100% | 1.00E-113 | 98.77% | MW471960.1 |
| X_01976 | 8 | Unidentified | *Cladophialophora* sp. | 100% | 6.00E-117 | 99.59% | MK536958.1 |

| | | | | | | | |
|---|---|---|---|---|---|---|---|
| X_02666 | 9 | EcMF | *Russula* sp. HMAS 276805 | 100% | 1.00E-105 | 96.34% | LT602953.1 |
| X_00436 | 9 | EcMF | *Russula* sp. | 93% | 4.00E-106 | 98.70% | OP133164.1 |
| X_00282 | 9 | EcMF | *Russula* sp. HMAS 276805 | 100% | 1.00E-100 | 94.40% | LT602953.1 |
| X_00723 | 9 | Endophyte | *Oidiodendron* sp. | 100% | 1.00E-119 | 100.00% | OM745603.1 |
| X_06373 | 10 | AMF | *Rhizophagus* sp. | 100% | 7.00E-70 | 93.72% | LC544104.1 |
| X_01506 | 10 | AMF | *Rhizophagus* sp. | 100% | 1.00E-72 | 94.24% | LC544104.1 |
| X_00966 | 10 | AMF | *Rhizophagus* sp. | 100% | 6.00E-77 | 96.32% | LC544104.1 |
| X_00511 | 10 | AMF | *Rhizophagus* sp. | 100% | 4.00E-67 | 93.16% | LC544104.1 |
| X_18065 | 10 | Unassigned | *Geoglossum simile* | 100% | 5.00E-105 | 98.68% | KF854293.1 |
| X_02620 | 11 | Unidentified | *Rhizophagus* sp. | 100% | 2.00E-45 | 83.58% | MK418525.1 |
| X_00645 | 11 | Unidentified | *Rhizophagus clarus* | 100% | 1.00E-47 | 84.92% | FN423697.1 |
| X_03403 | 12 | EcMF | *Genea hispidula* | 100% | 2.00E-117 | 99.59% | MT505214.1 |
| X_02343 | 12 | EcMF | *Cortinarius ferrugineovelatus* | 100% | 5.00E-118 | 99.59% | MH930161.1 |
| X_01134 | 12 | EcMF | *Clavulina amethystina* | 100% | 3.00E-107 | 96.33% | MN959776.1 |
| X_00894 | 12 | EcMF | *Russula cascadensis* | 100% | 1.00E-112 | 97.96% | KF359616.1 |
| X_04205 | 13 | AMF | Glomeraceae sp. | 100% | 4.00E-79 | 95.85% | MT765456.1 |
| X_03905 | 13 | AMF | Glomeraceae sp. | 100% | 2.00E-77 | 96.81% | MT765311.1 |
| X_02153 | 13 | AMF | Glomeraceae sp. | 100% | 5.00E-78 | 96.32% | MT765456.1 |

Table S10: Modules of densely associated sets of fungal OTUs. The modules of the fungal OTUs (cutoff sequence similarity = 97 %) in the positive co-occurrence network (Figure 8a) were inferred with the Louvain algorithm. Fungal OTUs are shown with their network modules, inferred guilds (functional groups), and BLAST top-hit results [the scientific name of the matched sequence, query cover, E-value, percent of identity (“Per. Ident”), and NCBI accession number].

| OTU | Number of Co-occurrence | Guild | Scientific Name | Query Cover | E value | Per. Ident | Accession |
|---|---|---|---|---|---|---|---|
| X_00361 | 7 | Unidentified | *Hyaloscypha bicolor* | 100% | 3.00E-108 | 96.75% | KX611541.1 |
| X_00249 | 6 | EcMF | *Russula* sp. | 100% | 2.00E-117 | 99.59% | OQ421803.1 |
| X_00054 | 5 | Unassigned | *Mortierella* sp. SM13-9-10-3 | 100% | 7.00E-115 | 100.00% | MN905910.1 |
| X_00149 | 5 | Unassigned | Hyaloscyphaceae sp. | 100% | 1.00E-112 | 97.96% | MT587170.1 |
| X_00226 | 5 | EcMF | *Lactarius tabidus* | 100% | 1.00E-119 | 100.00% | MH862189.1 |
| X_00084 | 4 | Unidentified | *Membranomyces* cf. delectabilis GB0090523 | 80% | 4.00E-93 | 99.49% | JQ638714.1 |
| X_00235 | 4 | Endophyte | *Cladophialophora chaetospira* | 100% | 5.00E-118 | 99.59% | KF359558.1 |
| X_00265 | 4 | EcMF | *Russula velenovskyi* | 100% | 1.00E-119 | 100.00% | AY061721.1 |
| X_00282 | 4 | EcMF | *Russula* sp. HMAS 276805 | 100% | 1.00E-100 | 94.40% | LT602953.1 |
| X_00452 | 4 | Endophyte | *Leptodontidium* sp. | 100% | 1.00E-119 | 100.00% | OM745579.1 |
| X_00467 | 4 | Endophyte | *Cladophialophora* sp. L3-3-NN-2016 | 100% | 4.00E-113 | 98.38% | MN148682.1 |
| X_00625 | 4 | Unassigned | *Leptodontidium* sp. | 100% | 4.00E-113 | 98.37% | OM745610.1 |
| X_00683 | 4 | Unassigned | *Psathyrella suavissima* | 100% | 4.00E-100 | 94.35% | KC992899.1 |
| X_00686 | 4 | Unassigned | *Mortierella* sp. | 100% | 6.00E-110 | 100.00% | ON075228.1 |
| X_06373 | 4 | AMF | *Rhizophagus* sp. | 100% | 7.00E-70 | 93.72% | LC544104.1 |
| X_09467 | 4 | Other_RAF | *Archaeorhizomyces* sp. 'victor nom. seq.' | 100% | 9.00E-88 | 97.98% | OQ676542.1 |
| X_00389 | 3 | Endophyte | Hyaloscyphaceae sp. CC 14-35 | 100% | 7.00E-110 | 97.58% | KF359568.1 |
| X_00511 | 3 | AMF | *Rhizophagus* sp. | 100% | 4.00E-67 | 93.16% | LC544104.1 |
| X_00693 | 3 | EcMF | *Tomentella* sp. T1228 | 100% | 5.00E-118 | 99.59% | AB634259.1 |
| X_00732 | 3 | EcMF | *Russula* sp. | 100% | 3.00E-114 | 98.78% | OQ322556.1 |

Table S11: Degree centrality of fungal OTUs (cutoff sequence similarity = 97%) in the positive fungus–fungus association network. With the SLR method of SPIEC-EASI analysis, fungus–fungus associations were inferred by controlling the potential effects of shared environmental preference (Figure S10a,b). Top-20 fungal OTUs with the highest degree centrality scores (i.e., the number of network links) within the positive association network are shown with the information of inferred guilds (functional groups) and BLAST top-hit results [the scientific name of the matched sequence, query cover, E-value, percent of identity ("Per. Ident"), and NCBI accession number].

| OTU | Number of associations | Guild | Scientific Name | Query Cover | E value | Per. Ident | Accession |
|---|---|---|---|---|---|---|---|
| X_00176 | 6 | Endophyte | *Phialocephala fortinii* | 100% | 1.00E-119 | 100.00% | MT294419.1 |
| X_00084 | 4 | Unidentified | *Membranomyces* cf. delectabilis GB0090523 | 80% | 4.00E-93 | 99.49% | JQ638714.1 |
| X_00452 | 3 | Endophyte | *Leptodontidium* sp. | 100% | 1.00E-119 | 100.00% | OM745579.1 |
| X_00120 | 2 | EcMF | *Russula* aff. adusta aff. 1 ST-2020 | 100% | 6.00E-98 | 94.29% | MW024879.1 |
| X_00149 | 2 | Unassigned | Hyaloscyphaceae sp. | 100% | 1.00E-112 | 97.96% | MT587170.1 |
| X_00218 | 2 | EcMF | *Russula vesca* | 100% | 1.00E-119 | 100.00% | KX094999.1 |
| X_00226 | 2 | EcMF | *Lactarius tabidus* | 100% | 1.00E-119 | 100.00% | KX095062.1 |
| X_00282 | 2 | EcMF | *Russula* sp. HMAS 276805 | 100% | 1.00E-100 | 94.40% | LT602953.1 |
| X_00361 | 2 | Unidentified | *Hyaloscypha bicolor* | 100% | 3.00E-108 | 96.75% | KX611541.1 |
| X_00030 | 1 | EcMF | *Tarzetta catinus* | 100% | 1.00E-113 | 100.00% | LC619231.1 |
| X_00235 | 1 | Endophyte | *Cladophialophora* sp. | 100% | 5.00E-118 | 99.59% | MK537116.1 |
| X_00265 | 1 | EcMF | *Russula velenovskyi* | 100% | 1.00E-119 | 100.00% | AY061721.1 |
| X_00301 | 1 | Unassigned | *Podila humilis* | 100% | 4.00E-113 | 99.58% | JF439486.1 |
| X_00389 | 1 | Endophyte | Hyaloscyphaceae sp. CC 14-35 | 100% | 7.00E-110 | 97.58% | KF359568.1 |
| X_00436 | 1 | EcMF | *Russula* sp. | 93% | 4.00E-106 | 98.70% | OP133164.1 |
| X_00618 | 1 | Unassigned | fungal sp. | 100% | 5.00E-118 | 99.59% | MH624582.1 |
| X_00625 | 1 | Unassigned | *Leptodontidium* sp. | 100% | 4.00E-113 | 98.37% | OM745610.1 |
| X_00683 | 1 | Unassigned | *Psathyrella suavissima* | 100% | 4.00E-100 | 94.35% | KC992899.1 |
| X_01023 | 1 | EcMF | *Sebacina* sp. | 96% | 9.00E-115 | 100.00% | OQ410872.1 |
| X_01334 | 1 | Unidentified | *Tylospora* sp. | 100% | 1.00E-119 | 100.00% | OR482711.1 |

Table S12: Degree centrality of the fungal OTUs (cutoff sequence similarity = 97%) in the negative fungus–fungus association network. With the SLR method of SPIEC-EASI analysis, fungus–fungus associations were inferred by controlling the potential effects of shared environmental preference (Figure S10a,b). Top-20 fungal OTUs with the highest degree centrality scores (i.e., the number of network links) within the negative association network are shown with the information of inferred guilds (functional groups) and BLAST top-hit results [the scientific name of the matched sequence, query cover, E-value, percent of identity ("Per. Ident"), and NCBI accession number].
